# A time-resolved meta-analysis of consensus gene expression profiles during human T-cell activation

**DOI:** 10.1101/2023.05.03.538418

**Authors:** Michael Rade, Sebastian Böhlen, Vanessa Neuhaus, Dennis Löffler, Conny Blumert, Ulrike Köhl, Susann Dehmel, Katherina Sewald, Kristin Reiche

**Author notes:** Shared last authorship (Katherina Sewald, Kristin Reiche).

## Abstract

**Background:** The coordinated transcriptional regulation of activated T-cells is based on the complex dynamic behavior of signaling networks. Given an external stimulus, T-cell gene expression is characterized by impulse and sustained patterns over the course. Here, we analyzed the temporal pattern of activation across different T-cell populations to develop consensus gene signatures for T-cell activation.

**Methods:** We applied a meta-analysis of anti-CD3/CD28 induced CD4+ T-cell activation kinetics of publicly available transcriptome-wide time series using a random effects model. We used non-negative matrix factorization, an unsupervised deconvolution method, to infer changes in biological patterns over time. For verification and to further map a wider variety of the T-cell landscape, we performed a time series of transcriptome-wide RNA sequencing on activated blood T-cells. Lastly, we matched the identified consensus biomarker signatures to single-cell RNA sequencing (scRNA-Seq) data of autologous anti-CD19 chimeric antigen receptor (CAR) T-cells from 24 patients with large B cell lymphoma (LBCL) to characterize activation status of the cell product before infusion.

**Results:** We identified time-resolved gene expression profiles comprising 521 genes of up to 10 disjunct time points during activation and different polarization conditions. The gene signatures include central transcriptional regulators of T-cell activation, representing successive waves as well as sustained patterns of induction. They cover early, intermediate, and late response expression rates across multiple T-cell populations, thus defining consensus biomarker signatures for T-cell activation. Intermediate and late response activation signatures in CAR T-cell infusion products were correlated to immune effector cell-associated neurotoxicity syndrome.

**Conclusion:** In conclusion, we describe temporally resolved gene expression patterns across T-cell populations. These biomarker signatures are a valuable source for e.g., monitoring transcriptional changes during T-cell activation with a reasonable number of genes, annotating T-cell states in single-cell transcriptome studies or assessing dysregulated functions of human T-cell immunity.

## Introduction

T-cells are a subgroup of lymphocytes and mediate the adaptive cellular immune response. T-cell priming and initiation of activation to an foreign antigen requires engagement of T-cell receptors (TCR) with the cognate peptide-MHC complex presented by antigen-presenting cells (APCs), inducing of costimulatory signals such as CD28 and OX40, and cytokines released by APCs that drive T-cell differentiation (1, 2). These processes initiate coordinated transcriptional changes of genes. T-cells exit the quiescent state by down-regulation of genes associated with maintenance of resting status (3–5) followed by metabolic re-programming to provide energy for growth. This includes, for example, increased mitochondrial biogenesis, glycolysis, or decreased oxidative phosphorylation, which is coupled with the tricarboxylic acid (TCA) cycle. Furthermore, T-cells respond to autocrine or paracrine Interleukin 2 (IL2) signaling and initiate effector functions, such as cytokine production (3, 6). Another essential aspect is post-transcriptional regulation defining T-cell rewiring and maintaining T-cell quiescence. Post-transcriptional regulation determines protein production by regulating RNA splicing, mRNA stability, RNA localization, and translation machinery (7, 8). All these processes must be coordinated in a temporal fashion.

In the past years’ time series of transcriptome-wide studies (microarray, bulk RNA-sequencing (RNA-Seq) and single-cell RNA-Seq) assessed activation and differentiation of T-cells into distinct T-cell populations from healthy donors (9– 18). With such studies, enormous progress has been made in defining molecular mechanisms during T-cell activation and differentiation processes. However, no transcriptome study with a time-resolved description of the transient expression changes from TCR stimulation via signal pathways to proliferation across T-cell populations is reported. But gene signatures robustly monitoring T-cell activation across different T-cell populations, if developed as biomarker gene signatures, i.e. as “measurable and evaluable indicators for the underlying” (19) T-cell activation processes, would enable robust monitoring of immune responses.

Applications of such a biomarker signature are wide-spread. For example, there is a need for human-based in vitro models that allow objective assessment of T-cell dependent immune responses during non-clinical development and safety assessment of new candidates for immunotherapies. Consensus biomarker signatures for T-cell activation accompanying non-clinical safety models enable risk prediction of immune-related adverse events.

Further, advances in manufacturing processes and increasing patient numbers suitable for cellular immunotherapies like autologous CAR T-cell therapy reinforce the importance of robust biomarker signatures predicting toxicity and/or efficacy of CAR T-cells (20). Response rates of CAR T-cell therapies strongly depend on cellular fitness of CAR T-cells in the infusion product. Biomarker signatures as consensus gene expression profiles characterizing temporal cellular changes throughout T-cell activation could function as T-cell intrinsic descriptors correlating with efficacy and safety of CAR T-cells.

Lastly, single cell or spatial transcriptomics often result in sparse datasets. Hence, quantifying expression is often only possible for highly expressed genes because low capture efficiency and abundance of dropout events lead to zero expression measurements. This limits annotation and interpretation of such datasets. Annotation of T-cell states would benefit from an extended list of markers also covering longitudinal expression changes.

We aim at biomarker signatures as consensus gene expression profiles characterizing temporal cellular changes throughout T-cell activation. In this study, we describe a novel and unbiased time-resolved set of genes whose expression patterns correlate strongly with T-cell stimulation processes. We performed a meta-analysis of T-cell activation using time-series transcriptome datasets to develop kinetic consensus gene signatures across multiple T-cell populations. Our results characterize gene expression changes in a coordinated and temporal fashion and provide general biomarker signatures for human T-cell activation.

## Results

### Towards time-resolved consensus gene expression signatures for T-cell activation

To identify a time-resolved consensus gene expression profiles across T-cell populations, we followed a three-step approach. Firstly, we used a *Discovery Set* of publicly available transcriptome data. We aimed at identifying genes with a significant combined effect size among T-cell populations and studies in at least one time point of T-cell activation. T-cells were activated by stimulation with anti-CD3/anti-CD28 coated beads or in the presence of stimulation beads and differentiating cytokines (see Additional file 1: Figure S1). Secondly, we combined genes into classes of coherent metagenes by unsupervised decomposition. Thirdly, we confirmed the time-resolved consensus signatures in 2 independent *Verification Sets* (Fig. 1).

**Fig. 1.**
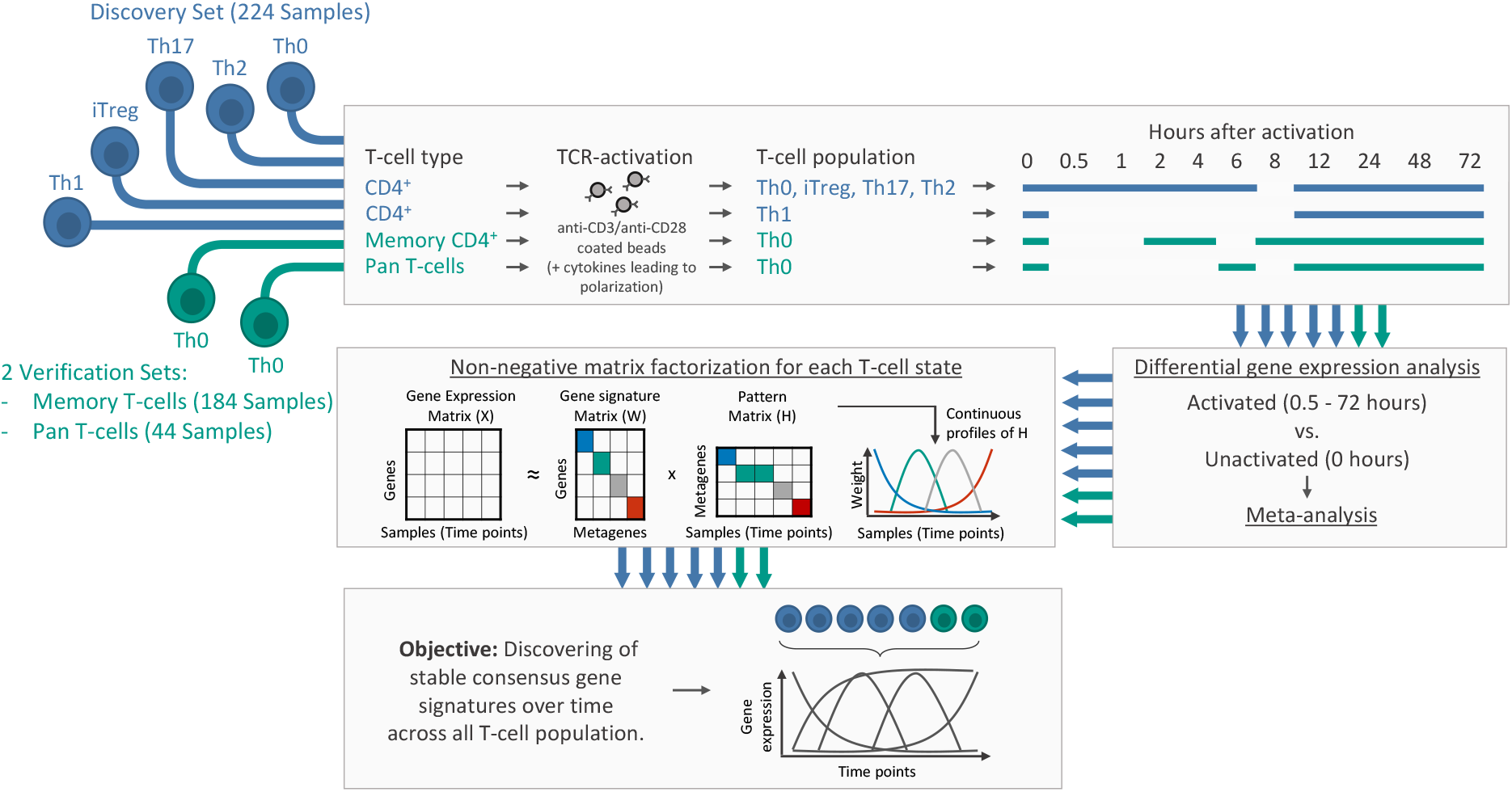
Workflow for discovering and verifying temporal consensus gene expression signatures of T-cells after in-vitro activation with anti-CD3/anti-CD28 coated beads (Th0) or in the presence of stimulation beads and differentiating cytokines (polarization towards Th1, Th2, Th17, and iTreg T-cell fates). For each T-cell population we performed DGEA to find DE (FDR <0.05) genes of activated T-cell populations at different analysis time points compared to unactivated T-cell populations (time series of gene expression arrays and RNA-seq data). To find DE genes with a significant combined effect size (FDR <0.05) across CD4+ T-cell populations from the *Discovery Set* (highlighted in blue), we conducted a meta-analysis. Only DE genes with a significant combined effect size in at least one contrast (0.5 to 72 hours vs. 0 hours) across the available populations (4 populations for time course 0.5 to 6 hours of activation, 5 populations for time course 12 to 72 hours of activation) were used for NMF. We conducted NMF to infer biological patterns over time and to discover stable continuous metagenes (i.e., sets genes with similar expression pattern across the analysis time points) across all T-cell populations. For verification of the temporal consensus gene signature, we analyzed 2 independent RNA-Seq datasets (highlighted in green). The *Discovery- and Memory T-cell Verification Set* are based on publicly available datasets.

The *Discovery Set* included transcriptome-wide expression data of in-vitro kinetic models of CD4+ T-cells, previously profiled by publicly available time series microarrays (10, 11) or RNA-Seq (12–14) studies. In these studies, CD4+ T-cells were obtained from healthy donors consisting of either adult PBMCs or neonatal cord blood. The CD4+ T-cells were collected between 0 (before activation) and 72 hours after activation with anti-CD3/anti-CD28 coated beads only (Th0) or in the presence of differentiating cytokines, leading to polarization towards Th1, Th2, Th17, and iTreg T-cell fates. The datasets of Th0 and iTreg consist of multiple studies and sample sources (PBMCs and cord blood). After removing potential batch effects across studies, we observed no significant correlation between study or sample source and principal components (see Additional file 1: Figure S4). Except for Th1, 9 overlapping time points of activation were available in the T-cell populations (Fig. 1). Overall, we used 224 samples with up to 12 replicates per time point of activation and CD4+ T-cell population in the *Discovery Set*.

For verification, we re-analyzed an independent RNA-Seq dataset (15) (referred to as *Memory T-cell Verification Set*) that included unactivated as well as with anti-CD3/CD28 beads activated memory CD4+ T-cells from 24 healthy donors between 2 and 72 hours (overall 184 samples).

In addition, we isolated Pan T-cells from 4 healthy donors (for MACS isolation strategy of Pan T-cells, see Additional file 1). RNA-Seq was performed at 5 different time points of anti-CD3/anti-CD28 activation (6, 12, 24, 48 and 72 hours) and from unactivated (0 hours) Pan T-cells. We additionally cultured and sequenced Pan T-cells as negative controls in serum-free medium without anti-CD3/anti-CD28 activation at the same time points as mentioned above. We refer to this dataset as *Pan T-cell Verifications Set*. Overall, 44 samples were analyzed (Fig. 1).

For a detailed overview of all datasets used for this study, see Additional file 1: Table S1 and Figure S1. Principal component analysis of all analyzed samples for all T-cell populations revealed no critical outlier (Additional file 1: Figure S5 and Figure S2)

### Time-resolved transcriptome-wide changes depict common trends across CD4+ T-cell populations

For each CD4+ T-cell population in the *Discovery Set* and at each time point of activation, we assessed differential gene expression between polarized or activated T-cells without differentiating cytokines and CD4+ T-cells before activation. We declared a gene as significantly differentially expressed (DE) if the FDR-adjusted p-value was <0.05. The maximum number of DE genes among all 5 CD4+ T-cell populations was 3,641 after 24 hours of activation (Fig. 2A and see Additional file 1: Figure S7 for a more detailed overview). We observed coherent signs of fold changes for more than 96.5% of all genes identified as DE in at least 2 populations (Fig. 2B).

**Fig. 2.**
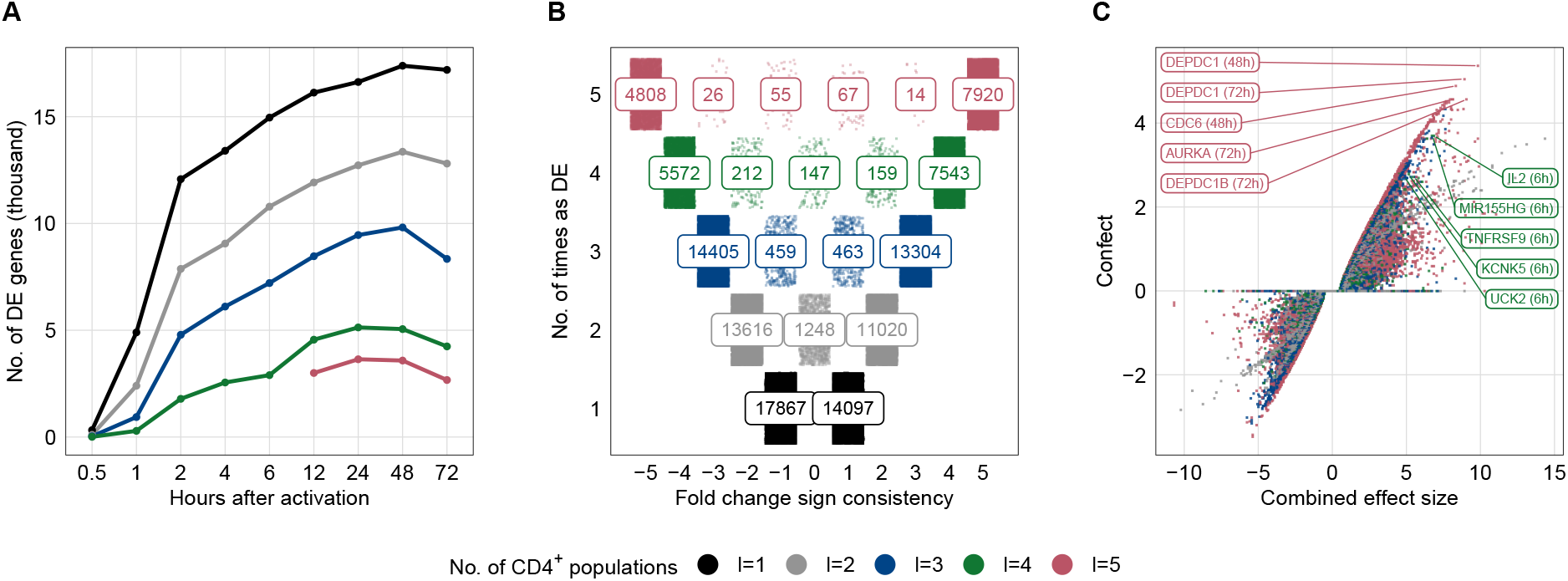
Common trends across CD4+ T-cell populations from the Discovery Set. **(A)** Inverse cumulative distributions of DE genes (FDR <0.05) in the *Discovery Set* for each time point during activation compared to unactivated CD4+ T-cells. Colors represent the number of T-cell populations in which genes were significantly differentially expressed. Analysis time points 12 to 72 hours were available for all 5 CD4+ T-cell populations. **(B)** Depicting the number of T-cell populations in which genes were identified as DE (y-axis) and the number of genes with consistent fold changes across T-cell populations (x-axis). For example, a gene that is significantly differentially expressed in 3 T-cell populations after 12 hours of activation, of which it is down-regulated in 2 populations, will get a fold change sign consistency of 1+(−2) = -1 (x-axis). An example of how to read the numbers: 7920 genes (top right) were identified as DE in all 5 T-cell populations. These DE genes were also upregulated in all 5 T-cell populations at each contrast (e.g., 12h vs. 0h). **(C)** Shown are DE genes with a combined effect size identified in the meta-analysis using a random effect model in at least 2 T-cell populations. The x-axis represents the combined effect size, the y-axis the “confect” value. Genes that do not show a significant combined effect size (FDR >0.05) have a “confect” value of 0. Gene labels for the 5 highest ranked “confect” values from the meta-analysis across 4 (0.5 to 6 hours, highlighted in green) and 5 (12 to 72 hours, highlighted in red) T-cell populations are shown.

Next, we conducted a meta-analysis to identify DE genes with a significant combined effect across CD4+ T-cell populations in the *Discovery Set*. For all DE genes at least 2 populations at the same time point of activation, we calculated Hedges’ g values as standardized effect sizes. Based on the Hedges’ g values, a combined effect size was estimated by applying a random-effects model. With an FDR <0.05, we identified 12,581 DE genes with a significant combined effect size across all 5 CD4+ T-cell populations after 12 to 72 hours of activation, the majority (90.4%) of which were in at least 2 time points. Overall, 94.2% of all genes that were DE in at least 4 CD4+ T-cell populations had also a significant combined effect size (Additional file 1: Figure S8D).

For each DE gene with a combined effect size, we calculated a confident effect size or “confect” (21). The “confect” values provide an inner bound on combined effect sizes while maintaining a given FDR <0.05. In this way, we ranked each gene according to its “confect”. This is a more conservative way than ranking by raw effect size because it avoids ranking genes by highest effect size, which can have a large within-group variability (see Additional file 1: Figure S8A).

The 5 highest ranked genes from the meta-analysis across 4 (0.5 to 6 hours) and 5 (12 to 72 hours) T-cell populations are depicted in Fig. 2C. IL2, TNF Receptor Superfamily Member 9 (TNFRSF9) and the miR-155 host gene (MIR155HG) were among the 5 highest ranked genes after 6 hours of activation across 4 T-cell populations. The most prominent genes across 5 T-cell populations included cell cycle-associated genes such as AURKA, CDC6, DEPDC1B, and DEPDC1 (22–25). The 20 highest ranked genes with a significant combined effect size and corresponding forest plots for each activation time point are described in Additional file 1: Figure S10. The 30 highest enriched Reactome pathways and GO terms (FDR <0.05) for DE genes with a significant combined effect size are shown in Additional file 1: Figure S11 and S12. Antigen stimulation of T-cell receptor (TCR) signaling to nuclear factor kappa B (NF-*κ*B) is required for T-cell proliferation and polarization and is represented among the enriched Reactome pathways. All DE genes enriched in TCR signaling are shown in Additional file 1: Figure S13. GO terms of T-cell activation were significantly enriched mainly after one hour of activation.

### Unsupervised decomposition revealed coherent metagenes across CD4+ T-cell populations in the *Discovery Set*

To identify coherent temporal expression profiles for each CD4+ T-cell population in the *Discovery Set*, we employed non-negative matrix factorization (NMF), an unsupervised decomposition approach. NMF allows a part-based representation by reducing dimensionality of the expression data and describing the samples as a composition of metagenes (i.e., sets of genes with similar expression pattern across time points). Based on consensus clustering as qualitative measurement combined with the cophenetic correlation coefficients and silhouette scores as quantitative measures, NMF resulted in the greatest stability with 3-5 metagenes for each CD4+ T-cell population (Additional file 1: Figure S15).

The resulting pattern matrix represents the relative weights of metagene k in sample m. A graphical representation of the metagene patterns is depicted in Fig. 3A. We observed different temporal metagene profiles for considered time points. All CD4+ T-cell populations were characterized by an early and late sustained response to activation and at least one intermediate expression pattern. In addition, metagenes showed consistent temporal peaks across all populations, which justifies an alignment of metagenes between CD4+ populations.

**Fig. 3.**
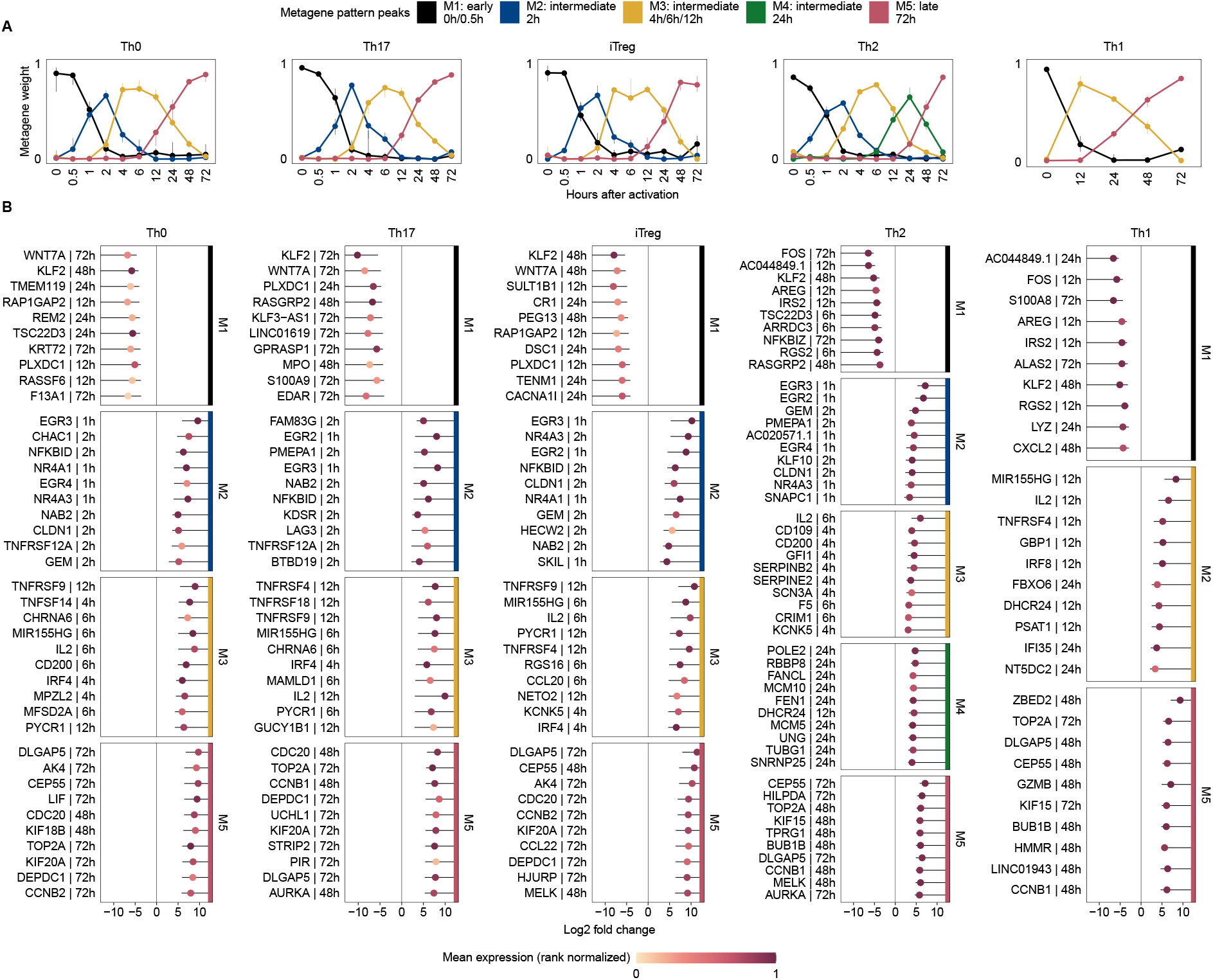
Temporal profiles obtained from NMF. **(A)** The pattern matrix for each CD4+ T-cell population from the Discovery Set is shown as continuous profiles, with samples assigned to time points of activation. We scaled each column in the pattern matrix to sum up to one. Dots depict median weights for all samples from identical analysis time points. Vertical lines represent interquartile ranges. We annotated and colored the metagenes based on their maximum median values across all analysis time points. The time point with maximum median value is depicted in the legend. **(B)** Top 10 genes associated with metagenes for each T-cell population. For each CD4+ T-cell population and gene used for NMF, we used the highest absolute “confect” value estimated in the DGEA across all contrasts (e.g., 12h vs. 0h). Genes are ranked by “confect” values. Dots represent log2 fold changes for contrasts with highest absolute “confect” value. The time point to the right of the gene represents the contrast with the highest absolute “confect” value. For example, “12h” represents following contrast: a sample group activated for 12 hours compared to unactivated samples of the same group. Color of dots correspond to rank normalized average expression values of the activation group in the contrast with the highest absolute “confect”. The inner end of the horizontal line shows the “confect” value (inner confidence bound). NFKBID denotes NF-*κ*B.

The gene signature matrix provided gene sets for each metagene, which can be linked to biological processes by enrichment analysis or functional annotation. The top 10 significantly enriched Reactome pathways and Gene Ontology (GO) terms are shown in Additional file 1: Figure S17 and S18.

FOXO-mediated transcription was identified in early response metagenes M1 in 4 out of 5 CD4+ T-cell populations. T-cell activation GO terms were significantly enriched in the intermediate metagene M2 with an expression pattern peak at 2 hours, while metabolic processes were enriched in intermediate metagene M3 with an expression pattern peak at 4 to 12 hours. Gene sets from the late response metagenes M5 were strongly enriched for cell cycle associated GO terms and pathways from the Reactome database. The 10 highest ranked genes for each metagene and T-cell population are shown in Fig. 3B (Additional file 1: Figure S16 for the 25 highest ranked genes).

### Consensus gene expression profiles for CD4+ T-cells

To obtain more robust time series and to explore the consensus transcriptional regulators of T-cell activation, we combined metagenes with similar expression patterns among CD4+ T-cell populations in the *Discovery Set*. This means that the metagenes had to be temporally consistent across all CD4+ T-cell populations. This resulted in a set of genes with similar expression profiles over the course across CD4+ T-cell populations. Furthermore, to refine the variety of gene expression profiles, we defined identity and shared metagene. For illustrative purposes, we developed a metagene landscape reflecting the relative relationship of genes and samples to metagenes. We used the combined pattern matrices from all CD4+ T-cell populations to embed the temporally consistent metagenes into a two-dimensional space. To embed the relative relation between metagenes to genes as density map into the metagenes landscape, we used the weights from the gene signature matrix from each CD4+ T-cell population. Samples were embedded relative to the metagenes using the pattern matrix. The resulted metagene landscape for the *Discovery Set* is shown in Fig. 4A. As an example, we used IL2, IL2RA, CD4, and NF-κB (NFKBID), which are part of our consensus gene expression profiles of the *Discovery Set* and depicted the mean metagene weights from the CD4+ T-cell populations as dots in the metagene landscape. CD4 was highly expressed in all T-cell populations. However, we observed that CD4 embedded between metagene M1 and M5, which represents the early and late response metagene and was expressed at a higher level compared to the other samples (Fig. 4B). On the other hand, IL2 was embedded near metagene M3, which represents an intermediate response metagene and is expressed at higher levels in samples at 6 to 24 hours after activation (Fig. 4B) compared with the other samples. This indicates that there are set of genes associated with more than one metagene. Therefore, we extended the definition of metagenes and genes associated with them. Genes with high weights in one metagene from the gene signature matrix were referred to as genes linked to identity metagenes. Genes with higher weights in more than one metagene were referred to as genes linked to shared metagenes (see methods).

**Fig. 4.**
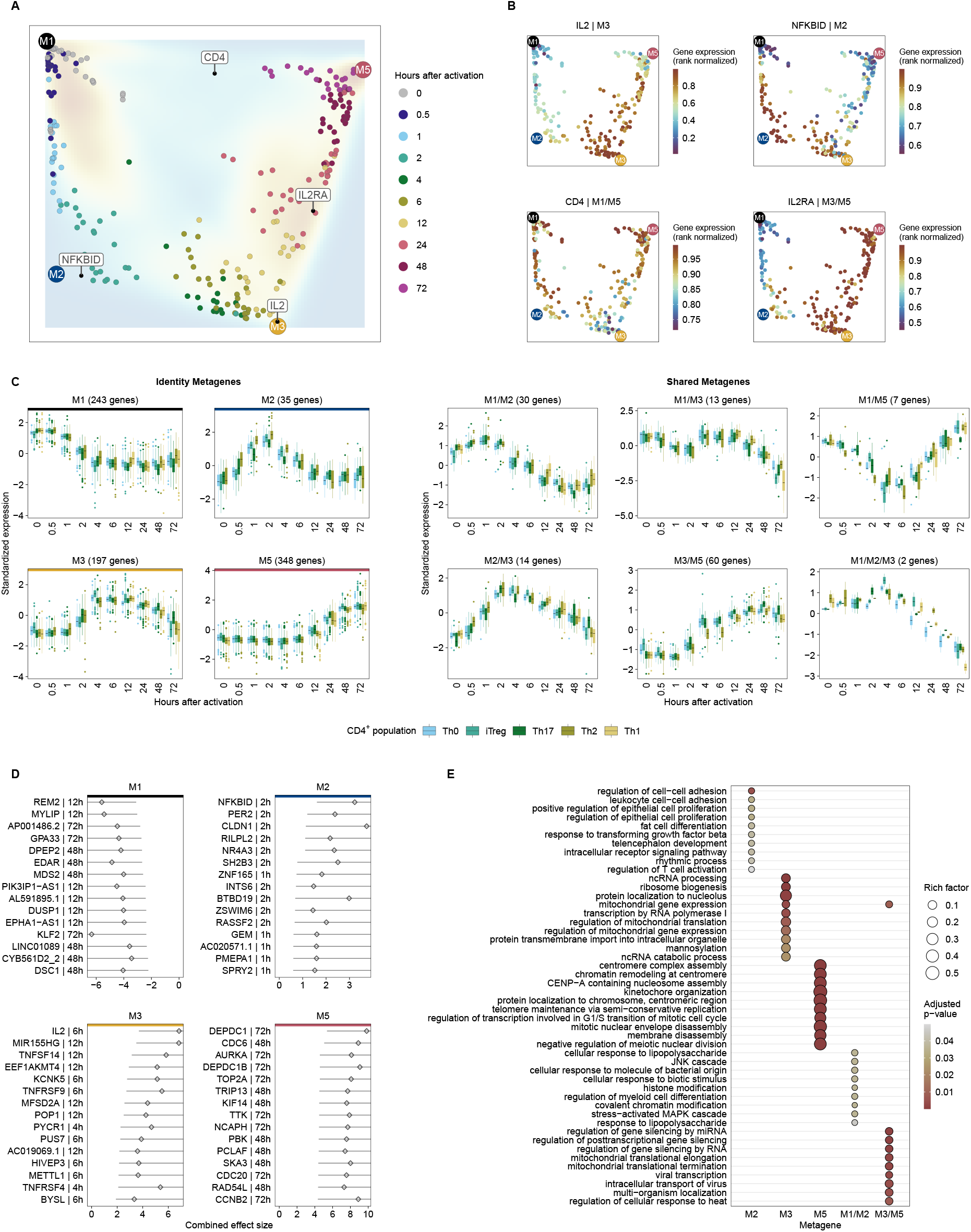
**(A)** Metagene landscape. We embedded all samples relative to temporally coherent metagenes using the pattern matrix. All genes that we used for NMF were embedded relative to the temporally coherent metagenes using the gene signature matrix and depicted as density map. For IL2, IL2RA, CD4, and NF-*κ*B (NFKBID) we calculated the average metagene weights across the CD4+ T-cell populations and depicted them as dots in the metagene landscape. **(B)** Each sample is colored by the rank normalized gene expression of the corresponding gene. **(C)** We grouped the consensus expression profiles over the course by genes associated with identity and shared metagenes. Each boxplot represents one CD4+ T-cell population from the *Discovery Set*. The y-axis depicts standardized median expression of genes from samples with identical analysis time points. The number in parentheses represents the number of genes for the corresponding metagene. **(D)** Top 15 genes associated with identity metagenes. For each gene belonging to the consistent metagenes, the time point with the highest absolute “confect” value estimated in the meta-analysis used. Genes were then ranked according to their absolute highest “confect” value. Diamonds represent combined effect size from the meta-analysis for the activation time point with the highest absolute “confect” value. The time point to the right of the gene represents the time point of the highest absolute “confect” value. The inner end the of horizontal line shows the “confect” value (inner confidence bound). **(E)** The top 10 (sorted by rich factor) significantly enriched GO terms of biological processes (FDR <0.05) of metagene associated genes identified by enrichment analysis. The dot size indicates the rich factor, which is the number of metagene associated genes in the GO term divided by the number of background genes of the term. Colors indicate adjusted p-values of significantly enriched GO terms.

Furthermore, we used the “Housekeeping and Reference Transcript Atlas database” (26) to remove 183 housekeeping genes from the consensus signatures constitutively expressed in different tissues and cell types (Fig. 5F).

**Fig. 5.**
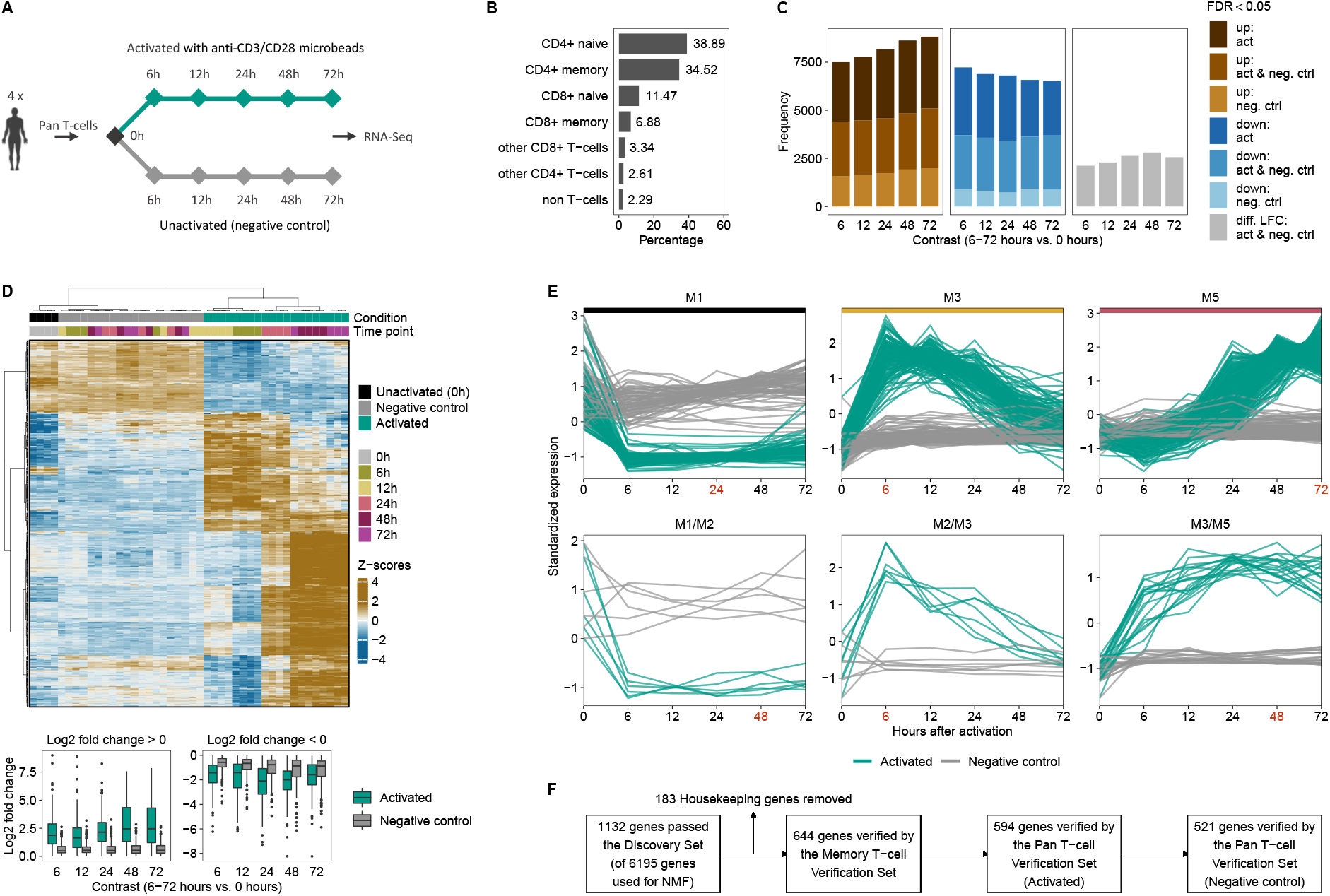
Verification of consensus temporal gene signature. **(A)** Experimental design for the *Pan T-cell Verifications Set*. We performed RNA Sequencing of Pan T-cells from 4 healthy donors at 5 different time points after anti-CD3/anti-CD28 activation (6 to 72 hours) and of unactivated (0 hours) T-cells. In addition, we sequenced Pan T-cells for the same time points without activation as negative controls. **(B)** Depicted are the fractions of Pan T-cell populations before activation. 200.000 cells were analyzed using seven human blood donors. **(C)** For each contrast (6 to 72 hours of activation and without activation vs. 0h), we performed a DGEA. The brown and blue bar plots depict the number of DE genes (FDR *<*0.05) for each contrast that are up- (brown) or down-regulated (blue) under activated (act), unactivated (neg. ctrl) conditions, or both (act & neg. ctrl). Gray bar plots show the number of DE genes under activated and unactivated conditions without consistent log2 fold change. **(D)** Hierarchical clustering of DE genes from the *Pan T-cell Verifications Set* in activated and unactivated conditions. Only genes from the consensus signatures that passed the activation kinetics of both Verification Sets are shown. Euclidean distance with Ward clustering was applied to visualize similarity between samples. Each column represents a sample, each row represents a gene. The y-axis depicts the standardized median CPM expression values of genes from samples with identical analysis time points. Bottom panel: For each contrast and condition, boxplots of log2 fold changes are shown. **(E)** For the Pan T-cell Verification Set the temporal expression pattern of genes from the consensus signatures that passed the 2 Verification Sets (activation and negative control kinetics) are shown. CPM values were z-score standardized. Identity metagenes are highlighted with colored horizontal bars (M1, M3 and M5). The red colored time points indicate the maximum centroid threshold. Only metagenes with more than 5 genes are shown. **(F)** Flowchart depicting the number of genes passing each filter step.

The resulting consensus gene expression profiles, separated into identity and shared metagenes associated genes, are shown in Fig. 4C. The top 15 genes associated with identity metagenes are shown in Fig. 4D. In addition to the 4 identity metagenes, we were able to identify another 6 shared metagenes. The top significantly enriched GO term for biological processes (FDR <0.05) for genes linked to identity and shared metagenes indicate a distinct functional context (Fig. 4E). Metagene M2 is enriched with GO terms of T-cell activation regulation, intracellular receptor signaling pathway and leukocyte cell-cell adhesion. Genes from the shared metagene M1/M2 are enriched with GO terms of cellular responses to stimuli and MAPK cascade. Metagene M3 and M5 are linked to metabolic processes and cell cycle terms, respectively (see Additional file 1: Figure S20 for the top enriched Reactome pathways and Additional file 2 for expression profiles of all genes from the consensus signatures and all associated pathways/terms).

### Verification and mapping to a wider variety of T-cell transcriptional landscape

To verify the consensus gene expression profiles and additionally represent a wider variety of the T-cell landscape, we used 2 *Verification Sets*. Firstly, we used RNA-Seq activation kinetics of memory CD4+ T-cells from Gutierrez-Arcelus et al. (15) (referred to as *Memory T-cell Verification Set*). Secondly, we performed time series RNA-seq of activated and unactivated Pan T-cells from 4 healthy donors (Fig. 5A). We used this dataset as *Pan T-cell Verifications Set*. The fractions of the Pan T-cell populations are shown in Fig. 5B. To find stable metagenes for each activation kinetic of the 2 *Verification Sets*, we used the same procedure as for the *Discovery Set* (Additional file 1: Figure S15). The temporal profiles of the *Verification Sets* calculated from the pattern matrix and metagene associated genes are shown in Additional file 1: Figure S21.

Next, we compared activated and unactivated (negative control) gene expression kinetics of T-cells from the *Pan T-cell Verifications Set*. We performed DGEA to find differentially expressed genes for activated and unactivated T cells at 6 to 72 hours compared to unactivated T-cells at 0 hours. DGEA revealed that across all contrasts (6 to 72 hours vs. 0 hours), an average of about 2,700 genes were significantly up- or downregulated (FDR *<*0.05) in both anti-CD3/CD28 bead activated and unactivated conditions (Figure 5C). Significantly enriched Reactome pathways for DE genes under activated and unactivated conditions are shown in Additional file 1:

Figure S24. For the unactivated conditions, we observed that pathways associated with metabolic processes, transcription and proliferation were significantly enriched for downregulated DE genes. Even though, mostly disjunct enriched pathways were observed for upregulated DE genes under activated conditions, they are also associated with metabolic processes, transcription, and proliferation. Heatmap visualization of DE genes in activated and unactivated conditions showed that temporal impulse patterns were not observed under unactivated conditions (Additional file 1: Figure S25). Although we observed two sustained temporal patterns under unactivated conditions, samples with identical analysis time points were not clustered together.

All genes from the consensus signatures that we identified based on Discovery Set were verified for temporal consistency by the *Verification Sets* (see methods). This results in 594 genes. However, of those 457 also showed significant expression changes in at least one contrast in negative controls. Unsupervised clustering of those genes resulted in distinct expression clusters according to conditions, but partly similar temporal patterns among conditions (see Fig. 5D). The majority showed only minor log2 fold changes in negative controls compared to unactivated condition at 0 hours. About 37% of all DE genes had a log2 fold change >1 in at least one contrast under unactivated conditions (Fig. 5D, bottom panel) Therefore, we compared activated T-cells with negative controls at the time point with maximum absolute distance to the centroid of the metagenes (see methods). We only retained genes from the consensus signatures with significant differential expression between activated and negative controls (FDR <0.05) and an absolute “confect” value >1 at the selected time point. The temporal expression patterns of genes passing this filter step are shown in Fig. 5E (see Additional file 1: Figure S26 for all metagenes and Figure S27 for genes expression profiles not passing this filter step).

Overall, passing all filtering steps this resulted in 521 genes included in the final consensus gene signatures (Fig. 5F). For the highest ranked genes, we provide an overview linking genes to the metagenes and the top enriched Reactome pathways/GO terms in Additional Figure S28 and S29 (see Additional file 2, which shows an overview of all genes and their temporal expression as an interactive HTML document). In addition, we also provide comprehensive characterization of genes passing the Memory T-cell Verification Set in Additional file 1: Figure S22 and Additional file 2, respectively.

### Consensus gene expression profiles are enriched in CAR T-cell products from patients with low-grade ICANS

We re-analyzed a scRNA-Seq study of autologous axicabtagene ciloleucel (axi-cel) anti-CD19 CAR T-cell infusion products (including non-transduced T-cells) from 24 patients with LBCL (27). As adverse event related to CAR T-cell therapy, 12 patients developed high-grade immune effector cell-associated neurotoxicity syndrome (ICANS, grade 3-4), whereas the other 12 patients developed low-grade ICANS (grade 0-2). The aim of this re-analysis was to investigate whether there is a significant difference in the expression of the temporal consensus gene signatures between low-grade and high-grade ICANS. Deng et al. observed only few significant differences in the gene expression of CAR+ T-cells compared to non-transduced T-cells. Therefore, they did not separate CAR+ and CAR-T-cells in their analyses, which is also our strategy in the following.

Firstly, we analyzed whether metagenes were associated with cell clusters. Among all CD8+CD4- and CD8-CD4+ T-cells, 3 metagenes (M5, M3 and M3/M5) with at least 10 genes were present in the scRNA-Seq data (Fig. 6A). We performed integration analysis to account for the heterogeneity between the samples, followed by unsupervised clustering for cell state identification. The cells were then projected into a two-dimensional space using tSNE and colored by standardized average expression of the present metagenes for each cluster (Fig. 6B). We performed the same procedure with known T-cell state markers (see methods) and markers associated with T-cell molecular mechanisms (28) (Additional file 1: Figure S30A).

**Fig. 6.**
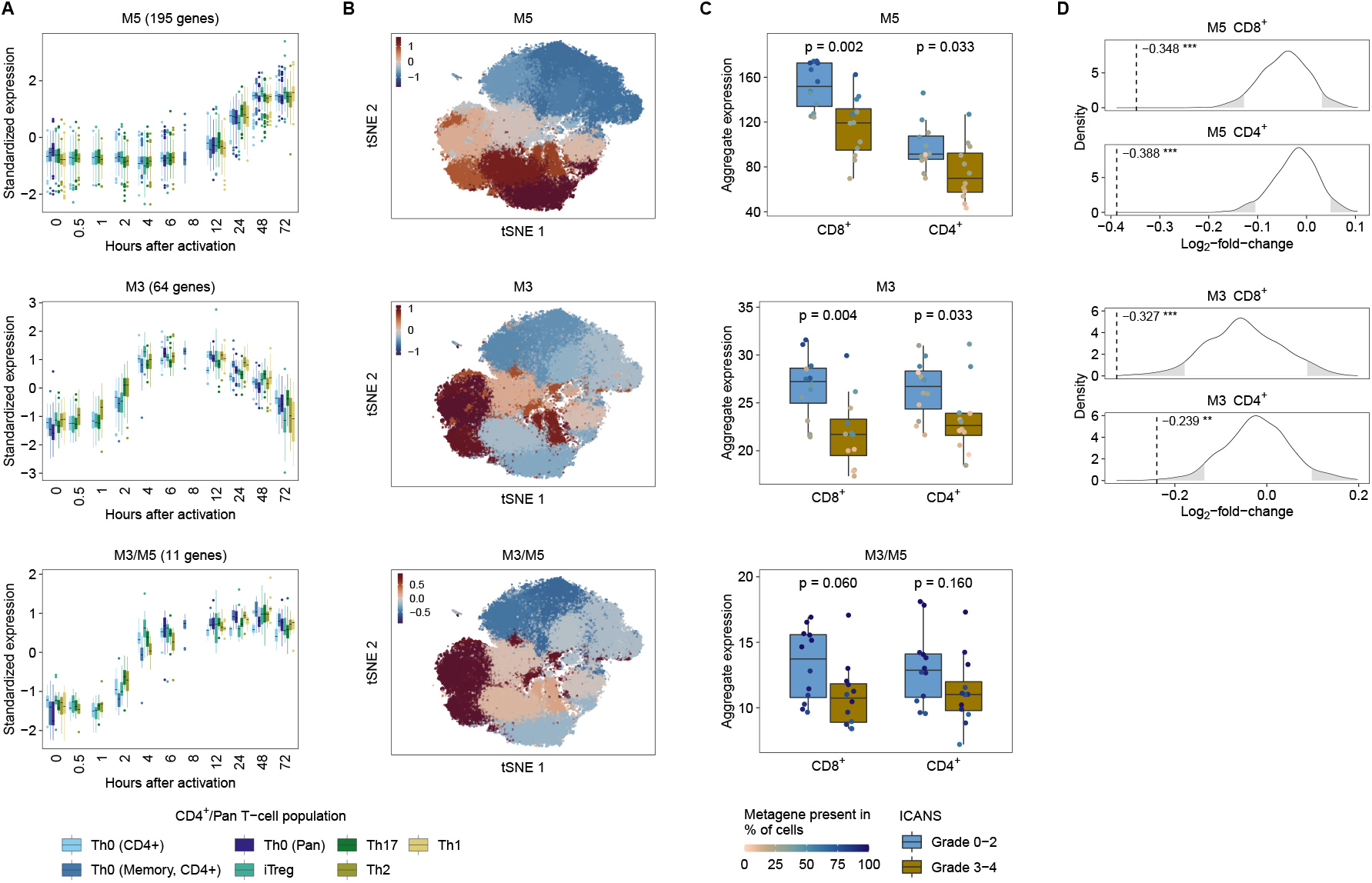
Re-analysis of data scRNA-Seq data of autologous anti-CD19 CAR T-cell infusion products from 24 patients with LBCL. **(A-C)** Metagenes with at least 10 genes among the most highly variable genes in the scRNA-Seq dataset were analyzed. **(A)** The boxplots in the left panel show grouped consensus expression profiles over the course by genes associated with identity and shared metagenes. Each boxplot represents one T-cell population from the Discovery and Verification Sets. The y-axis depicts the standardized median expression of genes from samples with identical analysis time points. The number in brackets above the boxplots indicates the number of highly variable genes of the metagene present in the scRNA-Seq data. **(B)** We embedded CD8+CD4- and CD8-CD4+ T-cells from 24 patients into a two-dimensional space by the t-distributed stochastic neighbor embedding (tSNE) method. The colors indicate the standardized average expression of the metagenes for each cluster (the same value is assigned to all cells in a cluster). **(C)** For each metagene and patient, we calculated aggregated expression (summed average expression) for CD8+CD4- and CD8-CD4+ cells. Differences in aggregated expression between patients with low- and high-grade ICANS were evaluated using the Wilcoxon rank-sum test. The colors of the dots in the boxplots indicate the percentage of cells for each patient in which the corresponding metagene is present. We considered a metagene as present in a cell if at least 25% of all associated genes had at least one UMI count. **(D)** For each metagene and T-cell population that was significant (p <0.05) in the Wilcoxon rank-sum test, we generated a null distribution in order to confirm the results (see methods). Dashed vertical lines indicate median log2 fold change of aggregated expression between low- and high-grade ICANS patients of the metagene. Rejection regions (empirical p-value <0.05) are highlighted in grey (* p <0.05, ** p <0.01, *** p <0.001).

We observed that metagene M5 is present in cell clusters associated with the cell cycle signatures G1/S and G2/M. (Additional file 1: Figure S30A). Metagenes M3 and M3/M5 were present in cell clusters linked to TCA cycle, glycolysis, Treg and Hypoxia/HIF signatures. All metagenes that were linked to cell clusters were associated with the CD8 lineage (Additional file 1: Figure S30B).

For each metagene and patient, we calculated the aggregated expression of CD8+CD4- and CD8-CD4+ cells. We evaluated differences in aggregated expression between patients with low- and high-grade ICANS using the Wilcoxon rank-sum test (Fig. 6C). We observed significant differences for metagene M5 (CD8+ p = 0.002, and CD4+ p = 0.033) and M3 (CD8+ p = 0.004 and CD4+ p = 0.033). To confirm this finding, we generated a null distribution by randomly drawing gene sets of the same size as the metagenes (see methods) and found that metagenes performed significantly (p <0.05) better than randomly generated gene sets (Fig. 6D). We also tested whether patient characteristics correlate with aggregate expression of the significant metagenes. Patient age correlated significantly (p-value = 0.045) with the aggregated expression of the gene set of metagene M3 in CD8+ T-cells (Additional file 1: Figure S32).

We conducted the same analysis as above for the adverse event, cytokine release syndrome (CRS) and also for patient outcome. However, when comparing aggregate expression between patients with low-grade and high-grade CRS or between complete response and partial response/progressive disease, we did not observe significant differences (Additional file 1: Figure S33).

For T-cell state markers as well as markers for T-cell molecular mechanisms associated with the same cell clusters as the metagenes, only the cell cycle signatures G1/S and G2/M showed significant differences between low- and high-grade ICANS in CD8+ cells (Additional file 1: Figure S31A, Wilcoxon rank-sum test). However, only the G1/S signature performed significantly better than random sets (Additional file 1: Figure S31B). For markers not associated with the same cell clusters as the metagenes, the pro-inflammatory CD8+ signature, the cytolytic effector pathway and the glucose deprivation signature showed also significant differences in aggregated expression between patients with low- and high-grade ICANS (Additional file 1: Figure S31A, Wilcoxon rank-sum test). All three were confirmed by comparison to randomly generated gene sets (Additional file 1: Figure S31B).

## Discussion

We analyzed kinetic changes in gene expression profiles of human T-cells isolated from PBMCs or cord blood after activation with anti-CD3/anti-CD28 only or after subsequent polarization with cytokines. To provide temporal details of the coordinated multiple functions before and after activation of T-cells, we used in-vitro kinetic models profiled by RNA sequencing and microarray platforms. We used a *Discovery*- and two *Verification Sets* consisting of multiple T-cell populations to find DE genes after activation. In the *Discovery Set*, we identified a reasonable large number of genes with consistent signs of log2 fold change across different T-cell populations (Fig. 2B). This points to non-negligible overlaps in transcriptional responses upon activation of T-cells, no matter of polarization conditions. We obtained genes with a significant combined effect size across different CD4+ T-cell populations by a meta-analysis. Based on these genes, we conducted NMF to aggregate the temporal expression profile of each T-cells population into metagenes, i.e., sets of temporally co-expressed genes.

NMF does not inherently preserve relative temporal ordering in pattern matrix, because metagenes are treated as nominal variables. Hence, we ordered samples in the pattern matrices according to time points in order to classify the expression pattern of each T-cell population into early, intermediate and late response metagenes

Krüppel-like factor 2 (KLF2) is among the highest ranked genes associated with the early-response metagene (M1) across all T-cell populations (Figure S29). KLF2 plays an important role in regulation of the T-cell homeostasis by promoting naïve-T-cell quiescence (29). Its expression decreases rapidly after TCR activation, and it has been observed that KLF2 protein degradation and subsequent loss of KLF2 mRNA occurs during T-cell stimulation (29, 30). KLF2 and NF-*κ*B are reciprocal antagonists (31, 32). NF-*κ*B is the highest ranked gene linked to the intermediate metagene M2 (Figure S28). The expression patterns of metagene M1 and M2 here illustrate the dual effect of KLF2 and NF-*κ*B.

Members of the early growth response family (EGR2 and EGR3) are among the prominent genes of the intermediate response metagene M2. These transcription factors are targets of the NFAT transcription factors (33, 34), act as transcriptional repressors and impair IL2 transcription in anergy (35). IL2 is the highest ranked gene for the intermediate metagene M3 (pattern peak at 6-12h) in the *Discovery Set* and *Pan T-cell Verifications Set*. However, in the *Memory T-cell Verification Set*, IL2 is associated with the intermediate metagene M2/M3 (pattern peak at 2-4h). This classification might be of biological origin because memory T-cells were activated in the *Memory T-cell Verification Set*, which may lead to a faster response to the activation signal. Otherwise, we were not able to compare the expression peak of IL2 in the *Memory T-cell Verification Set* with the *Discovery Set* or *Pan T-cell Verifications Set*, because the *Memory T-cell Verification Set* lacks the activation time point at 6 hours. In addition to IL2, this observation was also made for TNFRSF9, an activation-induced co-stimulatory molecule of the tumor necrosis factor receptor superfamily that is upregulated on activated T cells and antigen-presenting cells, including dendritic cells, macrophages and B cells (36–40) (for expression profile see Additional file 2).

MIR155HG, linked to metagene M3, is associated with immune cell infiltration and immune checkpoint molecule expression in several cancers and has been proposed as a prognostic marker (41). The highest ranked gene for the shared metagene M3/M5 is Interleukin-2 receptor (IL2RA), which play a crucial role in immune homeostasis (42) (Figure S28). AURKA, CDC6, TOP2A, DEPDC1B, and DEPDC1 are the most prominent genes for the late response metagene M5, which are mainly involved in cell cycle processes (22–25).

Although filtering metagenes specific to a T-cell population provides added value, we did not include this analysis in our study. Since the Th1, Th2 and Th17 populations consist of only three biological replicates, we cannot guarantee sufficient robustness. Further kinetic studies of these populations with overlapping time points are needed to develop T-cell population specific time series signatures.

For the *Pan T-cell Verifications Set*, in addition to the activation kinetics, we also sequenced unactivated Pan T-cells after 6 to 72 hours, which we used as negative controls. With this strategy, we were able to exclude genes with similar expression in negative control and activated Pan T-cells from the consensus gene signatures. It is worth noting that we observed an average of 10,760 DE genes across the contrasts for the negative controls (Fig. 5C). This indicates an unspecific regulation due to cultivation effects and emphasizes the importance of using matched negative controls in time series experiments.

Our study provides temporal consensus signatures of T-cell regulatory dynamics from healthy donors that could also be useful for defining T-cell states in disease, as proposed in Szabo et al.(43). We demonstrated this by re-analysis of a scRNA-Seq study by Deng et al. (27) in which autologous axicel anti-CD19 CAR T-cell infusion products from 24 patients with LBCL were assessed by single-cell RNA sequencing. Metagenes M3 and M5, which are associated with metabolic processes and proliferation, respectively, were significantly enriched in patients with low-grade ICANS compared to patients with high-grade ICANS. These observations are novel with respect to ICANs but need to be verified by further experiments, as the metagenes were trained using samples derived from healthy donors and not from heavily pretreated patients, as is the case with LBLC patients (27). We did not find a significant correlation between metagenes and CRS. However, this could be due to the fact that the high-grade CRS group (grade 3-4) included 4 patients, while the low-grade group (grade 1-2) consisted of 18 patients. Deng et al. showed that CAR T-cell products which had a significant enrichment of cells with a CD8 T-cell exhaustion lead to a partial response or progressive disease, whereas cell products enriched with the memory phenotype were correlated to a complete remission of LBCL. This phenomenon was also confirmed in chronic lymphoblastic leukemia patients treated with anti-CD19 CART cells (44). However, we did not find a significant correlation between metagenes and patient outcome. Overall, our activation signature could be helpful for evaluating CAR T-cell products prior or after the manufacturing process.

Of note, expression peaks of the signatures are interpreted relative to the unactivated condition since we did not filter genes according to low expression prior to activation.

We are aware that the consensus signatures developed are based on an artificial immune response where T-cells were activated with anti-CD3/anti-CD28 coated beads. This experimental design only partially reflects human immune response in-vivo. Therefore, the expression pattern of the genes from our signatures might behave differently under in-vivo conditions. For this reason, we are investigating the temporal resolution of the recall immune response to antigens in an ongoing study.

Further, with the increasing number of available temporal transcriptome-wide studies, we expect to transfer our concept of defining and validating temporal biomarker signatures to differentiation processes of distinct T-cell populations. This was not the aim of the current study due to small number of replicates. Low sample sizes limit separation of data sets into disjunct training and verification data sets, a substantial requirement for the development of biomarker gene signatures.

## Conclusion

In this study, we identified and verified general biomarker signatures robustly evaluating T-cell activation in a time-resolved manner. It describes general temporal changes in gene expression patterns upon T-cell activation no matter of the polarization condition. It serves as a resource for studies of human T-cell immunity in disease or immunotherapies to classify T-cell states and may be used to define optimal T-cell biomarkers in dependence of experimental analysis time points. For easy access and interpretation of the consensus gene signatures, we provide an interactive HTML document (Additional file 2). In addition, the developed method for analyzing time-series transcriptome data can be applied to other cell types.

## Methods

### Data sources used for the development of the consensus gene signature

We downloaded the publicly available RNA-Seq and microarray data sets from the Sequence Read Archive (SRA) (45) and NBCI’s Gene Expression Omnibus (GEO) (46), respectively. The analysis consists of one *Discovery* and two *Verification Sets*. For the *Discovery Set* we used following sources: Raw sequencing data in FASTQ format from the RNA-Seq projects (GSE52260, GSE90569, GSE94396, GSE96538) (12–14) were obtained using the `prefetch` and `fastq-dump` commands in the `SRA Toolkit` v2.9.2 (47). Raw CEL files from Affymetrix Human Genome U133 Plus 2.0 Arrays for GSE17974 (10) and GSE32959 (11) were obtained using the `R` package `GEOquery` v2.58.0 (48).

Since some of the analyzed datasets examined gene expression of CD4+ T-cells at multiple time points of activation, we decided that one time point must be available for at least 3 CD4+ T-cell populations. With exception of CD4+ cells cultured under Th1 cell polarization condition (11) overall 9 different time points were analyzed for each population. Activated (Th0) CD4+ T-cells comprised 4, iTreg 3 and the other populations one dataset each. A detailed description of the experimental conditions and RNA-Seq or microarray technical specifications for each dataset is shown in Additional file 1: Table S1 and Figure S1.

We downloaded the RNA-Seq gene counts from the study Gutierrez-Arcelus et al. (GSE140244) (15) and used them as Memory T-cell Verification Set.

For the Pan T-cell *Verification Set* (GSE197067), we activated Pan T-cells from 4 healthy individuals with anti-CD3/CD28 beads (for MACS isolation strategy of Pan T-cells, see Additional file 1). Briefly, 2×10^5^ Pan T-cells were seeded in a 96 U-well plate with quadruplicates per condition. 2×10^5^ Dynabeads human T-Activator CD3/CD28 (Gibco, 11132D) per sample were washed with PIB and were then resuspended in T cell medium (CG-DC-medium supplemented with 5% human AB serum and 1% penicillin/streptomycin). 2×10^5^ CD3/CD28 beads were added to activate T-cells and wells were filled up to 200 μL total volume. Unstimulated T-cells were used as controls. T-cells were harvested at different time points (0h, 6h, 12h, 24h, 48h, 72h). Therefore, unactivated and activated T-cells were collected and centrifuged (10 min, 300 xg, room temperature). 700 μL Trizol were added and sampels were stored at - 80 ^*°*^C till RNA isolation. We performed RNA-seq before activation (0 hours) and 6, 12, 24, 48, 72 hours after activation. In addition, as a control, we sequenced Pan T-cells for the same time series without activation condition. For experimental details, see Additional file 1.

### Pre-processing and normalization

To facilitate the multi-step analysis of the RNA sequencing datasets, we applied the workflow-manager uap v0.0.1 (49). A detailed description of all processing steps from FASTQ files to gene quantification can be found in Additional file 1. The according configuration files for uap are available in Additional file 3. For pre-processing of microarray datasets, we used the R package affy v1.68.0. Gene expression of microarray and RNA-Seq datasets was quantified using the human reference gene annotation GENCODE v29. Expression data of the same T-cell population and platform were normalized in one junk (see Additional file 1 for more details).

### Differential gene expression analysis

We performed a differential gene expression analysis (DGEA) for each contrast (activated T-cells at a given time point compared to unactivated T-cells) using the empirical Bayes moderated t-test implemented in the package `limma` v3.46.0 (50, 51). For each contrast, p-values were adjusted using the method of Benjamini-Hochberg (or false discovery rate (FDR)). A gene was considered as significantly differentially expressed (DE) if the FDR-adjusted p-value was <0.05. DE genes were ranked using the `Topconfects` v1.6.0 `R`/`Bioconductor` package (21) (see Additional file 1 for a more detailed method description of the DGEA). The same procedure was carried out for the negative controls from the *Pan T-cell Verifications Set* with following contrasts: Firstly, unactivated Pan T-cell at 6 to 72 hours compared to unactivated Pan T-cells at 0 hours. Secondly, activated Pan T-cells at a given time point after activation compared to unactivated Pan T-cells at the same time point. A volcano plot representation of the DGEA is shown in Additional file 1: Figure S6.

### Meta-analyses

For each DE gene in the *Discovery Set* we calculated a standardized effect sizes as Hedges’ g values (52–54). Briefly, this involves log2 fold changes of the DE genes for activated compared to unactivated CD4+ T-cell populations at each analysis time point divided by the standard deviation as estimated with the empirical Bayes method in limma, followed by an adjustment for small sample sizes (see Additional file 1 for more details).

We estimated a combined effect size for each DE gene across the CD4+ T-cell populations and time points in the Discovery Set using a random effects model. We used a random effects model because we did not assume that there is one true effect size, which is shared by all the included datasets, but rather a range of true effect sizes with additional sources or variation, such as different platforms (RNA-Seq and Microarray). The model was fitted with the restricted maximum-likelihood estimator in the `R` package `metafor` v2.4.0 (55). We used the `Topconfects` v1.6.0 `R`/`Bioconductor` package (21) to to calculate for each estimated combined effect size and standard error a “confect” value. This “confect” value or confident effect size represents a confident inner bound of the calculated combined effect size by metafor while maintaining a given FDR of <0.05. We declared that at a given time point of activation, a DE gene had a significant combined effect size in at least 2 CD4+ T-cell populations if the FDR adjusted p-value was <0.05. To assess the amount of heterogeneity post-hoc the I2 statistic was calculated using the metafor package (see Additional file 1: Figure S8B).

### Unsupervised decomposition

To infer biological patterns over time, non-negative matrix factorization (NMF) was performed. We performed NMF for each T-cell population (including unactivated T-cells) from the *Discovery Set* and *Verification Set*s separately. For the *Discovery Set* the following set of genes were defined as input to NMF: All DE genes with a significant combined effect size in at least one contrast (e.g., 12 hours vs. 0 hours) across 4 (0.5 to 6 hours) and 5 (12 to 72 hours) T-cell populations. For the activation kinetics of the *Verification Set*s, we factorized the set of genes defined for the *Discovery Set*. However, we only used genes from this set that were also significantly differentially expressed in the *Verification Set*s. To preserve the linearity of NMF modeling, quantile normalized CPM values from RNA-Seq and normalized intensities from gene expression arrays were used in non-log space (56, 57). We used the R-package `NMF` v0.23.0 (58) with the algorithm from Brunet et al. (59) based on the Kullback-Leibler divergence as objective with multiplicative updates rules to solve the following approximation:

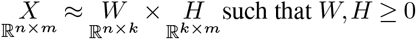

The aim is to factorize a non-negative matrix *X* into two lower rank matrices with strictly non-negative elements where *X* is the expression matrix whose rows contain the expression levels of the *n* genes in the *m* samples. Matrix *W* (referred to as gene signature matrix) has size *n* × *k*, where *n > k*. Each of the *k* columns defined a latent factor and was used to describe the original data in a sub-dimensional space. In the context of time series gene expression data, these latent factors were referred to as metagenes that reflect genes with similar expression patterns over time. Each entry *w*_*ij*_ contains the contribution for gene *i* to metagene *j*. Matrix *H* represented the pattern matrix and has size *k* × *m*, where the entry *h*_*ij*_ is the weight of metagene *i* in sample *j*.

The initialization of *H* and *W* was generated by random seeding where the entries of each factor are drawn from a uniform distribution within the same range as the entries in the matrix *X*.

To determine the optimal factorization rank *k* (number of metagenes) for each T-cell population we performed NMF from actual and randomized data by repeating the rank value in the interval 2–10. For each rank, 200 iterations were performed. We used consensus clustering as qualitative measurement, cophenetic correlation coefficients and average silhouette scores as a quantitative measure to assess stability of the clusters (59, 60) (see Additional file 1 for a more detailed method description). As proposed in Brunet et al. we selected the factorization rank where the magnitude of the cophenetic correlation coefficient begins to fall. In addition, we used the average silhouette scores to choose the optimal rank (see Additional file 1: Figure S15). After determining the optimal rank, the pattern and gene signature matrix was obtained from the factorization that achieved the lowest approximation error for the selected rank in 200 runs.

### Temporal categorization and visualization of metagenes

We used the entries in matrix *H*, representing the weights of metagene *i* in sample *j* for temporal annotation and visualization of metagenes (Fig. 3A). Each column in the matrix H was scaled to sum up to one. For each metagene *i* we calculated median weights for all samples from identical analysis time points. We annotated metagene *i* as early, intermediate, or late response metagene based on its maximum median weight across all analysis time points.

### Defining metagene associated genes

We used the gene signature matrix to assign DE genes to annotated metagenes. Each row in matrix *W* was min-max normalized such that the values, which contain the contribution of gene *i* to metagene *j*, are in the interval [0,1]. The resulted bimodal distribution for each T-cell population is shown in Figure S14. Gene *i* was associated with metagene *j* when the normalized weight *w*_*ij*_ was >0.5. In this way, we were able to identify genes that are specific to one metagene (*w*_*ij*_ == 1) and are referred to as identity metagenes or genes with higher weights (*>*0.5) in more than one metagene. In this case, we have referred to the metagenes as shared metagenes. We did not filter out genes post hoc because we factorized only informative genes, that is, genes with a significant combined effect size.

### Consensus gene expression profiles from the *Discovery Set* and verification

For the consensus gene signatures from the *Discovery Set*, the annotated metagenes had to be temporally consistent across all CD4+ T-cell populations. This means, for example, that all genes associated with the late-response metagene in one T-cell population must also be associated with the late-response metagene in all other T-cell populations. We used 2 Verification Sets to verify the consensus signatures (Fig. 4C) from the *Discovery Set* using the same procedure. Since only the analysis time points 12 to 72 hours for the Th1 population (*Discovery Set*) and 6 to 72 hours for the Th0 population (*Pan T-cell Verifications Set*) are available, we could not make any conclusion about the time course of expression with respect to intermediate metagene 2 (expression peak after 2 hours). Therefore, for the two populations mentioned, we excluded metagene 2 from this analysis.

For the *Pan T-cell Verifications Set*, we used the time series negative controls (unactivated Pan T-cells after 6 to 72 hours) to compare their gene expression profiles with the kinetics of the activated Pan T-cells. For each metagene from the *Discovery Set*, its centroid was calculated, defined as the median expression value of all genes for samples from identical analysis time points associated with the metagene. The calculation was performed with FSQN normalized data. For each centroid we calculated distances between time points of activation (0.5 to 72 hours) compared to unactivated (0 hours) T-cells. Provided that the analysis time point is present in the *Pan T-cell Verifications Set*, we selected the time point with the maximum absolute distance for filtering genes. The aim was to exclude genes with similar expression in negative control and activated Pan T-cells. We analyzed only genes that are part of the consensus gene signatures and with significant expression changes in negative controls (i.e., with FDR <0.05 in at least one contrast when comparing unactivated Pan T-cells at 6 to 72 hours vs. 0 hours). At the selected time point, i.e., the time point with maximum absolute distance to the centroid of the metagene, we retained genes with absolute “confect” value >1 (FDR <0.05). “Confect” values and their significance between activated and negative controls at this given time point were estimated in the DGEA.

### Metagene landscape

To visualize the relation of sample and gene to metagenes in a two-dimensional space we followed the methodology described previously (61, 62): We first concatenated the metagene expression profile matrices (H) based on temporally coherent metagenes across all CD4+ T-cell subtypes from the *Discovery Set* (the combined matrix *H* in shown in Additional file 1: Figure S19). We calculated a pairwise distance matrix between the metagenes (rows of combined pattern matrix) using Euclidean distance. Then, we projected the distance matrix into two dimensions using the Sammon mapping method from the `MASS R` library (63). The x and y coordinates for each metagene obtained by dimension reduction were standardized. Based the combined pattern matrix H we computed the x and y coordinates for each sample in the two-dimensional space according to:

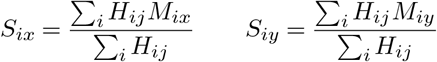

where *M*_*ix*_, *M*_*iy*_ represents the x and y coordinates for metagene *i*. For gene embedding, we used the gene signature matrix of each CD4+ T-cell population. Embedding of genes was performed with the same methodology as for sample embedding. The resulting metagene landscape is shown in Fig. 4A.

### Pathway analysis

We performed enrichment analysis for DE genes with a significant combined effect size and for metagene associated genes using the `Bioconductor` package `clusterProfiler` v3.18.1 (64) on the Gene ontology (GO) database (65) for the ontology biological process and the Reactome database (66). To compare the enrichment of pathways/GO terms between different contrasts and metagenes, we used the `compareCluster` function of the `clusterProfiler` package. For GO analysis, we used the `simplify` function of the `clusterProfiler` with default parameters to remove redundant terms. Significance of enrichment was assessed by a hypergeometric test and adjusted p-values for multiple testing were calculated based on the Benjamini-Hochberg method. All pathways/GO term with FDR *<*0.05 are considered significantly over-represented. To colorize enriched pathways with the uppermost hierarchical level of Reactome database, we used the hierarchical pathway relationship file (ReactomePathwaysRelation.txt, available on www.reactome.org).

### Re-analysis of scRNA-Seq data

We downloaded the raw read counts of the scRNA-Seq study by Deng et al. (27) under the GEO accession number GSE151511. As proposed in Deng et al. only cells with less than 7000 genes and less than 15% of reads mapped to mitochondrial genes were retained. Raw counts were normalized and transformed to log-space using the `NormalizeData()` function implemented in the R package Seurat v4.0.3 [47] with default settings. Only cells with expression of CD3D or CD3E or CD3G >1 normalized count value were used for subsequent analysis. In addition, we filtered for CD8-CD4+ (15936 cells) and CD8+CD4- (59002 cells) CAR T-cells based on the gene expression data using following strategy: Given the normalized expression, one cell was considered as CD8 positive or negative if the normalized count value of CD8A or CD8B was *>*1 or ≤1, respectively. One cell was considered as CD4 positive or negative if the normalized count value of CD4 was *>*1 or ≤1, respectively.

We detected highly variable genes using the “vst” method of the `FindVariableFeatures()` function in `Seurat` at default settings, resulting in 3000 highly variable genes. Principal component analysis (PCA) was conducted using standardized and normalized highly variable genes with the function `RunPCA` implemented in `Seurat`. We used 39 principal components, which explained 95% of the variance, to integrate the dataset from different samples using `RunHarmony()` from the Harmony v1.0 `R`package (67). Using the Harmony-corrected cell embeddings, we computed a shared nearest neighbors graph, as implemented in `FindNeighbors()` function in `Seurat` with default settings.Cluster identification using the SNN graph was computed by the `FindCluster` function in `Seurat` with default setting. The Harmony-corrected cell embeddings were projected into a two-dimensional space using the t-distributed stochastic neighbor embedding (tSNE) method. For this, we used 30 Harmony-corrected components, which explained 95% of the variance.

Known T-cell state marker genes (68, 69) including 5 exhausted genes (CTLA4, HAVCR2, LAG3, PDCD1, TIGIT), 12 cytotoxic genes (PRF1, IFNG, GNLY, NKG7, GZMB, GZMA, GZMH, KLRK1, KLRB1, KLRD1, CTSW, CST7) and 5 naïve/memory genes (CCR7, TCF7, LEF1, SELL, CD44) were used to calculate the standardized average expression in each cluster. We also used gene signatures from Azizi et al. (28) to associate cell clusters with a functional context.

For each metagene, T-cell state marker and gene signatures (referred to as gene sets), we conducted a permutation test. We generated a null distribution for each gene set by performing 1000 permutations. For each permutation, a random gene set was formed. Random gene sets are not part of genes associated with the original gene sets and have the same number of genes as the corresponding original gene set (based on 3000 present highly variable genes). For each random gene set and patient, we calculated the aggregate expression (summed average expression) of CD8+CD4- and CD8-CD4+ cells. We then computed the log2 fold change in the median of aggregate expression between low- and high-grade ICANS patient groups. Based on the log2 fold change between low- and high-grade ICANS from the original gene set, we calculated the statistical significance using a left-tailed test for positives log2 fold changes or a right-tailed test for negative log2 fold changes.

## Ethics approval and consent to participate

For experiments using human cells (PBMCs) all of the Patients or their next of kin, caretakers, or guardians gave written informed consent to the Blutspendedienst NSTOB. Fraunhofer received only anonymized samples. All samples were received between 2017 and 2018.

## Consent for publication

Not applicable.

## Availability of data and materials

The RNA-Seq and microarray datasets analyzed in this study are available in the NCBI GEO database (https://www.ncbi.nlm.nih.gov/geo/) with accession numbers GSE52260 (12), GSE90569 (14), GSE94396 (13), GSE96538 (13), GSE17974 (10), GSE32959 (11), and GSE140244 (15). Raw and normalized gene expression counts for the *Pan T-cell Verifications Set* are available from GSE197067 (accessible to the public when the manuscript is published). In Additional file 2 we summarize all genes, including interactive visualization of expression profiles of individual genes of the consensus signatures.

## Authors’ contributions

KR, KS, and MR designed the study. SB, SD, and VN performed sample collection, T-cell isolation, and activation from the *Pan T-cell Verifications Set*. CB and DL performed sample preparation and transcriptome-wide sequencing of the *Pan T-cell Verifications Set*. MR performed data analysis and wrote the manuscript. KR revised the manuscript and supervised the study. All authors were responsible for reviewing and editing the final version of the paper. All authors read and approved the final manuscript.

