## Additional File 1 for "A time-resolved meta-analysis of consensus gene expression profiles during human T-cell activation"

**1 Data sources** 1
- 1.1 Transcriptome datasets used for the development of the consensus gene signature . . 1

S27 *Pan T-cell Verification Set*: Temporal expression pattern of genes from the consensus signature that passed the two *Verification Sets* but not the negative control kinetics. 42
S28 Consensus gene expression profiles verified for temporal consistency by both *Verification Sets* (activation and negative control kinetics) . . . 44
S29 Reactome enrichment analysis for genes from the consensus gene expression profiles that passed both *Verification Sets* . . . 44

S30 Standardized average expression of known T-cell state/molecular mechanisms markers for each cluster . . . 45
S31 Aggregated expression between patients with low- and high-grade ICANS for known T-cell state/molecular mechanisms markers . . . 46
S32 otential batch effects between aggregated expression and patient characteristics . . . 47
S33 Test for differences in aggregate expression between patients with low- and high-grade CRS or clinical outcome (CR compared to PR/PD) . . . 47

### List of Tables

S1 Overview of the datasets included for discovery and verification . . . 2
S2 References used for FastQ Screen . . . 7

### 1 Data sources

#### 1.1 Transcriptome datasets used for the development of the consensus gene signature

Table S1 and Figure S1 provide an overview of the datasets used for the discovery and verification of temporal consensus gene expression signatures of T cells.

#### 1.2 *Pan T-cell Verification Set*

##### 1.2.1 Preparation of peripheral mononuclear blood cells (PBMCs) and pan T-cell isolation

Buffy coats were obtained from the Springe Blutspendedienst and were kept at 4 °C until PBMC isolation. PBMCs were isolated by density centrifugation. The cell count and viability was determined with a hemocytometer and trypan blue staining. CD14+ monocytes, CD19+ B-cells and CD56+ natural killer cells were depleted using CD14, CD19 and CD56 magnetic cell sorting (MACS) beads (Miltenyi) following the manufactures instruction. The flow through was kept for CD2+ T-cell isolation. Therefore $1x10^7$ cells were resuspended and CD2 microbeads were added for 15 min at 4 °C After centrifugation (10 min, 300 xg, 4 °C) and resuspension of the cells in MACS-buffer. LS columns (Miltenyi) were placed into a magnet and cells were loaded onto the column through a pre-seperation filter (Miltenyi). The cell count was determined by trypan blue staining using a hemocytometer. Quality of the CD2+ isolation was confirmed by fluorescence-activated cell sorting (FACS).Isolated pan T-cells ($0.5\text{-}1\times10^6$) were incubated with 2.5 µL human TruStain FcXTM-block for 10 min at 4 °C. Viable cells were stained with Zombie NIRTM for 15 min at RT. Afterwards, cells were stained with CD3-FITC, CD45-AF700, CD-BV510, CD4-PacificBlue, Cd45RO-PE/Cy7, CD45RA-APC, CD14PerpCy/Cy5 (BioLegend, San Diego, USA) or respective isotype controls for 30 min on ice. Zombie NIRTM was used as a viability marker due to manufactures instructions. Samples were washed with 2 mL FACS buffer (PBS supplemented with 0.25% bovine serum albumin and 2 mM ethylenediaminetetraaceticacid (EDTA)). After centrifugation at 350 xg for 10 min, cells were resuspended. Viable leukocytes were discriminated from cell debris by forward (FSC) and sideward scatter (SSC) properties as well as CD45+, Zombie NIR+ cells Cells were separated by size in FS (Forward Scatter) and by granularity in SS (Sidewards Scatter). Subsequently, duplicates of singular cells were eliminated via the disproportion of height (PEAK) and width (TOF). The singular leukocytes negative for the dead dye ZombieNIR were further differentiated into monocytes (CD14) and CD3 T-cells. NK cells and B cells were identified on CD3- cells using CD56 and CD19. CD3+ cells were further differentiated into T helper CD4+ and cytotoxic CD8+ cells. Naive T-cells were discriminated as CD45RA+ cells and memory T-cells as CD45RO+ cells.

For quality control, 200.000 cells were analyzed using a Beckman Coulter Navios 3 L 10 C. Flow cytometry was analyzed using Kaluza 2.1. software. Analysis were done using seven human blood donors. Pan T-cell isolation strategy resulted in a CD3+ T-cell purity of 97,7 ± 1,8%. Fractions of the T-cell populations are shown in Figure 5B (main part) Stimulation of the T-cells with either CD3/CD28 beads to activate T-cells via CD3 and CD28 receptors, Phorbol Myristate Acetate (PMA) to activate protein kinase C together with ionomycin, a calcium ionophore or Phytohämagglutinin (PHA) a mitogen resulted in proliferation and cytokine (IL-2) secretion of the cells. Viability was proven in ATP assay. Controls stimulation assays were performed with LPS and medium (see Figure S2).

##### 1.2.2 RNA-seq sample preparation and sequencing

For RNA sequencing, up to $0.5x10^6$ treated or untreated T-cells per sample were collected. The cell pellets were re-suspended in Qiazol and stored at –80°C. RNA was extracted according to the miRNeasy mini protocol (Qiagen). After two steps of DNase-digestion (TURBO DNA free

Tab. S1: Overview of datasets included for discovery and verification. Green colored time points were used for the meta-analysis.

Discovery set

| Study | Organisation | PMID | GEO | Layout | Reads (bp) | Platform | Strandness | Extraction | Sample source | All\|Bio\|Tech Repl. | 0h naive | 0.5h | 1h | 2h | 4h | 6h | 8h | 12h | 24h | 2d | 3d | 4d | 5d | 6d | induced to following phenotye: | |
|---|---|---|---|---|---|---|---|---|---|---|---|---|---|---|---|---|---|---|---|---|---|---|---|---|---|---|
| Tuomela et al. | Aalto University | 26967054 | GSE52260 | SE | 1x50 | HiSeq 2000 | N | total RNA | Cord blood | 51\|3\|1 | Y | Y | Y | Y | Y | Y | | Y | Y | Y | Y | | | | Th0 | Th17 |
| Ullah et al. | Aalto University | 29466736 | GSE90569 | PE | 2x101 | HiSeq 2000 | N | polyA RNA | Cord blood | 51\|3\|1 | Y | Y | Y | Y | Y | Y | | Y | Y | Y | Y | | | | Th0 | iTreg |
| Schmidt-ind | Aalto University | 29730990 | GSE96538 | PE | 2x101 | HiSeq 2500 | Y | polyA RNA | PBMC | 51\|3\|2* | Y | Y | Y | Y | | Y | | Y | Y | Y | Y | Y | Y | | Th0 | iTreg |
| Schmidt et al. | Karolinska Institut | 29730990 | GSE94396 | PE | 2x75 | HiSeq 2500 | N | total RNA | PBMC | 27\|3\|1 | Y | | | Y | | Y | | | Y | Y | | | | Y | Th0 | iTreg |
| Elo et al. | University of Turku | 20620947 | GSE17974 | | | Affymetrix HG-U133 Plus 2.0 | | | Cord blood | 49\|3\|1 | Y | Y | Y | Y | Y | Y | | Y | Y | Y | Y | | | | Th0 | Th2 |
| Äijö et al. | University of Turku | 23110343 | GSE32959 | | | Affymetrix HG-U133 Plus 2.0 | | | Cord blood | 27\|3**\|1 | Y | | | | | | | Y | Y | Y | Y | | | | | Th1 |

SE = single-end; PE = paired-end; Th0 = activated CD4+ T-cells without differenation cytokines; Bio Repl. = Biological replicates; Tech Repl. = Technical replicates; * For 2, 6 and 12h only one techn. replicate is available: ** For 72h only two biological replicate

GSE52260: Only human data were analyzed. Five samples were sequenced with 2x75 bp paired-end reads. As in Tuomela et al. those reads were truncated to 50 bp and only one of the paired-end reads was used.

GSE94396 : G01 (unstimulated), G02 (Th0) and G04 (iTreg) datasets were used. To be consistent with the treatment protocols for iTreg differentiation across datasets, we used only samples induced with TGF-β + ATRA reagents.

GSE17974: unstimulated and induces Th2 samples were analyzed

GSE32959: unstimulated and induces Th1 samples were analyzed. **Note**: The same pooled cells as for the Th2 (IL-4 stimulated) culture conditions by Elo et al. (GSE17974) were used.

Verification set

| Study | Organisation | PMID | GEO | Layout | Reads (bp) | Platform | Strandness | Extraction | Sample source | All\|Bio\|Tech Repl. | 0h | 0.5h | 1h | 2h | 4h | 6h | 8h | 12h | 24h | 2d | 3d | 4d | 5d | 6d | induced to following phenotye: |
|---|---|---|---|---|---|---|---|---|---|---|---|---|---|---|---|---|---|---|---|---|---|---|---|---|---|
| Rade et al. | Fraunhofer ITEM/IZI | | GSE197067 | PE | 2x101 | HiSeq 2000 | Y | total RNA | PBMC | 44\|4\|1 | Y | | | | | Y | | Y | Y | Y | Y | | | | Th0 & negative controls |
| Gutierrez-Arcelus et al. | Brigham and Women's Hospital, Harvard Medical School | 32066938 | GSE140244 | PE | 2x101 | HiSeq 2000 | Y | polyA RNA | PBMC | 184\|24\|1 | Y | | | Y | Y | | Y | Y | Y | Y | Y | | | | Th0 |

GSE140244: Only replicate A was used (2 biological replicates have technical replicates)

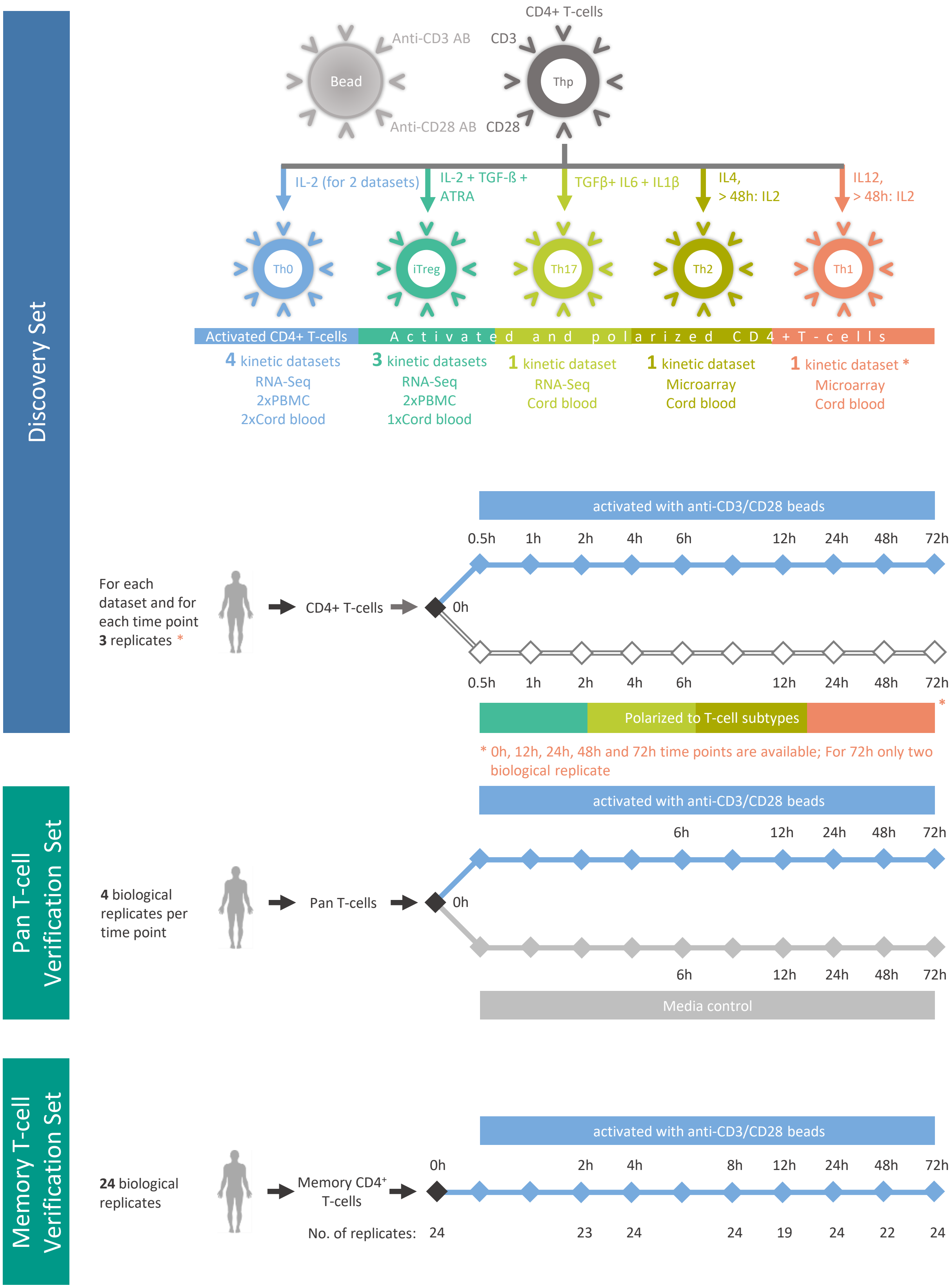

Fig. S1: Overview of the datasets included for discovery and verification.

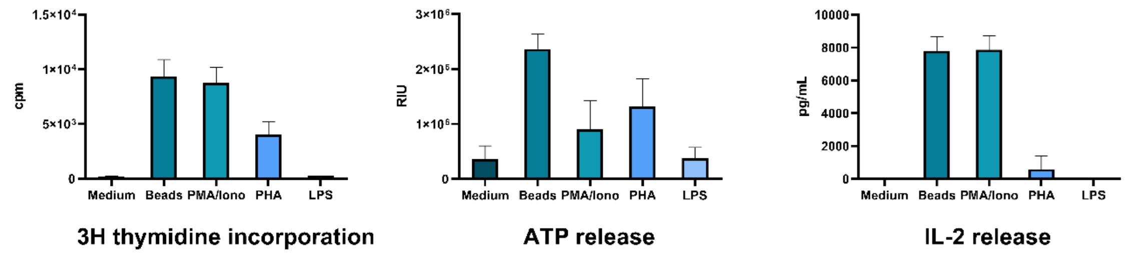

Fig. S2: ***Pan T-cell Verification Set*: Biological activity of isolated T-cells.** Proliferation capacity, viability and cytokine secretion was analyzed after three days stimulation with either CD3/28 receptor activating beads, PMA/Ionomycin or PHA using ATP assay, 3H thymidine proliferation assay and IL-2 ELISA. Controls stimulation assays were performed with LPS and medium. Assays were done using T cell isolations from three human blood donors in technical triplicates. Analysis were performed using GraphPad Prism 9.0.

Kit, Ambion), extracted RNA was quantified using a Qubit RNA-Kit and the DeNovix instrument (Biozym). RNA quality was analyzed on a Bioanalyzer 2100 instrument (Agilent Technologies). For subsequent RNA-sequencing analyses, 150 ng of total RNA per sample were used. Library preparation was conducted using Truseq-Stranded total RNA Sample Prep kit (Illumina, Inc, San Diego, CA) according to the manufacturers' protocol. Molarity of each library was calculated and equal amounts were pooled and used for sequencing (12 pM). Sequencing was performed with 2 $\times$ 126-bp paired-end reads using HiSeq SBS Kit v4 chemistry on a HiSeq 2500 instrument (Illumina). Three flowcell à 8 lanes were sequenced with all 44-pooled libraries each.

### 2 Pre-processing

#### 2.1 Microarray datasets

Each T-cell population dataset was processed separately as follows: normalization and probeset summarization was performed using the robust multi-array average algorithm (RMA) implemented in the `rma()` function of the `affy` v1.68.0 `R` package [1]. An updated probeset annotation chip definition file (CDF) based on GENCODE v29 and provided by BrainArray [2] (version 23.0.0 for Affymetrix HGU133Plus2 arrays) was used (File available at `http://brainarray.mbni.med.umich.edu/Brainarray/Database/CustomCDF/23.0.0/gencodeg.asp`). Each sample was evaluated by visual inspection of the array pseudo-images, density plots of probe intensities, Normalized Unscaled Standard Error (NUSE) and Relative Log Expression (RLE) implemented in the `affyPLM` v1.66.0 `R` package [3] (data not shown). An unspecific gene filtering step was incorporated into the analysis. The normalized intensities for at least 3 samples, which represent the smallest experimental group, must be larger than the 25th percentile across all gene intensities. Furthermore, low-intensity probes were filtered out based on interquartile range (IQR) as described in Chockalingam et al. [4]. The principal component analysis did not reveal any critical outlier (see Figure S5D-E).

#### 2.2 RNA-Seq datasets

Demultiplexing of Illumina raw files from the *Pan T-cell Verification Set* was performed with the Illumina `bcl2fastq` software v2.19 [5]. The following steps were performed for all RNA-Seq datasets from the *Discovery Set* and *Pan T-cell Verification Set*: The single-end or paired-end FASTQ reads were trimmed and filtered using `AdaptorRemoval` v2.3.1 [6] with additional parameters to trim ambiguous bases (N) at 5'/3' termini ( –trimns), remove low-quality bases (–trimqualities, –minquality 20) and keep reads with a minimum read length of 30bp (–minlength 30). Trimmed reads were mapped to the human reference genome version GRCh38/hg38 by `HISAT2` v2.1.0 [7]. Gene level quantification for the human reference gene annotation GENCODE (release 29 GRCh38.p12) was obtained by using HTSeq v0.11.2 [8]. Sample QC was reported using FastQC v0.11.8 [9] to assess base call accuracy, Preseq v2.0.3 [10] to evaluate the library complexity. Duplication metrics were collected using Picard tools v2.18.29 (http://broadinstitute.github.io/picard/) function MarkDuplicates using BAM files generated by HISAT2. Picard's CollectRnaSeqMetrics was used to collect mapping percentages on intergenic, intronic, coding and UTR regions as well as gene body coverage. RSeQC v3.0.0 [11], was used to determine, read GC content, junction saturation, read pair inner distance, and strandness of reads. FastQScreen v0.14.0 [12] in conjunction with bowtie2 [13] was conducted to assess RNA library composition. Aggregated data visualization for the secondary analysis and quality control were generated using the MultiQC [14] framework. A quality control summary report for the *Pan T-cell Verification Set* is shown in Figure S3. Two samples mapped with at least 25% against the human rRNA reference genome (Figure S3C). However, we did not observe a noteworthy difference of the number of uniquely mapping reads against the human genome reads or the number of reads assign to genes (Figure S3D-F) compared to the other samples. Principal component analysis of all samples did not reveal any critical outliers (Figure S3F).

Gene counts generated by HTSeq from different studies and based on the same $CD4^+$ T-cell population were concatenated. For each population gene counts were converted to count-per-million (CPM). Low expressed genes were filtered out from downstream analysis using the `filterByExpr()` function from the edgeR v3.32.1 R/Bioconductor package [15] with default settings. Genes having a CPM above $c$ in $m$ samples were kept, where $m$ the size of the smallest experimental group and $c$ is determined by 10 counts divided by the median library sizes times one million. Normalization factors to scale the raw library size were calculated using the function calcNormFactors() from edgeR with the Trimmed mean of M-Values (TMM) method. [16]. For T-cell populations consisting of multiple studies (Th0 and iTreg), log-transformed CPM expression data was adjusted for the factor *study* using the `removeBatchEffect()` function in the R/Bioconductor limma v3.46.0 package [17, 18]. After batch correction no significant correlation was observed between the first 50 principal components and potential confounding factors such as *study* or *sample source* (PBMCs or neonatal cord blood) (Figure S4). For principal component analysis of the T-cell populations see Figure S5. The batch corrected CPM expression data was used as input for non-negative matrix factorization.

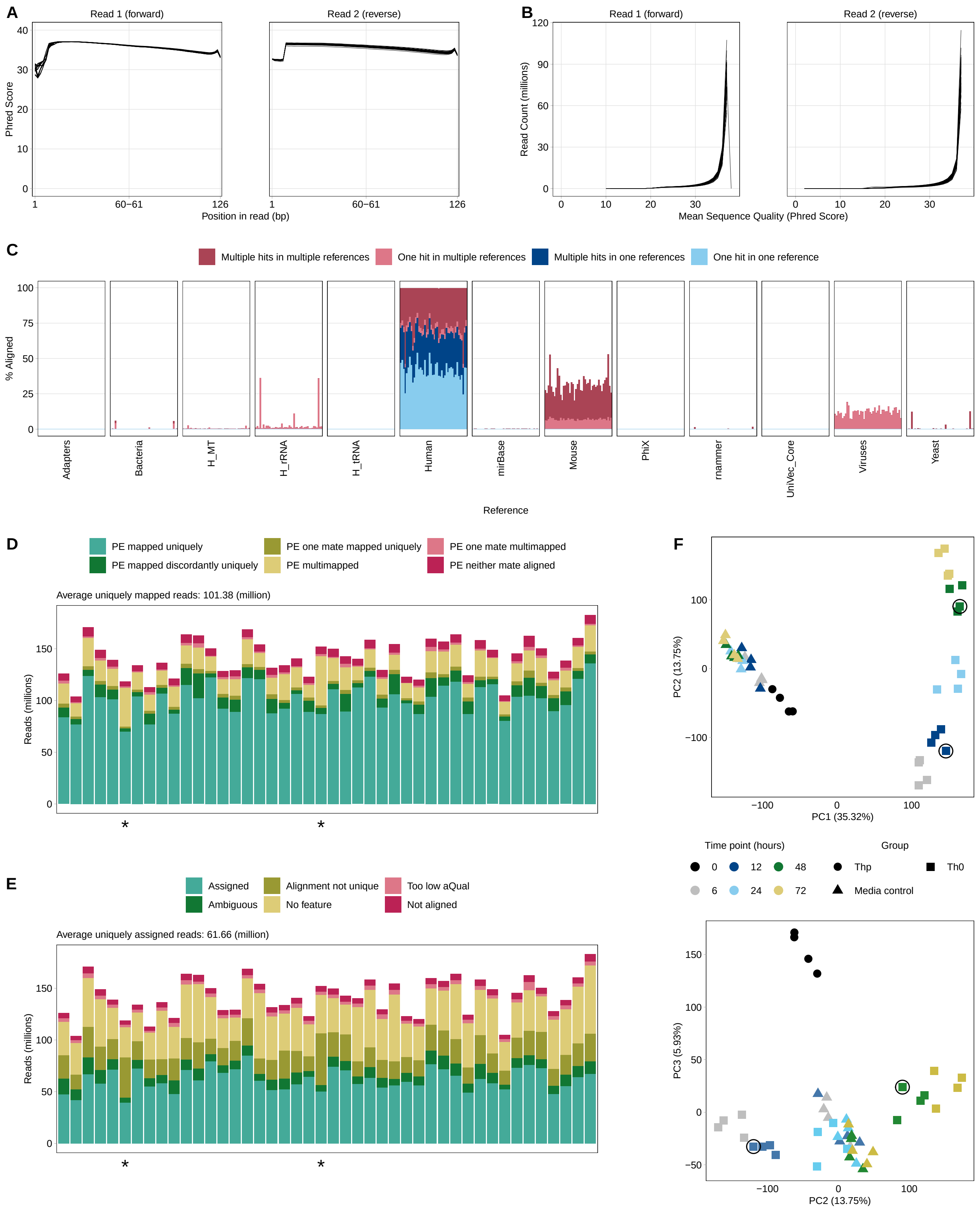
A
Read 1 (forward)
Read 2 (reverse)
Phred Score
Position in read (bp)
B
Read 1 (forward)
Read 2 (reverse)
Read Count (millions)
Mean Sequence Quality (Phred Score)
C
Multiple hits in multiple references
One hit in multiple references
Multiple hits in one references
One hit in one reference
% Aligned
Adapters
Bacteria
H_MT
H_rRNA
H_tRNA
Human
mirBase
Mouse
PhiX
rnammer
UniVec_Core
Viruses
Yeast
Reference
D
PE mapped uniquely
PE mapped discordantly uniquely
PE one mate mapped uniquely
PE multimapped
PE one mate multimapped
PE neither mate aligned
Average uniquely mapped reads: 101.38 (million)
Reads (millions)
E
Assigned
Ambiguous
Alignment not unique
No feature
Too low aQual
Not aligned
Average uniquely assigned reads: 61.66 (million)
Reads (millions)
F
PC2 (13.75%)
PC1 (35.32%)
Time point (hours)
0
12
48
6
24
72
Group
Thp
Th0
Media control
PC3 (5.93%)
PC2 (13.75%)

Fig. S3: ***Pan T-cell Verification Set*: Quality Control of samples with respect to sequencing library composition, read alignment and sample variance**. Following adapter trimming, each FASTQ file were assessed for average per base **(A)** and per sequence **(B)** quality as measured by Phred score. **(C)** To assess the sequencing library composition, each sample was subsampled to randomly 1 million trimmed paired-end reads. `FastQ Screen` in conjunction with `bowtie2` was conducted to detect possible contamination like for example bacteria and overrepresented fractions of RNA species like human rRNA. The y-axis depicts the percentage of first reads for each sample that aligned against the references from Table S2. Reads are classified into four distinct types indicating reads uniquely mapping in one sequence reference (one hit in one reference), reads with multiple mappings in one sequence reference (multiple hits in one reference), reads uniquely mapping in distinct sequence references (one hit in multiple references) and reads with multiple mappings in distinct sequence databases (multiple hits in multiple references). **(D)** Quality assessment of read alignment results for each sample. Mapped uniquely: read pairs aligned concordant 1 time. Multi-mapped: read pairs aligned concordant >1 times. Mapped discordantly uniquely: read pairs aligned discordantly 1 time. One mate mapped uniquely: one read of the pair maps to the genome 1 time. One mate multi-mapped: one read of the pair maps to the genome >1 times. **(E)** Results of gene quantification for each sample. Quantification was performed with `HTSeq`. For **(D)** and **(E)**: Samples mapped with at least 25% against the human rRNA reference depicted with asterisks. **(F)** Principal component analysis of variance-stabilized counts based on the 5000 most variable genes. The upper plot depicts first and second principal components, the bottom plot the second and third principal component. Samples are shaped as asterisks, if the subsampled reads mapped with at least 25% against the human rRNA reference.

Tab. S2: References used for FastQ Screen

| **Reference** | **Source** |
|---|---|
| Adapter sequences | https://github.com/csf-ngs/fastqc/blob/master/Contaminants/contaminant_list.txt |
| Bacteria | ftp://ftp.ncbi.nih.gov/genomes/refseq/bacteria/, Oct 2014 |
| H_MT | human mitochondrial reference sequence from GRCh37/hg38 |
| H_rRNA (human ribosomal RNA sequences) | NR_003286.1 (18S), NT_003287.1 (28S), NR_003285.2 (5.8S), V00589.1 (5S), NC_012920.1: gi\|251831106:1671-3229 (MT 16S) and NC_012920.1: gi\|251831106:648-1601 (MT 12S) |
| H_tRNA (human transfer RNA sequences) | http://gtrnadb.ucsc.edu/genomes/eukaryota/Hsapi38/hg38-tRNAs.fa |
| Human genome | GRCh38/hg38, reference chromosomes only |
| mirBase | miRNA sequences from mirBase v21 |
| Mouse genome | UCSC/mm10 |
| PhiX | gi\|9626q372\|ref\|NC_001422.1\| Enterobacteria phage phiX174 sensu lato, complete genome |
| RNAmmer (predicted rRNA sequences) | http://www.cbs.dtu.dk/services/RNAmmer/, v1.2 |
| UniVec Core | ftp://ftp.ncbi.nlm.nih.gov/pub/UniVec/UniVec_Core build 8.0, May 2015 |
| Viruses | ftp://ftp.ncbi.nlm.nih.gov/genomes/refseq/viral/, March 2014 |
| Yeast | Genome assembly SacCer3 |

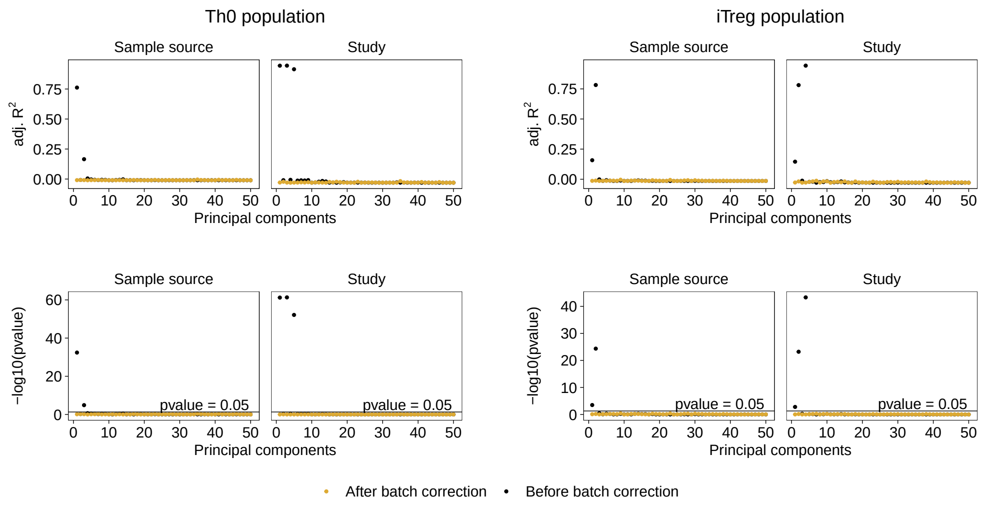

Fig. S4: ***Discovery Set*: Correlation analysis of potential confounding factors.** Analyses was performed for T-cell populations (Th0 and iTreg) consisting of multiple studies. Correlation and F-statistic based on linear regression with the factors *sample source* (PBMCs or cord blood) or *study* as covariate for the first 50 principal components. The 5000 most variable genes were used as input to PCA. Linear regression was performed before and after batch correction by adjusting for the factor *study* using the `removeBatchEffect()` function from the `limma` package. The top panels depicted the adjusted coefficient of determination and the bottom panels the p-values from the F-stastic calculated by the lm() function in `R`. After batch correction no significantly (p-value $< 0.05$) correlation could be observed between the factors *sample source* or *study* and the first 50 PCs.

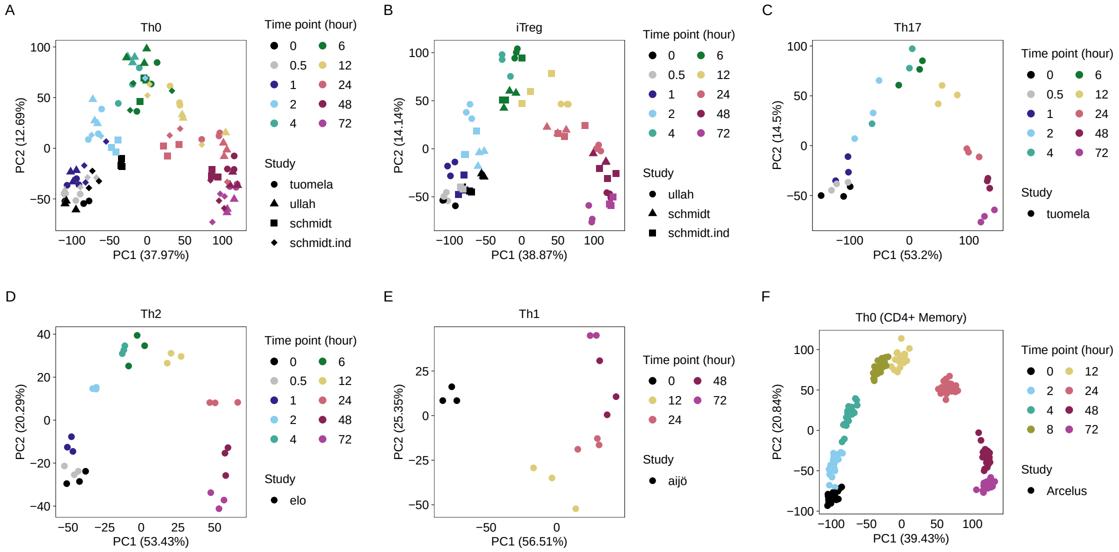

Fig. S5: ***Discovery and Verification Sets*: Principal component analysis (PCA).** PCA of time-course RNA-Seq and microarray datasets was performed separately for each T-cell population from the *Discovery Set* and *Verification Sets*. The 5000 most variable genes were used for PCA. For RNA-Seq data, we used log-transformed counts per million (CPM) values as input to PCA. In case of microarrays, we used the log-transformed normalized intensities using the multi-array average (RMA) normalization method. For T-cell populations (Th0 and iTreg) consisting of multiple studies, potential batch effects across the studies were removed using the `removeBatchEffect()` function from the `limma` package. The plot depicts the first and second principal components with samples colored according time point of activation and shaped by study. Plot **(A-E)** depict samples from the *Discovery Set* and plot **(F)** from the *Memory T-cell Verification Set* . A PCA plot from the *Pan T-cell Verification Set* is shown in Figure S3F. PCA of all samples did not reveal any critical outliers.

#### 3 Differential gene expression analysis

As input to `limma` we used for RNA-Seq data the filtered gene counts (see subsection 2.2) and the normalized factors from the TMM method. In case of Microarrays we used the filtered normalized intensities from the RMA method (see subsection 2.1). For the T-cell populations Th1, Th2, Th17 from the *Discovery Set* and Th0 from both *Verifications Sets* a paired design of the form $\sim 0 + block + contrast$ was used, where *block* encodes the replicates and *contrast* encodes the pre- and post-treatment information for each time point. For the T-cell populations Th0 and iTreg from the *Discovery Set*, *block* encodes the different studies due to unbalanced paired design. In case of RNA-Seq data, the `voom` [19] function of `limma` was used to accommodated the mean-variance relationship using precision weights. Linear modelling and empirical Bayes moderation to assess differential expression was performed using the `lmFit()` and `contrasts.fit()` function from `limma`. In addition, the `limma_confects()` function of the `Topconfects` v1.6.0 `R/Bioconductor` package [20] was used. This function calls the function `eBayes()` from `limma` itself and builds on the `TREAT` method [21]. `Topconfects` calculates for each gene a confident effect size or "confect", a confident inner bound of the calculated effect size by `limma` while maintaining a given false discovery rate (FDR). "Confects" in this case can be considered as moderated log2 fold changes and are used to rank significant differentially expressed (DE) genes which is a more conservative method than ranking by raw log2 fold changes. A gene was considered as DE if the FDR-adjusted p-value was <0.05. Volcano plot representation of differential expression analysis is shown in Figure S6 An overview of DE genes from the *Discovery Set* genes is shown in Figure S7.

We observed a greater difference in the number of DE genes between Th1/Th2 compared to the other T-cell populations, especially after 2 hours of activation (see blue vertical bars in Figure S7). Th1 and Th2 are based on gene expression arrays, while Th0, Th17 and iTreg are RNA-Seq data. For gene expression arrays 21302 Ensembl gene ID from GENCODE could be mapped to microarray propes and evaluated, whereas for RNA-Seq 58721 Ensembl gene ID were evaluated. This could explain the difference in the number of DE genes between the populations.

#### 3.1 Volcano plots

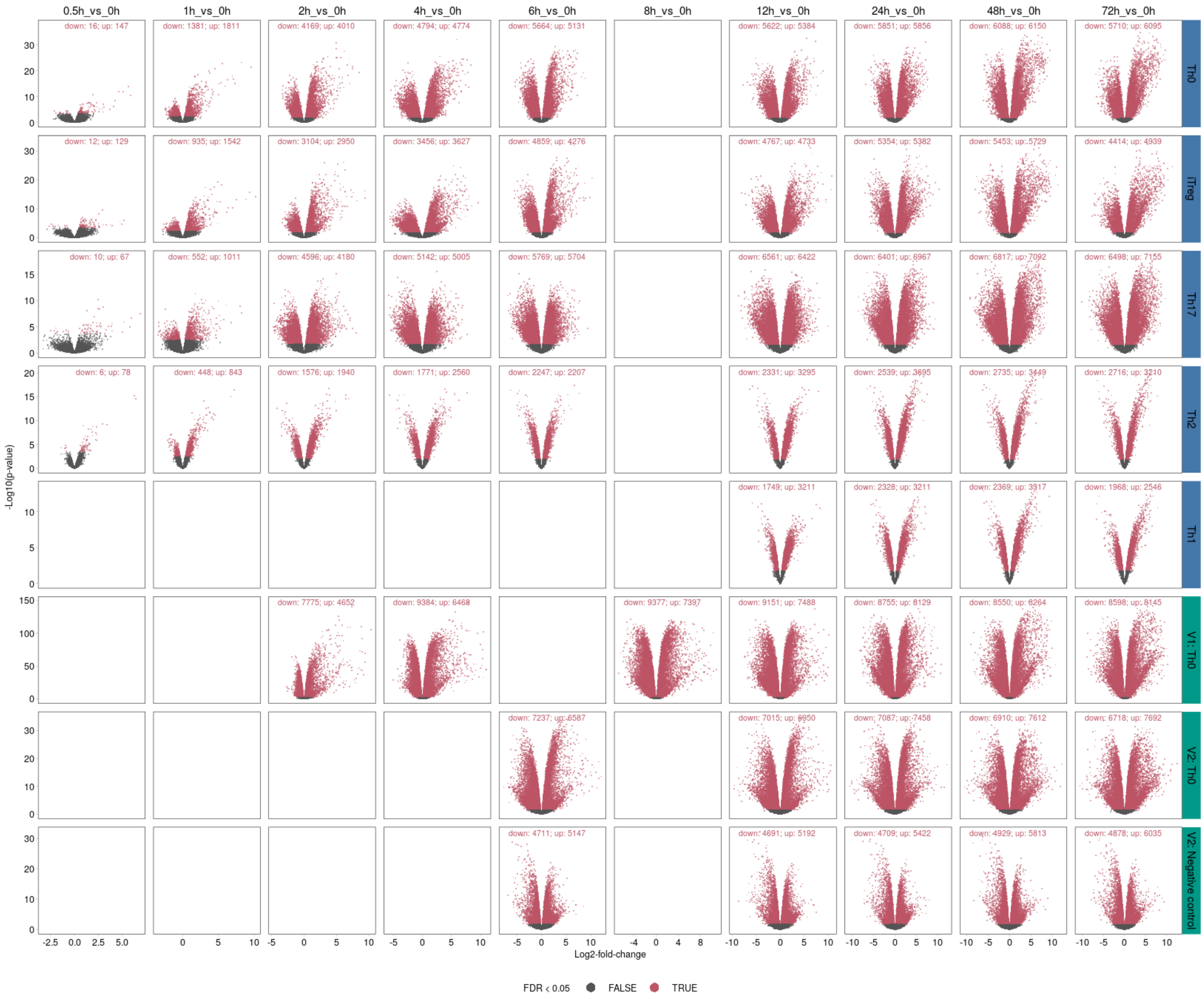

Fig. S6: ***Discovery and Verification Sets*: Volcano plots of DE genes for each contrast and T-cell population.** Red dots represent DE genes (FDR <0.05). The number of down- and up-regulated DE genes are shown at the top of each volcano plot. The y-axis denotes -log10 p-values while the x-axis represents log2 fold change values. Labels with blue color indicate the *Discovery Set*. Labels with green color indicate the *Verification Sets* (V1 = *Memory T-cell Verification Set*, V2 = *Pan T-cell Verification Set*.

#### 3.2 *Discovery Set*: UpSet plots of DE genes

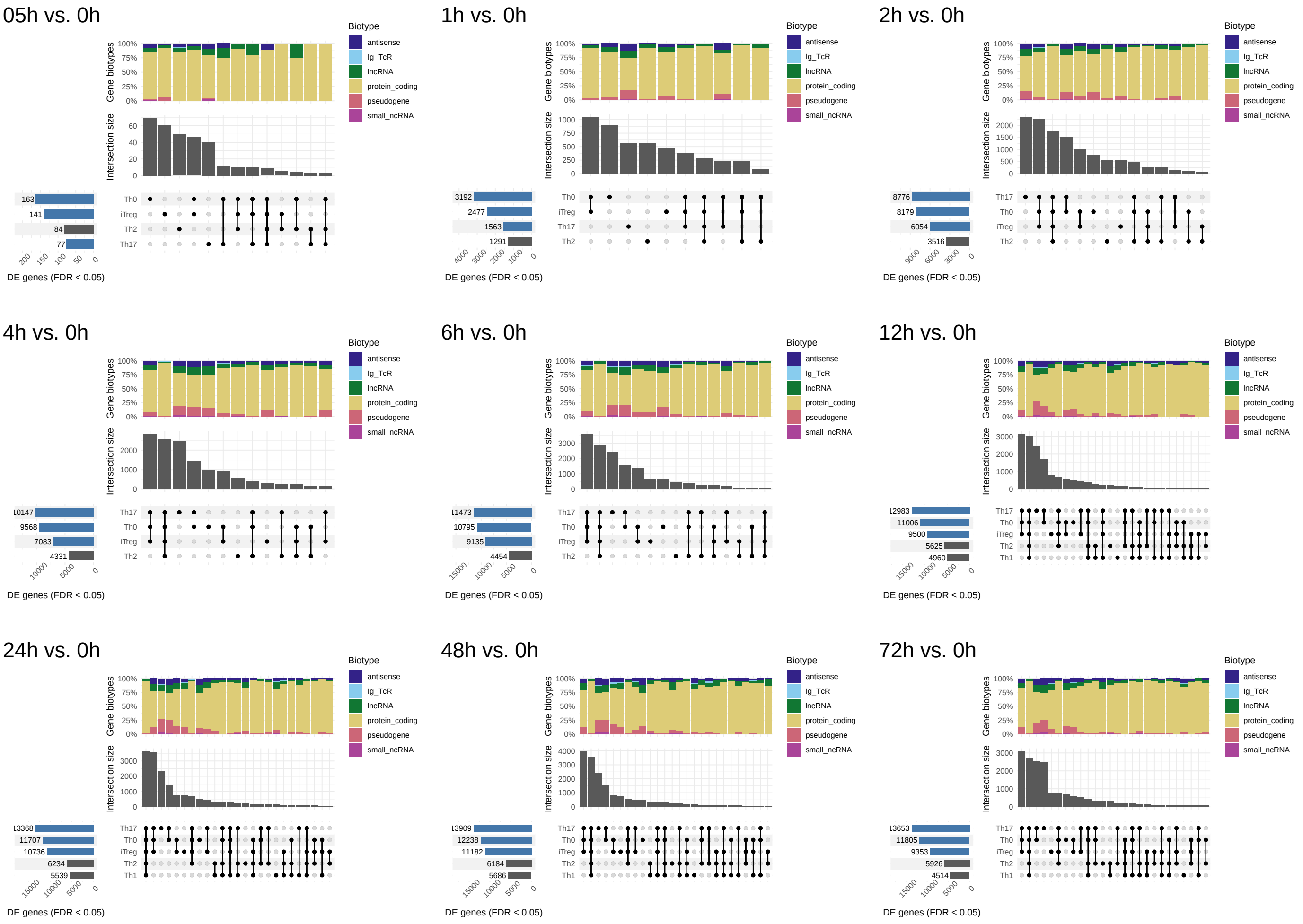

Fig. S7: ***Discovery Set*: Overview of DE genes for all CD4$^{+}$ T-cell populations** UpSet plots summarizes all DE genes (FDR <0.05) of activated CD4$^{+}$ T-cell populations compared to unactivated (0h) CD4$^{+}$ T-cell populations for each analyzed time point. In each panel, the bottom left horizontal bar graph indicates the total DE genes for each T-cell population. The blue horizontal bar graph indicate datasets from RNA-seq. The circles in each matrix represent unique and overlapping DE genes for the T-cell populations. Connected circles indicate a certain intersection of DE genes between the populations. The bar graph above the matrix in each panel summarizes the number of DE genes for each unique or overlapping combination. The top stacked barplot depicting the fraction of gene types (biotypes) for each unique or overlapping combination. With exception to the contrast "05h vs. 0h" the minimal number of observations in an intersection is set to 50

#### 4 Meta-analysis

The calculation of the Hedges' g values [22, 23, 24] involves the calculation of the Cohen's d value:

$$\text{Cohen's } d_{ij} = \frac{\overline{x}_{ijPost} - \overline{x}_{ijPre}}{s_{ij}} \tag{1}$$

where $x_{ijPost}$ and $x_{ijPre}$ is the mean expression for each DE gene $i$ of the post- and pre-treatment group in population $j$. Here we used the log2 fold changes as estimated in `limma`. For the pooled standard deviation $s_{ij}$ we adopted the definition of the pooled variance as in the `limma` procedure, by borrowing information of variances from all tested genes, more specific, the square root of the posterior estimate of the variance as calculated by the empirical Bayes method from the eBayes() function. Since the Cohen's d value overestimates the effect size for studies with small sample sizes following correction factor was incorporated:

$$\text{Correction factor } J_{ij} = 1 - \frac{3}{4(n_{jPost} + n_{jPre}) - 9} \tag{2}$$

where $n_{jPost}$ and $n_{jPre}$ is the number of samples for the post- and pre-treatment group in the T-cell population $j$. The Hedges' g value for each DE gene can be computed as:

$$\text{Hedges' } g_{ij} = d_{ij} \times J_{ij} \tag{3}$$

The variance of the Hedges' g value was calculated by:

$$\text{V}g_{ij} = J_{ij}^2 \times (\frac{n_{jPost} + n_{jPre}}{n_{jPost} n_{jPre}} + \frac{d_{ij}}{2(n_{jPost} + n_{jPre})}) \tag{4}$$

Using the estimated Hedges'g values and the corresponding variances of the Hedges'g values, a meta-analysis with a random effects model was then performed. We analyzed each DE gene in at least 2 T-cell populations and time points of the *Discovery Set*. To assess the amount of heterogeneity post-hoc, the $I^2$ statistic was calculated. We observed increased heterogeneity ($I^2 > 75\%$) for 1548 genes with a significant combined effect size (FDR <0.05) across all 5 T-cell populations (Figure S8B). Considering the estimated standard deviation of log2 fold changes from the differential gene expression analysis, we observed that the estimated standard deviations are lower for the microarrays compared to the RNA-Seq datasets (Figure S9A). This results in higher Hedges'g values (equation 1 and 3) and consequently to higher estimated variances (equation 4 and Figure S9B). One explanation could be a wider dynamic range of the gene expression measurements between platforms. Gene expression intensities for microarrays are limited by background at the low end and signal saturation at the high end. On the other hand, RNA-Seq platforms provide discrete read counts and can quantify gene expression across a wider dynamic range, as it is determined only by the amount of sequencing obtained [25]. This larger dynamic range can lead to a higher variance of the read counts between replicates.

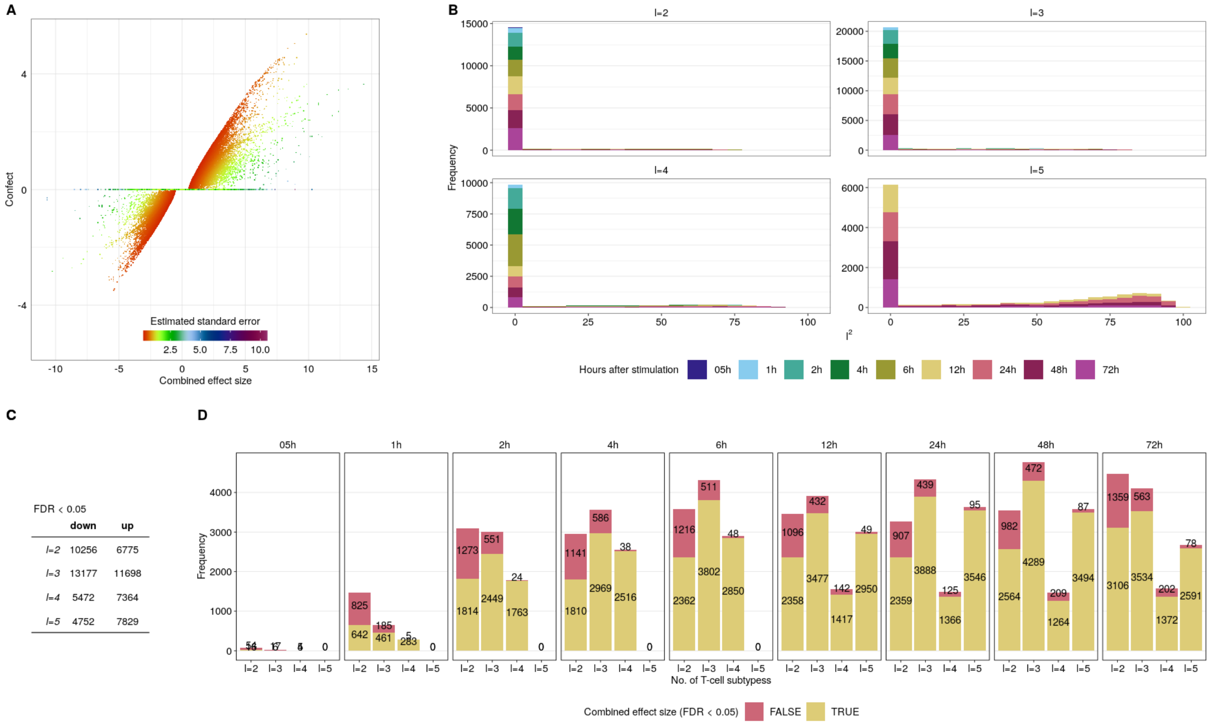

Fig. S8: ***Discovery Set*: Differentially expressed genes with a significant combined effect size across the CD4$^+$ T-cell populations identified by meta-analysis. (A)** Shown are DE genes with a combined effect size identified in the meta-analysis in at least 2 CD4$^+$ T-cell populations. The x-axis represents the combined effect size, the y-axis the "confect" value. Genes that do not show a significant combined effect size (FDR >0.05) have a "confect" value of 0. Genes are color-coded according to the estimated standard error calculated using the `R` package `metafor`. **(B)** To assess the amount of heterogeneity post-hoc, the $I^2$ statistic was calculated. The different plots indicate DE genes with a significant combined effect size across $l$ T-Cell populations. **(C-D)** Number of DE genes with a significant combined effect size (FDR <0.05) across $l$ populations are illustrated by yellow bar graphs

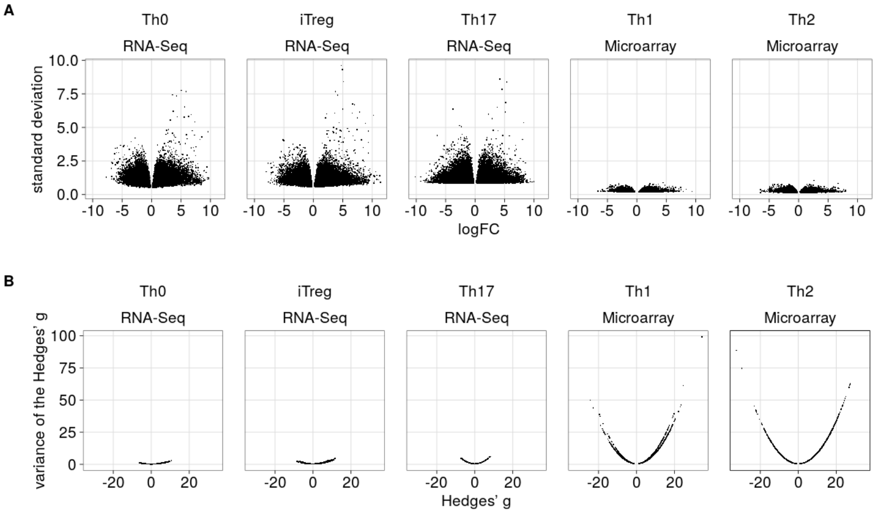

Fig. S9: ***Discovery Set*: Volcano plots of log2 fold changes, Hedges' g values and corresponding estimated variances**. **(A)** Log2 fold changes and estimated standard deviations of all DE genes from the differential gene expression analysis by `limma` for each T-cell population. **(B)** Hedges' g values (equation 3) and estimated variances of the Hedges' g values (equation 4) used as input for meta-analysis.

#### 4.1 Highest ranked genes from the meta-analysis

1h

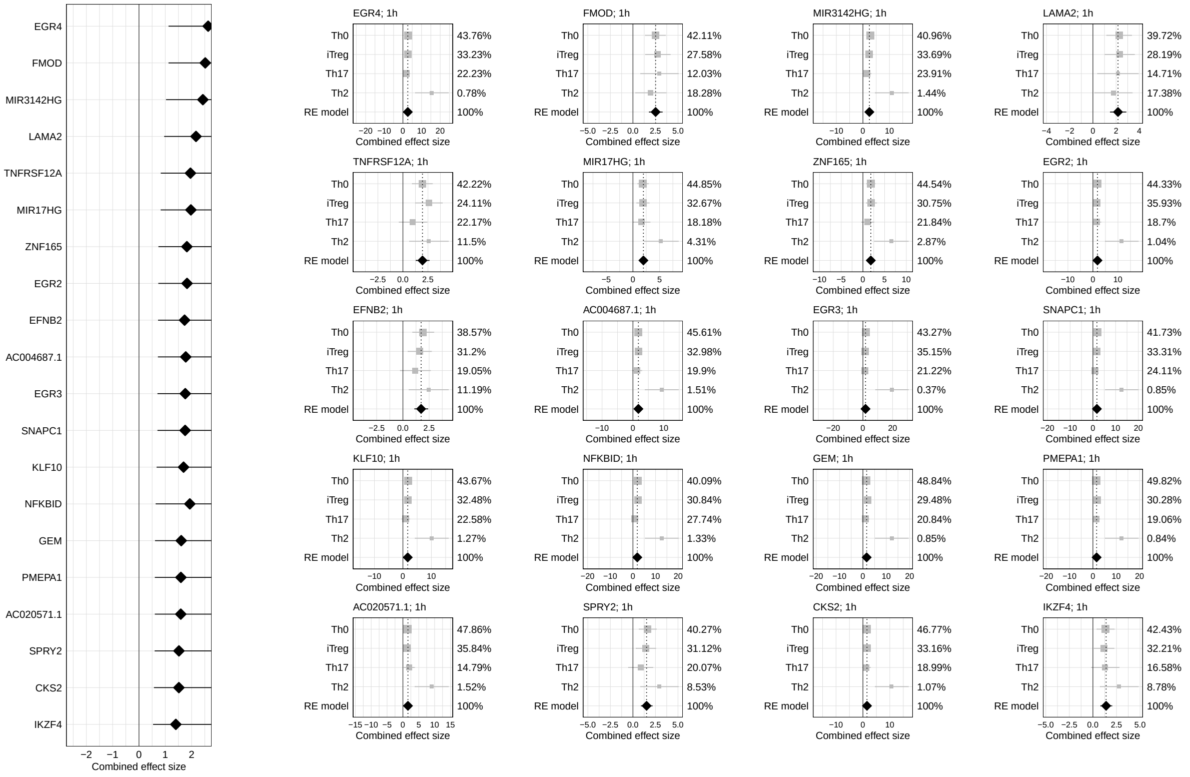

2h

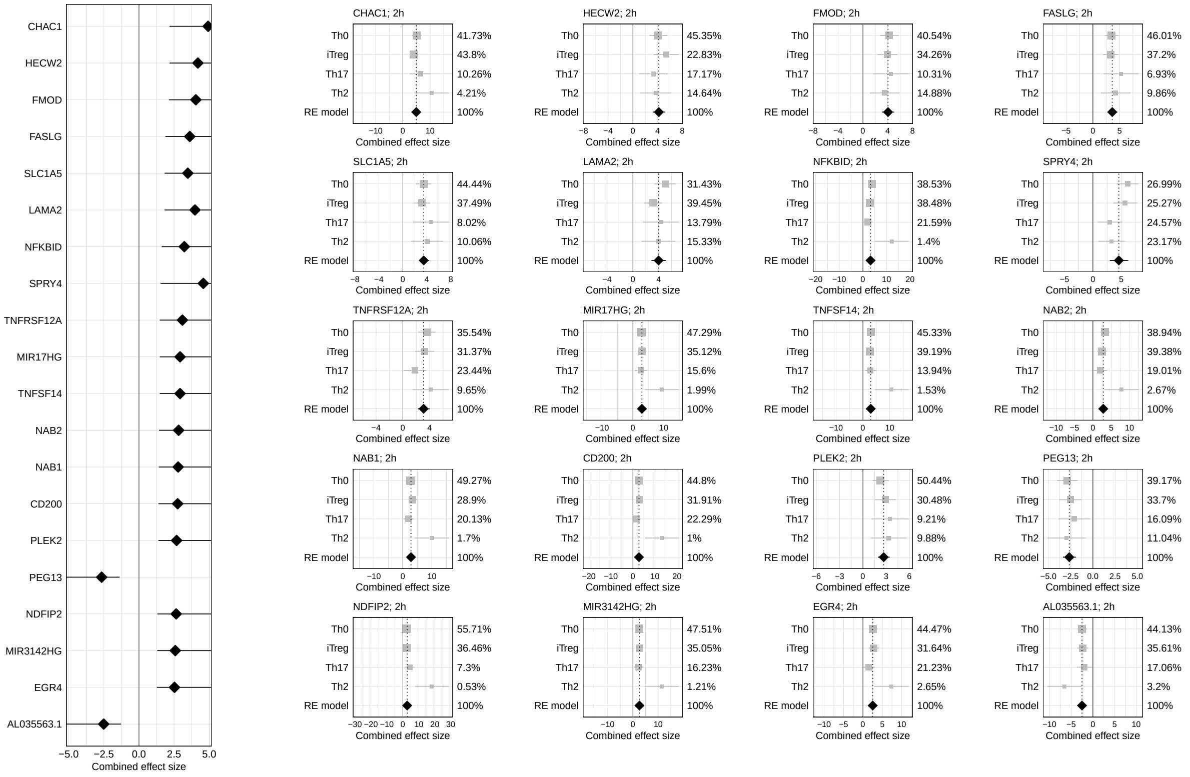

4h

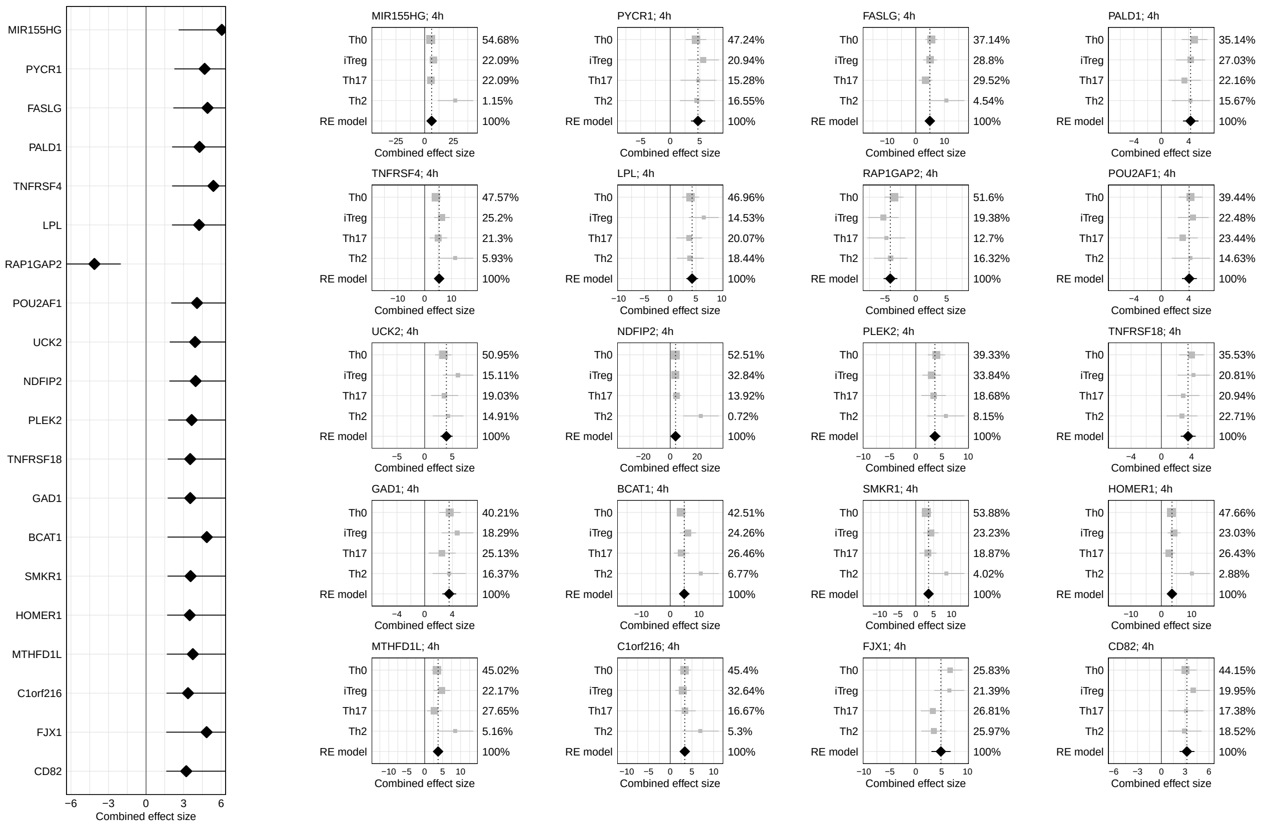
MIR155HG
PYCR1
FASLG
PALD1
TNFRSF4
LPL
RAP1GAP2
POU2AF1
UCK2
NDFIP2
PLEK2
TNFRSF18
GAD1
BCAT1
SMKR1
HOMER1
MTHFD1L
C1orf216
FJX1
CD82
Combined effect size
MIR155HG; 4h
Th0 54.68%
iTreg 22.09%
Th17 22.09%
Th2 1.15%
RE model 100%
PYCR1; 4h
Th0 47.24%
iTreg 20.94%
Th17 15.28%
Th2 16.55%
RE model 100%
FASLG; 4h
Th0 37.14%
iTreg 28.8%
Th17 29.52%
Th2 4.54%
RE model 100%
PALD1; 4h
Th0 35.14%
iTreg 27.03%
Th17 22.16%
Th2 15.67%
RE model 100%
TNFRSF4; 4h
Th0 47.57%
iTreg 25.2%
Th17 21.3%
Th2 5.93%
RE model 100%
LPL; 4h
Th0 46.96%
iTreg 14.53%
Th17 20.07%
Th2 18.44%
RE model 100%
RAP1GAP2; 4h
Th0 51.6%
iTreg 19.38%
Th17 12.7%
Th2 16.32%
RE model 100%
POU2AF1; 4h
Th0 39.44%
iTreg 22.48%
Th17 23.44%
Th2 14.63%
RE model 100%
UCK2; 4h
Th0 50.95%
iTreg 15.11%
Th17 19.03%
Th2 14.91%
RE model 100%
NDFIP2; 4h
Th0 52.51%
iTreg 32.84%
Th17 13.92%
Th2 0.72%
RE model 100%
PLEK2; 4h
Th0 39.33%
iTreg 33.84%
Th17 18.68%
Th2 8.15%
RE model 100%
TNFRSF18; 4h
Th0 35.53%
iTreg 20.81%
Th17 20.94%
Th2 22.71%
RE model 100%
GAD1; 4h
Th0 40.21%
iTreg 18.29%
Th17 25.13%
Th2 16.37%
RE model 100%
BCAT1; 4h
Th0 42.51%
iTreg 24.26%
Th17 26.46%
Th2 6.77%
RE model 100%
SMKR1; 4h
Th0 53.88%
iTreg 23.23%
Th17 18.87%
Th2 4.02%
RE model 100%
HOMER1; 4h
Th0 47.66%
iTreg 23.03%
Th17 26.43%
Th2 2.88%
RE model 100%
MTHFD1L; 4h
Th0 45.02%
iTreg 22.17%
Th17 27.65%
Th2 5.16%
RE model 100%
C1orf216; 4h
Th0 45.4%
iTreg 32.64%
Th17 16.67%
Th2 5.3%
RE model 100%
FJX1; 4h
Th0 25.83%
iTreg 21.39%
Th17 26.81%
Th2 25.97%
RE model 100%
CD82; 4h
Th0 44.15%
iTreg 19.95%
Th17 17.38%
Th2 18.52%
RE model 100%

6h

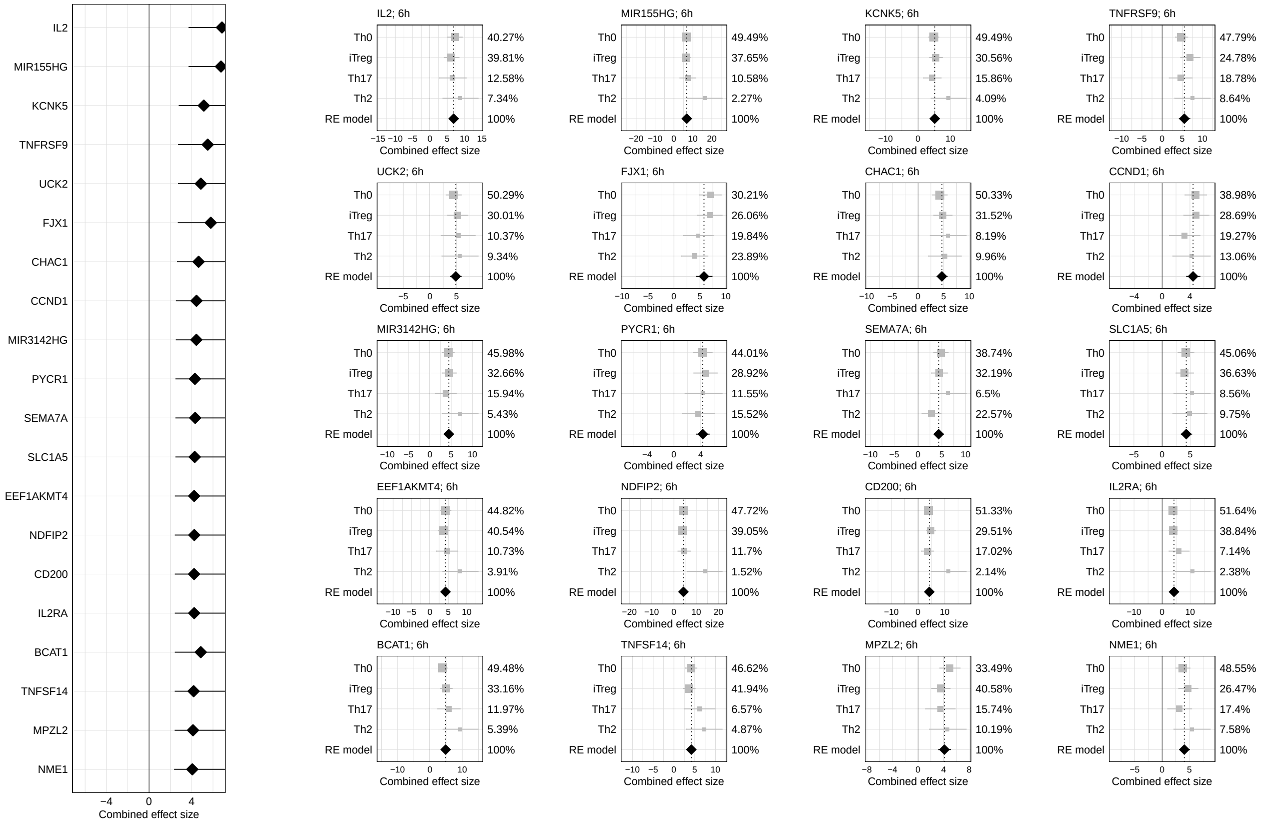
IL2
MIR155HG
KCNK5
TNFRSF9
UCK2
FJX1
CHAC1
CCND1
MIR3142HG
PYCR1
SEMA7A
SLC1A5
EEF1AKMT4
NDFIP2
CD200
IL2RA
BCAT1
TNFSF14
MPZL2
NME1
Combined effect size
IL2; 6h
Th0 40.27%
iTreg 39.81%
Th17 12.58%
Th2 7.34%
RE model 100%
MIR155HG; 6h
Th0 49.49%
iTreg 37.65%
Th17 10.58%
Th2 2.27%
RE model 100%
KCNK5; 6h
Th0 49.49%
iTreg 30.56%
Th17 15.86%
Th2 4.09%
RE model 100%
TNFRSF9; 6h
Th0 47.79%
iTreg 24.78%
Th17 18.78%
Th2 8.64%
RE model 100%
UCK2; 6h
Th0 50.29%
iTreg 30.01%
Th17 10.37%
Th2 9.34%
RE model 100%
FJX1; 6h
Th0 30.21%
iTreg 26.06%
Th17 19.84%
Th2 23.89%
RE model 100%
CHAC1; 6h
Th0 50.33%
iTreg 31.52%
Th17 8.19%
Th2 9.96%
RE model 100%
CCND1; 6h
Th0 38.98%
iTreg 28.69%
Th17 19.27%
Th2 13.06%
RE model 100%
MIR3142HG; 6h
Th0 45.98%
iTreg 32.66%
Th17 15.94%
Th2 5.43%
RE model 100%
PYCR1; 6h
Th0 44.01%
iTreg 28.92%
Th17 11.55%
Th2 15.52%
RE model 100%
SEMA7A; 6h
Th0 38.74%
iTreg 32.19%
Th17 6.5%
Th2 22.57%
RE model 100%
SLC1A5; 6h
Th0 45.06%
iTreg 36.63%
Th17 8.56%
Th2 9.75%
RE model 100%
EEF1AKMT4; 6h
Th0 44.82%
iTreg 40.54%
Th17 10.73%
Th2 3.91%
RE model 100%
NDFIP2; 6h
Th0 47.72%
iTreg 39.05%
Th17 11.7%
Th2 1.52%
RE model 100%
CD200; 6h
Th0 51.33%
iTreg 29.51%
Th17 17.02%
Th2 2.14%
RE model 100%
IL2RA; 6h
Th0 51.64%
iTreg 38.84%
Th17 7.14%
Th2 2.38%
RE model 100%
BCAT1; 6h
Th0 49.48%
iTreg 33.16%
Th17 11.97%
Th2 5.39%
RE model 100%
TNFSF14; 6h
Th0 46.62%
iTreg 41.94%
Th17 6.57%
Th2 4.87%
RE model 100%
MPZL2; 6h
Th0 33.49%
iTreg 40.58%
Th17 15.74%
Th2 10.19%
RE model 100%
NME1; 6h
Th0 48.55%
iTreg 26.47%
Th17 17.4%
Th2 7.58%
RE model 100%

# 12h

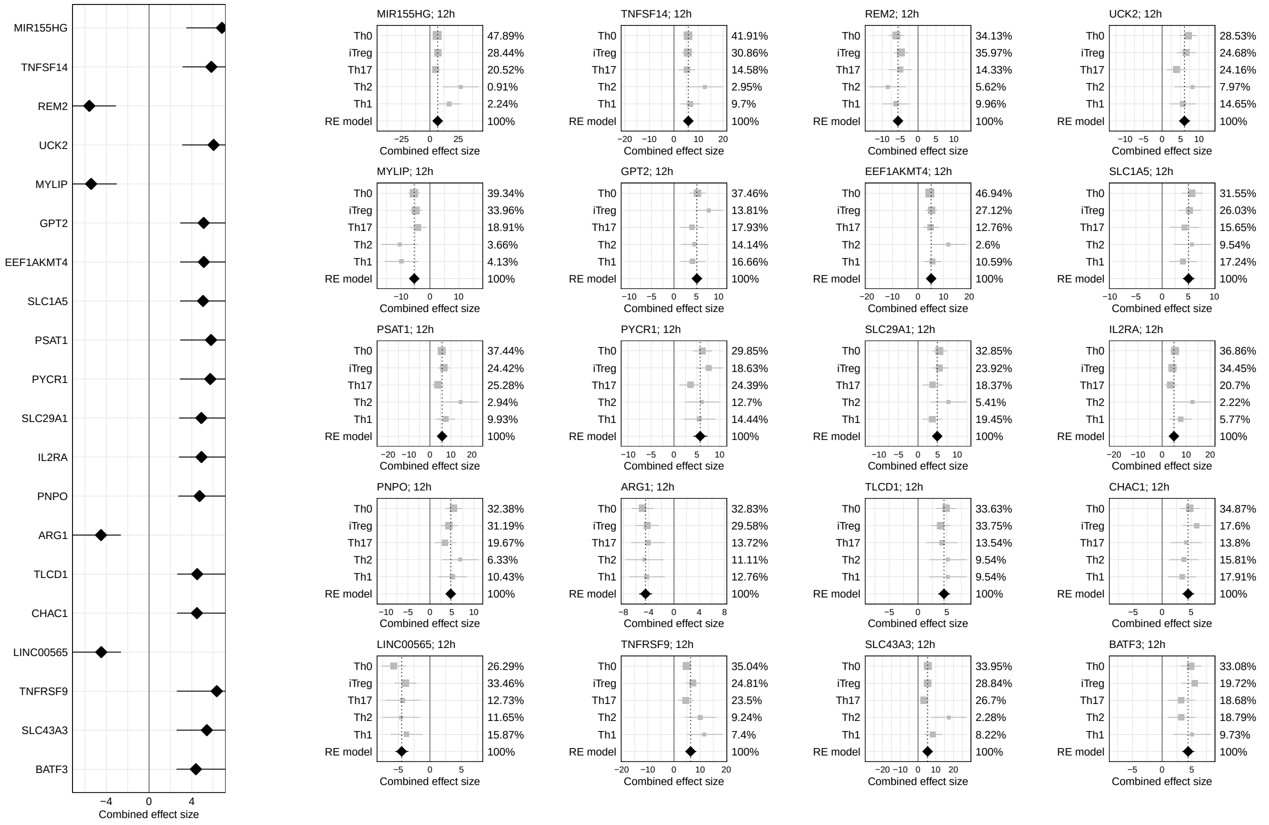
MIR155HG
TNFSF14
REM2
UCK2
MYLIP
GPT2
EEF1AKMT4
SLC1A5
PSAT1
PYCR1
SLC29A1
IL2RA
PNPO
ARG1
TLCD1
CHAC1
LINC00565
TNFRSF9
SLC43A3
BATF3
−4 0 4
Combined effect size
MIR155HG; 12h
Th0 47.89% iTreg 28.44% Th17 20.52% Th2 0.91% Th1 2.24% RE model 100%
TNFSF14; 12h
Th0 41.91% iTreg 30.86% Th17 14.58% Th2 2.95% Th1 9.7% RE model 100%
REM2; 12h
Th0 34.13% iTreg 35.97% Th17 14.33% Th2 5.62% Th1 9.96% RE model 100%
UCK2; 12h
Th0 28.53% iTreg 24.68% Th17 24.16% Th2 7.97% Th1 14.65% RE model 100%
MYLIP; 12h
Th0 39.34% iTreg 33.96% Th17 18.91% Th2 3.66% Th1 4.13% RE model 100%
GPT2; 12h
Th0 37.46% iTreg 13.81% Th17 17.93% Th2 14.14% Th1 16.66% RE model 100%
EEF1AKMT4; 12h
Th0 46.94% iTreg 27.12% Th17 12.76% Th2 2.6% Th1 10.59% RE model 100%
SLC1A5; 12h
Th0 31.55% iTreg 26.03% Th17 15.65% Th2 9.54% Th1 17.24% RE model 100%
PSAT1; 12h
Th0 37.44% iTreg 24.42% Th17 25.28% Th2 2.94% Th1 9.93% RE model 100%
PYCR1; 12h
Th0 29.85% iTreg 18.63% Th17 24.39% Th2 12.7% Th1 14.44% RE model 100%
SLC29A1; 12h
Th0 32.85% iTreg 23.92% Th17 18.37% Th2 5.41% Th1 19.45% RE model 100%
IL2RA; 12h
Th0 36.86% iTreg 34.45% Th17 20.7% Th2 2.22% Th1 5.77% RE model 100%
PNPO; 12h
Th0 32.38% iTreg 31.19% Th17 19.67% Th2 6.33% Th1 10.43% RE model 100%
ARG1; 12h
Th0 32.83% iTreg 29.58% Th17 13.72% Th2 11.11% Th1 12.76% RE model 100%
TLCD1; 12h
Th0 33.63% iTreg 33.75% Th17 13.54% Th2 9.54% Th1 9.54% RE model 100%
CHAC1; 12h
Th0 34.87% iTreg 17.6% Th17 13.8% Th2 15.81% Th1 17.91% RE model 100%
LINC00565; 12h
Th0 26.29% iTreg 33.46% Th17 12.73% Th2 11.65% Th1 15.87% RE model 100%
TNFRSF9; 12h
Th0 35.04% iTreg 24.81% Th17 23.5% Th2 9.24% Th1 7.4% RE model 100%
SLC43A3; 12h
Th0 33.95% iTreg 28.84% Th17 26.7% Th2 2.28% Th1 8.22% RE model 100%
BATF3; 12h
Th0 33.08% iTreg 19.72% Th17 18.68% Th2 18.79% Th1 9.73% RE model 100%
Combined effect size

# 24h

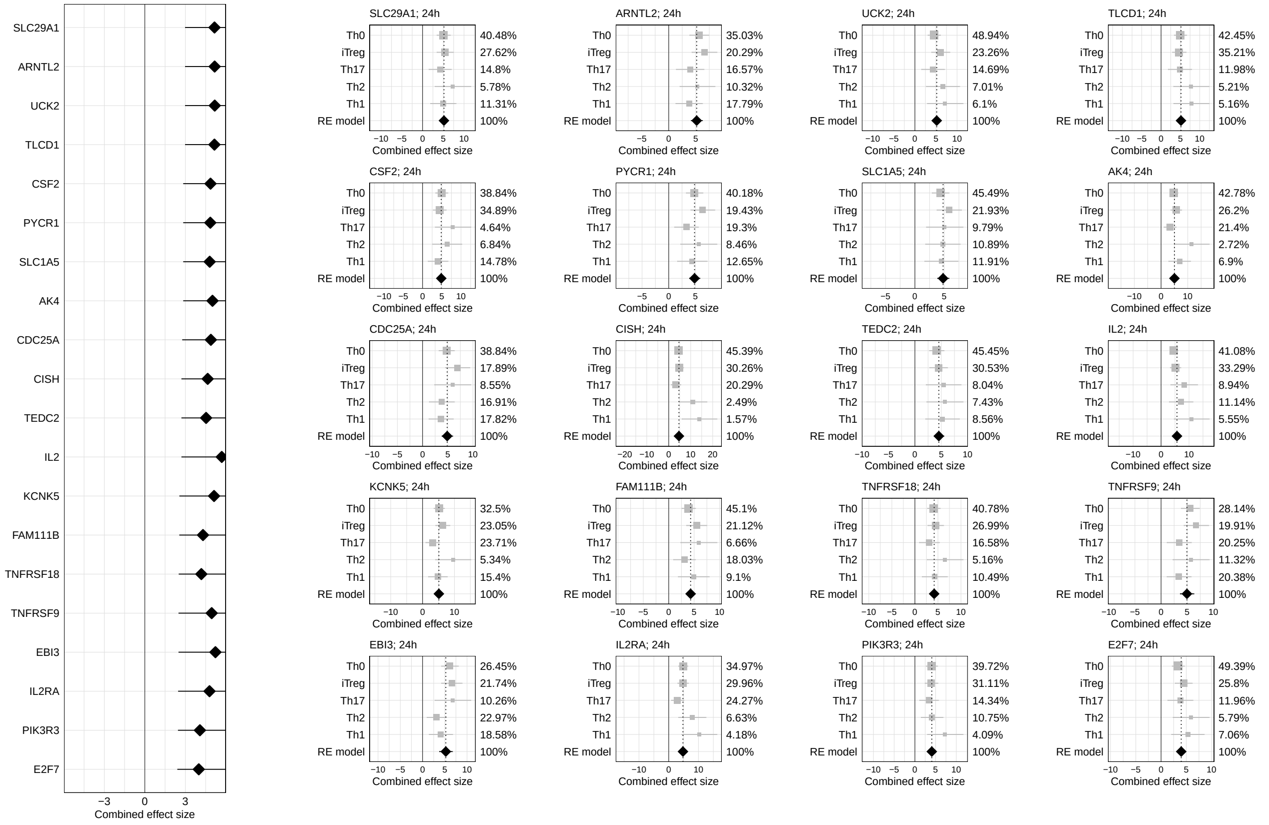
SLC29A1
ARNTL2
UCK2
TLCD1
CSF2
PYCR1
SLC1A5
AK4
CDC25A
CISH
TEDC2
IL2
KCNK5
FAM111B
TNFRSF18
TNFRSF9
EBI3
IL2RA
PIK3R3
E2F7
−3 0 3
Combined effect size
SLC29A1; 24h
Th0 40.48% iTreg 27.62% Th17 14.8% Th2 5.78% Th1 11.31% RE model 100%
ARNTL2; 24h
Th0 35.03% iTreg 20.29% Th17 16.57% Th2 10.32% Th1 17.79% RE model 100%
UCK2; 24h
Th0 48.94% iTreg 23.26% Th17 14.69% Th2 7.01% Th1 6.1% RE model 100%
TLCD1; 24h
Th0 42.45% iTreg 35.21% Th17 11.98% Th2 5.21% Th1 5.16% RE model 100%
CSF2; 24h
Th0 38.84% iTreg 34.89% Th17 4.64% Th2 6.84% Th1 14.78% RE model 100%
PYCR1; 24h
Th0 40.18% iTreg 19.43% Th17 19.3% Th2 8.46% Th1 12.65% RE model 100%
SLC1A5; 24h
Th0 45.49% iTreg 21.93% Th17 9.79% Th2 10.89% Th1 11.91% RE model 100%
AK4; 24h
Th0 42.78% iTreg 26.2% Th17 21.4% Th2 2.72% Th1 6.9% RE model 100%
CDC25A; 24h
Th0 38.84% iTreg 17.89% Th17 8.55% Th2 16.91% Th1 17.82% RE model 100%
CISH; 24h
Th0 45.39% iTreg 30.26% Th17 20.29% Th2 2.49% Th1 1.57% RE model 100%
TEDC2; 24h
Th0 45.45% iTreg 30.53% Th17 8.04% Th2 7.43% Th1 8.56% RE model 100%
IL2; 24h
Th0 41.08% iTreg 33.29% Th17 8.94% Th2 11.14% Th1 5.55% RE model 100%
KCNK5; 24h
Th0 32.5% iTreg 23.05% Th17 23.71% Th2 5.34% Th1 15.4% RE model 100%
FAM111B; 24h
Th0 45.1% iTreg 21.12% Th17 6.66% Th2 18.03% Th1 9.1% RE model 100%
TNFRSF18; 24h
Th0 40.78% iTreg 26.99% Th17 16.58% Th2 5.16% Th1 10.49% RE model 100%
TNFRSF9; 24h
Th0 28.14% iTreg 19.91% Th17 20.25% Th2 11.32% Th1 20.38% RE model 100%
EBI3; 24h
Th0 26.45% iTreg 21.74% Th17 10.26% Th2 22.97% Th1 18.58% RE model 100%
IL2RA; 24h
Th0 34.97% iTreg 29.96% Th17 24.27% Th2 6.63% Th1 4.18% RE model 100%
PIK3R3; 24h
Th0 39.72% iTreg 31.11% Th17 14.34% Th2 10.75% Th1 4.09% RE model 100%
E2F7; 24h
Th0 49.39% iTreg 25.8% Th17 11.96% Th2 5.79% Th1 7.06% RE model 100%
Combined effect size

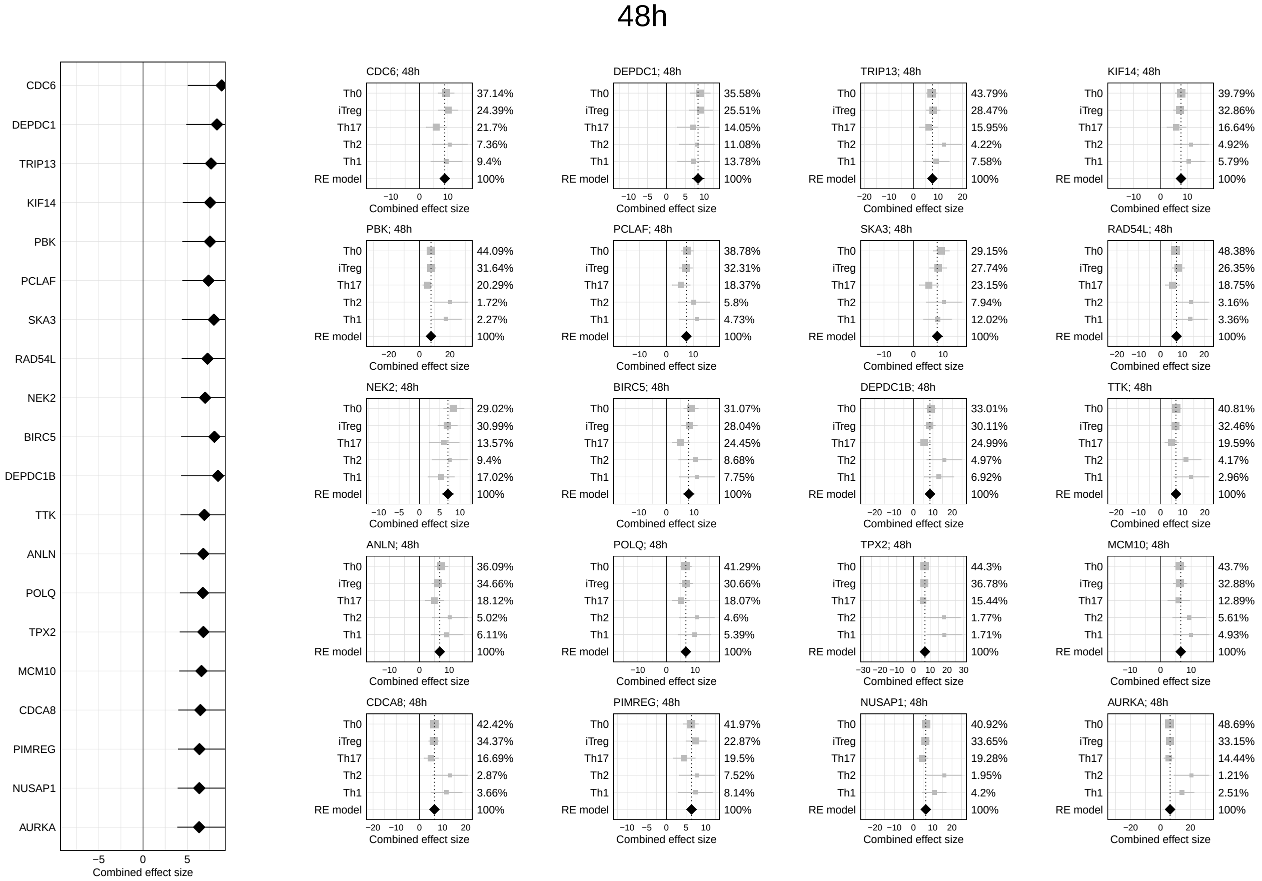
48h
CDC6
DEPDC1
TRIP13
KIF14
PBK
PCLAF
SKA3
RAD54L
NEK2
BIRC5
DEPDC1B
TTK
ANLN
POLQ
TPX2
MCM10
CDCA8
PIMREG
NUSAP1
AURKA
−5 0 5
Combined effect size
CDC6; 48h
Th0 37.14%
iTreg 24.39%
Th17 21.7%
Th2 7.36%
Th1 9.4%
RE model 100%
−10 0 10
Combined effect size
DEPDC1; 48h
Th0 35.58%
iTreg 25.51%
Th17 14.05%
Th2 11.08%
Th1 13.78%
RE model 100%
−10 −5 0 5 10
Combined effect size
TRIP13; 48h
Th0 43.79%
iTreg 28.47%
Th17 15.95%
Th2 4.22%
Th1 7.58%
RE model 100%
−20 −10 0 10 20
Combined effect size
KIF14; 48h
Th0 39.79%
iTreg 32.86%
Th17 16.64%
Th2 4.92%
Th1 5.79%
RE model 100%
−10 0 10
Combined effect size
PBK; 48h
Th0 44.09%
iTreg 31.64%
Th17 20.29%
Th2 1.72%
Th1 2.27%
RE model 100%
−20 0 20
Combined effect size
PCLAF; 48h
Th0 38.78%
iTreg 32.31%
Th17 18.37%
Th2 5.8%
Th1 4.73%
RE model 100%
−10 0 10
Combined effect size
SKA3; 48h
Th0 29.15%
iTreg 27.74%
Th17 23.15%
Th2 7.94%
Th1 12.02%
RE model 100%
−10 0 10
Combined effect size
RAD54L; 48h
Th0 48.38%
iTreg 26.35%
Th17 18.75%
Th2 3.16%
Th1 3.36%
RE model 100%
−20 −10 0 10 20
Combined effect size
NEK2; 48h
Th0 29.02%
iTreg 30.99%
Th17 13.57%
Th2 9.4%
Th1 17.02%
RE model 100%
−10 −5 0 5 10
Combined effect size
BIRC5; 48h
Th0 31.07%
iTreg 28.04%
Th17 24.45%
Th2 8.68%
Th1 7.75%
RE model 100%
−10 0 10
Combined effect size
DEPDC1B; 48h
Th0 33.01%
iTreg 30.11%
Th17 24.99%
Th2 4.97%
Th1 6.92%
RE model 100%
−20 −10 0 10 20
Combined effect size
TTK; 48h
Th0 40.81%
iTreg 32.46%
Th17 19.59%
Th2 4.17%
Th1 2.96%
RE model 100%
−20 −10 0 10 20
Combined effect size
ANLN; 48h
Th0 36.09%
iTreg 34.66%
Th17 18.12%
Th2 5.02%
Th1 6.11%
RE model 100%
−10 0 10
Combined effect size
POLQ; 48h
Th0 41.29%
iTreg 30.66%
Th17 18.07%
Th2 4.6%
Th1 5.39%
RE model 100%
−10 0 10
Combined effect size
TPX2; 48h
Th0 44.3%
iTreg 36.78%
Th17 15.44%
Th2 1.77%
Th1 1.71%
RE model 100%
−30 −20 −10 0 10 20 30
Combined effect size
MCM10; 48h
Th0 43.7%
iTreg 32.88%
Th17 12.89%
Th2 5.61%
Th1 4.93%
RE model 100%
−10 0 10
Combined effect size
CDCA8; 48h
Th0 42.42%
iTreg 34.37%
Th17 16.69%
Th2 2.87%
Th1 3.66%
RE model 100%
−20 −10 0 10 20
Combined effect size
PIMREG; 48h
Th0 41.97%
iTreg 22.87%
Th17 19.5%
Th2 7.52%
Th1 8.14%
RE model 100%
−10 −5 0 5 10
Combined effect size
NUSAP1; 48h
Th0 40.92%
iTreg 33.65%
Th17 19.28%
Th2 1.95%
Th1 4.2%
RE model 100%
−20 −10 0 10 20
Combined effect size
AURKA; 48h
Th0 48.69%
iTreg 33.15%
Th17 14.44%
Th2 1.21%
Th1 2.51%
RE model 100%
−20 0 20
Combined effect size

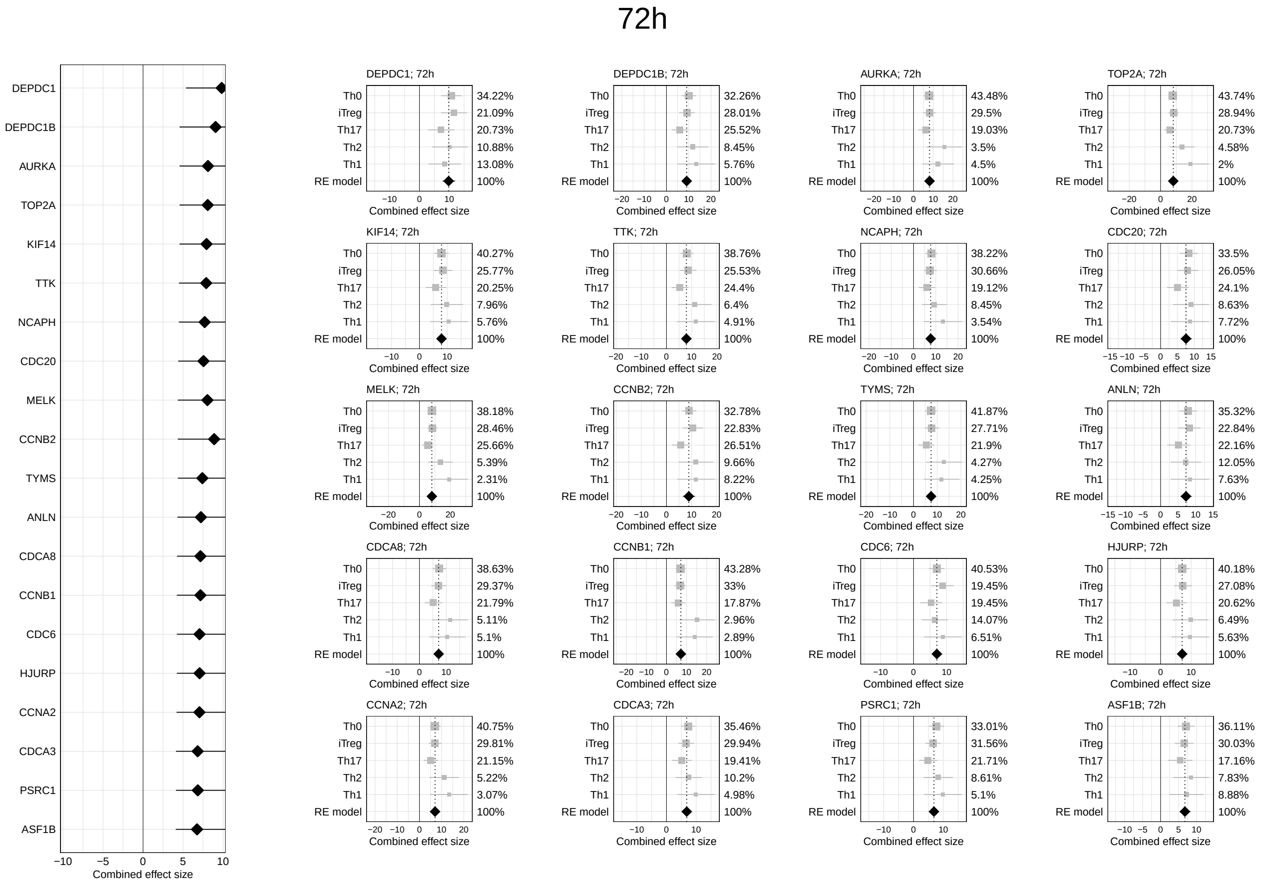
72h
DEPDC1
DEPDC1B
AURKA
TOP2A
KIF14
TTK
NCAPH
CDC20
MELK
CCNB2
TYMS
ANLN
CDCA8
CCNB1
CDC6
HJURP
CCNA2
CDCA3
PSRC1
ASF1B
−10 −5 0 5 10
Combined effect size
DEPDC1; 72h
Th0 34.22%
iTreg 21.09%
Th17 20.73%
Th2 10.88%
Th1 13.08%
RE model 100%
−10 0 10
Combined effect size
DEPDC1B; 72h
Th0 32.26%
iTreg 28.01%
Th17 25.52%
Th2 8.45%
Th1 5.76%
RE model 100%
−20 −10 0 10 20
Combined effect size
AURKA; 72h
Th0 43.48%
iTreg 29.5%
Th17 19.03%
Th2 3.5%
Th1 4.5%
RE model 100%
−20 −10 0 10 20
Combined effect size
TOP2A; 72h
Th0 43.74%
iTreg 28.94%
Th17 20.73%
Th2 4.58%
Th1 2%
RE model 100%
−20 0 20
Combined effect size
KIF14; 72h
Th0 40.27%
iTreg 25.77%
Th17 20.25%
Th2 7.96%
Th1 5.76%
RE model 100%
−10 0 10
Combined effect size
TTK; 72h
Th0 38.76%
iTreg 25.53%
Th17 24.4%
Th2 6.4%
Th1 4.91%
RE model 100%
−20 −10 0 10 20
Combined effect size
NCAPH; 72h
Th0 38.22%
iTreg 30.66%
Th17 19.12%
Th2 8.45%
Th1 3.54%
RE model 100%
−20 −10 0 10 20
Combined effect size
CDC20; 72h
Th0 33.5%
iTreg 26.05%
Th17 24.1%
Th2 8.63%
Th1 7.72%
RE model 100%
−15 −10 −5 0 5 10 15
Combined effect size
MELK; 72h
Th0 38.18%
iTreg 28.46%
Th17 25.66%
Th2 5.39%
Th1 2.31%
RE model 100%
−20 0 20
Combined effect size
CCNB2; 72h
Th0 32.78%
iTreg 22.83%
Th17 26.51%
Th2 9.66%
Th1 8.22%
RE model 100%
−20 −10 0 10 20
Combined effect size
TYMS; 72h
Th0 41.87%
iTreg 27.71%
Th17 21.9%
Th2 4.27%
Th1 4.25%
RE model 100%
−20 −10 0 10 20
Combined effect size
ANLN; 72h
Th0 35.32%
iTreg 22.84%
Th17 22.16%
Th2 12.05%
Th1 7.63%
RE model 100%
−15 −10 −5 0 5 10 15
Combined effect size
CDCA8; 72h
Th0 38.63%
iTreg 29.37%
Th17 21.79%
Th2 5.11%
Th1 5.1%
RE model 100%
−10 0 10
Combined effect size
CCNB1; 72h
Th0 43.28%
iTreg 33%
Th17 17.87%
Th2 2.96%
Th1 2.89%
RE model 100%
−20 −10 0 10 20
Combined effect size
CDC6; 72h
Th0 40.53%
iTreg 19.45%
Th17 19.45%
Th2 14.07%
Th1 6.51%
RE model 100%
−10 0 10
Combined effect size
HJURP; 72h
Th0 40.18%
iTreg 27.08%
Th17 20.62%
Th2 6.49%
Th1 5.63%
RE model 100%
−10 0 10
Combined effect size
CCNA2; 72h
Th0 40.75%
iTreg 29.81%
Th17 21.15%
Th2 5.22%
Th1 3.07%
RE model 100%
−20 −10 0 10 20
Combined effect size
CDCA3; 72h
Th0 35.46%
iTreg 29.94%
Th17 19.41%
Th2 10.2%
Th1 4.98%
RE model 100%
−10 0 10
Combined effect size
PSRC1; 72h
Th0 33.01%
iTreg 31.56%
Th17 21.71%
Th2 8.61%
Th1 5.1%
RE model 100%
−10 0 10
Combined effect size
ASF1B; 72h
Th0 36.11%
iTreg 30.03%
Th17 17.16%
Th2 7.83%
Th1 8.88%
RE model 100%
−10 −5 0 5 10
Combined effect size

Fig. S10: ***Discovery Set*: "Confect" values and forest plots of the 20 highest ranked genes from the meta-analysis using a random-effect model. Left panel:** DE genes with a significant combined effect size (FDR <0.05) across the 4 available populations after 0.5 to 6 hours and across all 5 populations after 12 to 72 hours of activation are shown. Genes are ranked by "confect" values. The diamonds show the estimated combined effect size from the meta-analysis and the inner end of the horizontal line shows the "confect" value (inner confidence bound). **Right panel:** Corresponding forest plots for 20 highest ranked genes. The confidence interval (CI) for each T-cell population is represented by a horizontal line, and the effect size (Hedges'g values) is represented by a square. The size of the square corresponds to the weight of the population in the meta-analysis. These weights are also shown as numbers on the right-hand side. The combined effect is represented by a diamond and the CI by the vertical line.

#### 4.2 Enrichment analysis

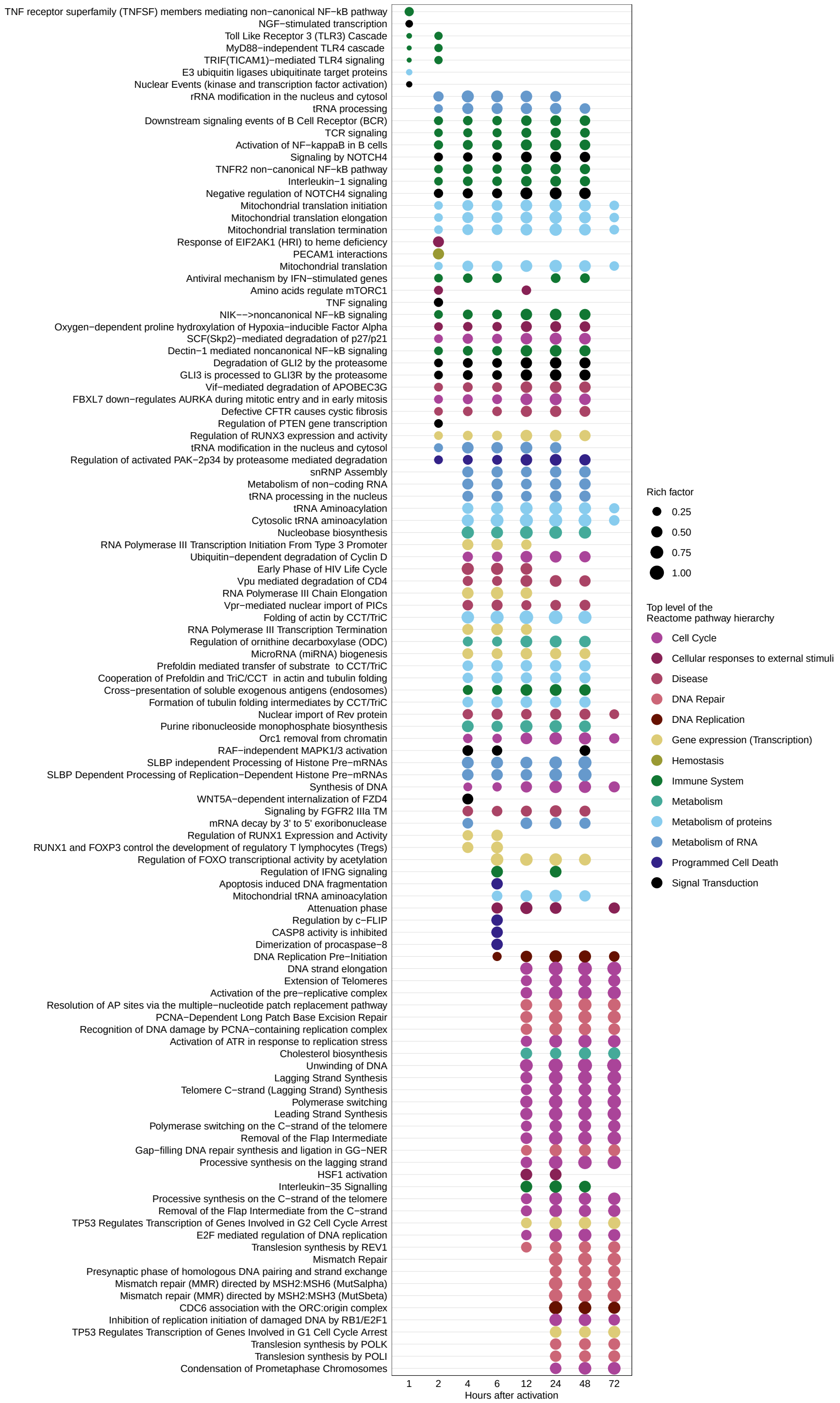

Fig. S11: ***Discovery Set*: Reactome pathway enrichment analysis of DE genes with a significant combined effect size for each analysis time point.** Dot plot depict the top 30 (sorted by rich factor) significantly enriched Reactome pathways (FDR <0.05) for DE genes with a significant combined effect size in 4 (0.5 to 6 hours) and 5 (12 to 72 hours) $CD4^{+}$ T-cell populations. The dot size indicates the rich factor which is the number of DE genes with a significant combined effect size in the pathway divided by the number of background genes in the pathway. Colors depicted the uppermost hierarchical level of Reactome database.

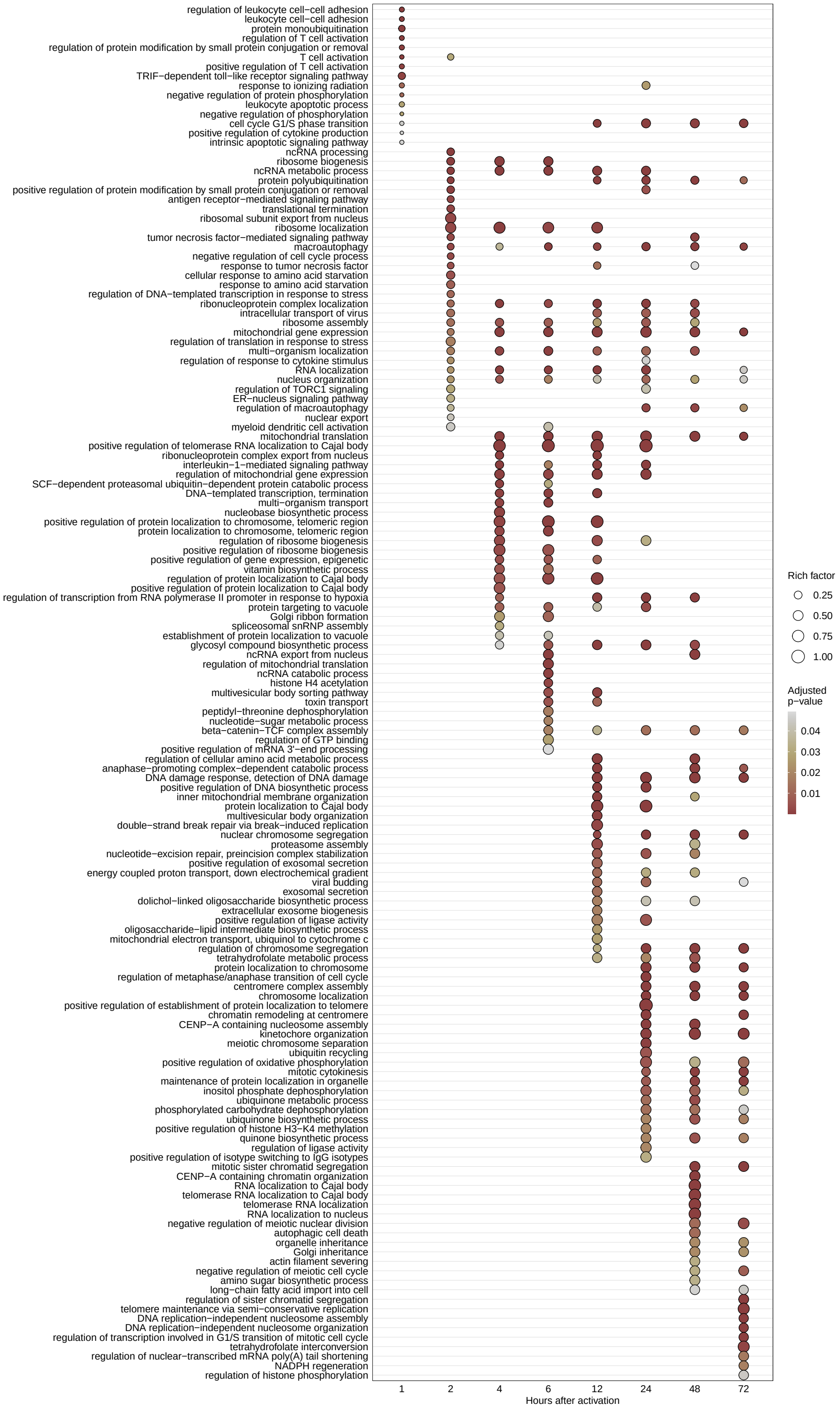

Fig. S12: ***Discovery Set*: GO term enrichment analysis of DE genes with a significant combined effect for each analysis time point.** Dot plot depict the top 30 (sorted by rich factor) significantly enriched GO term for biological processes (FDR <0.05) for DE genes with a significant combined effect size in 4 (0.5 to 6 hours) and 5 (12 to 72 hours) $CD4^{+}$ T-cell populations. The dot size indicates the rich factor which is the number of DE genes with a significant combined effect size in the GO term divided by the number of background genes in the GO term.

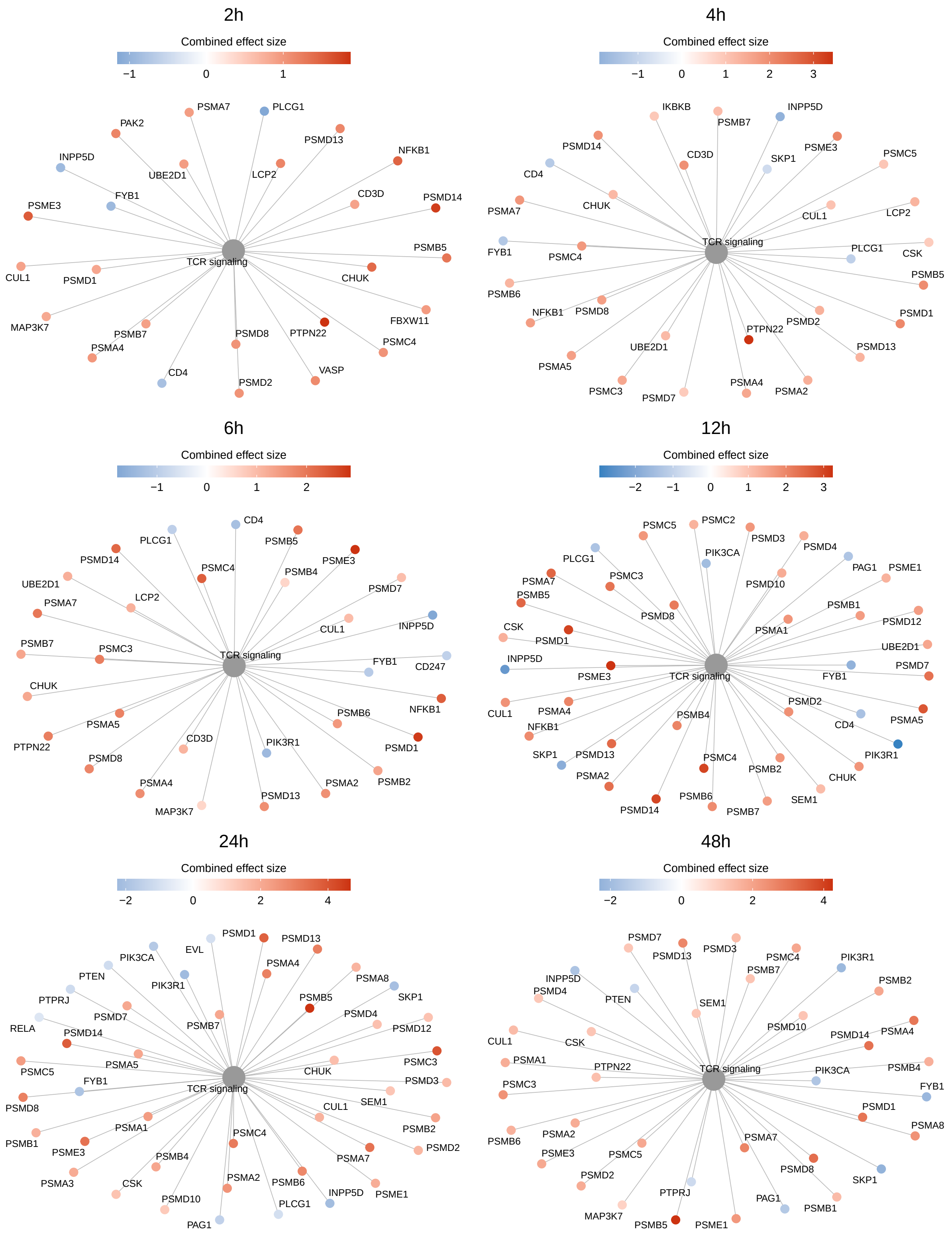

Fig. S13: ***Discovery Set*: T-cell receptor pathway**. Significantly (FDR <0.05) enriched Reactome T-cell receptor (TCR) pathway of DE genes with a significant combined effect size (FDR <0.05) from the meta-analysis. The linkages of genes and the TCR pathway are shown as a network. Genes with a significantly negative combined effect size are shown in blue, genes with a significantly positive combined effect are shown in red. Only those time points are shown in which the TCR signalling pathway is also significantly enriched.

#### 5 Non-negative matrix factorization

Non-negative matrix factorization (NMF) is a form of dimension reduction where the $n \times m$ matrix $X$ is approximated by:

$$\underset{\mathbb{R}^{n\times m}}{X} \approx \underset{\mathbb{R}^{n\times k}}{W} \times \underset{\mathbb{R}^{k\times m}}{H} \text{ such that } W, H \geq 0$$

where the rank $k \leq max(m, n)$ ($k$ = number of metagenes). We want to find a basis $W*$ and coefficients $H*$ matrix which minimize the reconstruction error according to:

$$\underset{W,H\geq 0}{min}\, D(X, WH) \text{ w.r.t. } W, H$$

where $D(X, WH)$ is a loss function that measures the quality of the approximation. You can use any distance measure which evaluates how well the low-dimensional matrix $WH$ approximates X. Here we used the Kullback-Leibler divergence

$$D(X, WH) = \sum_{i,j}(X_{ij}(\log \frac{X_{ij}}{(WH)_{ij}}) - X_{ij} + (WH)_{ij}).$$

The matrices $W$ and $H$ are found by maximizing the log-likelihood under an independent Poisson assumption of $X$. However, this is a non-convex optimization problem. The convexity is given for W or H, but is not given in respect to both variables together and therefore there is no unique solution. The optimization can be done with multiplicative rules [26] which update W and H according to the following rule:

$$H_{kj} \leftarrow H_{kj} \frac{\sum_{i=1}^{n}(W_{ik}X_{ij})/(WH)_{ij}}{\sum_{i=1}^{n} W_{ik}} \quad W_{ik} \leftarrow W_{ik} \frac{\sum_{j=1}^{m}(H_{kj}X_{ij})/(WH)_{ij}}{\sum_{j=1}^{m}}$$

The initialization of $W$ and $H$ was generated as random seed matrices drawn from a uniform distribution within the same range as the entries in the matrix $X$.
We used the model selection method proposed by Brunet et al. [26] to estimate the optimal number of ranks. We performed NMF on the normalized data (see main methods) by repeating the rank value in the interval 2–10. Due to the unavoidably converges to local minima, we ran the NMF algorithm 200 times for each rank. For each run, a connectivity matrix $C$ of the size $m \times m$ was calculated. Matrix $C$ has an entry of 1 if samples $i$ and $j$ cluster together (assigned to the same metagene) and 0 otherwise, where $ij = 1, \ldots, m$. The sample assignment here depends on the relative values in each column of $H$. The consensus matrix $\overline{C}$ is computed as the average connectivity matrix over the 200 runs. The entries of $\overline{C}$ reflect the probability that samples i and j belong to the same cluster. This yields the empirical probability for each pair of samples clustered together. Hierarchical clustering was performed to reorder the samples (rows and columns of the consensus matrix).
The visualization of the clustered matrix $\overline{C}$ can be used as a qualitative measurement to assess the stability of the clusters (see Figure S15A). As a quantitative measurement we can use the cophenetic correlation coefficient, which indicates the dispersion of the consensus matrix $\overline{C}$. It is computed as the Pearson correlation of two distance matrices. The first represent the sample distances induced by the consensus matrix. The second is the cophenetic distance obtained by hierarchical clustering [26] (see Figure S15B). If all entries in the matrix $\overline{C}$ are 1 or 0, the cophenetic correlation coefficient is 1. As another measure for cluster performance of the consensus matrix, we used the average silhouette width (see Figure S15C).

##### 5.0.1 Defining of metagene associated genes

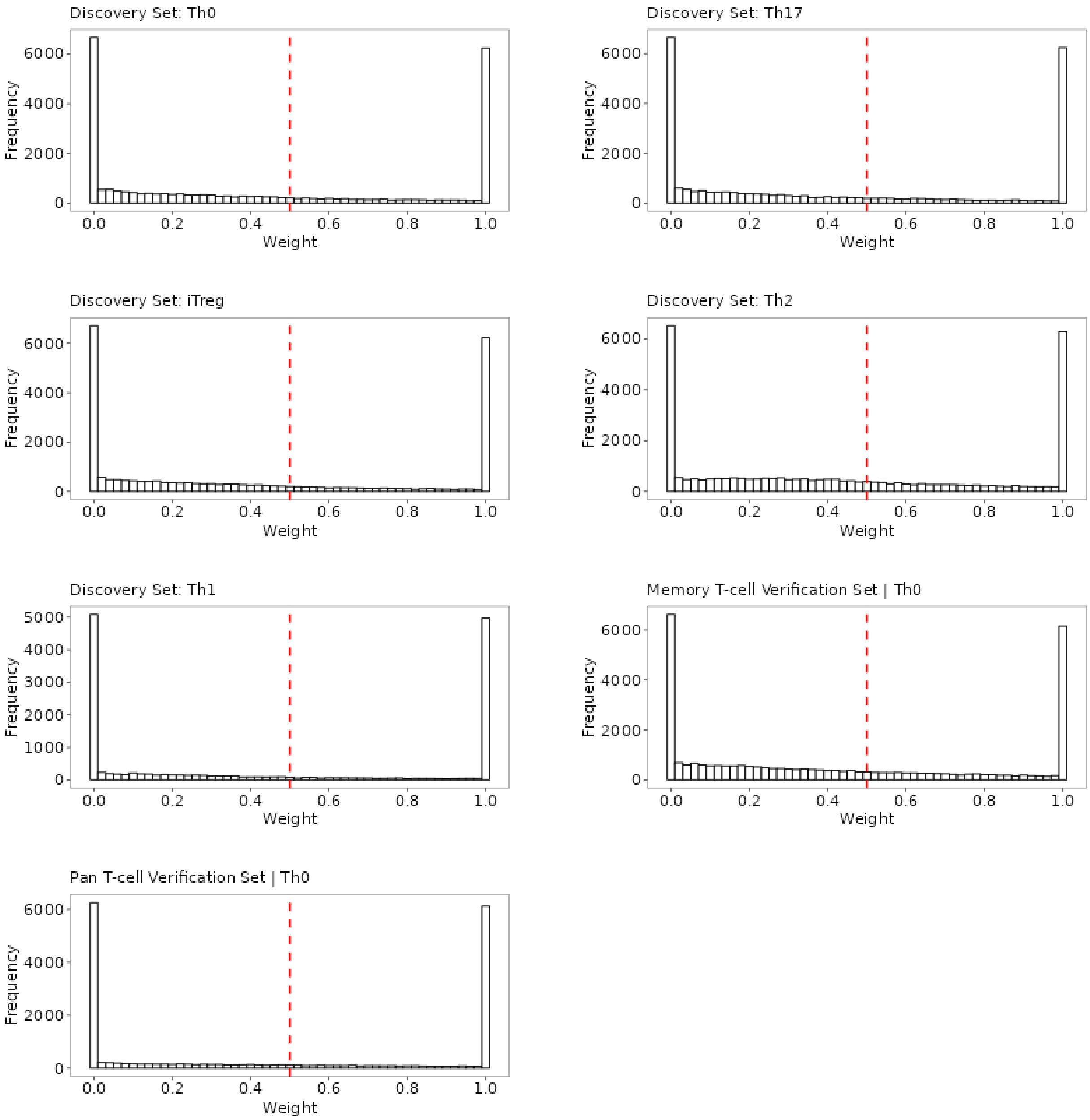

Fig. S14: ***Discovery and Verification Sets*: Distribution of normalized weights from the gene signature matrix $W$ for each T-cell population**. Each row in the gene signature matrix $W$ was min-max normalized. The red dashed line represents the cutoff ( >0.5) at which gene $i$ from the matrix $W$ is assigned to metagene $j$.

##### 5.0.2 NMF consensus clustering

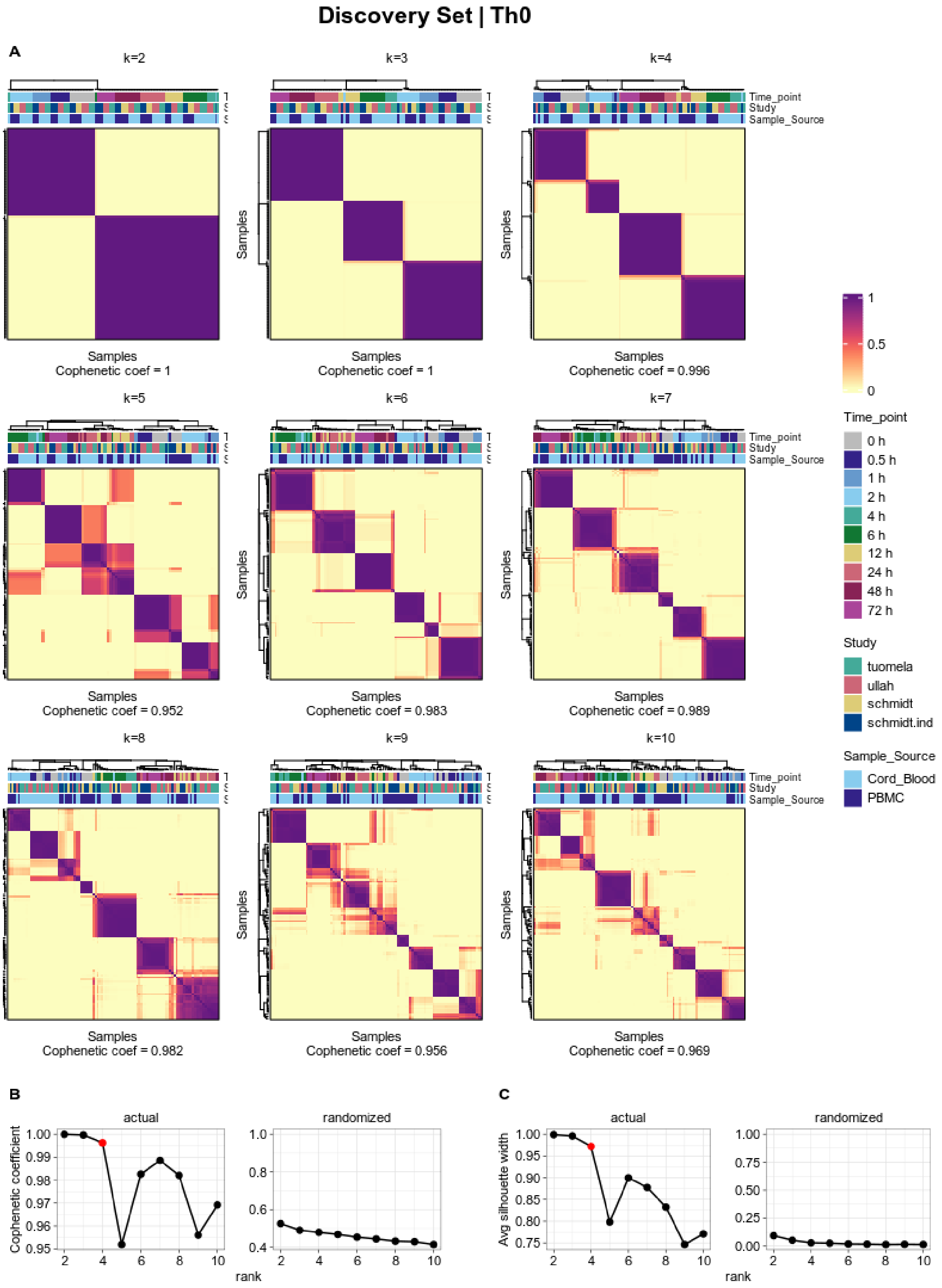

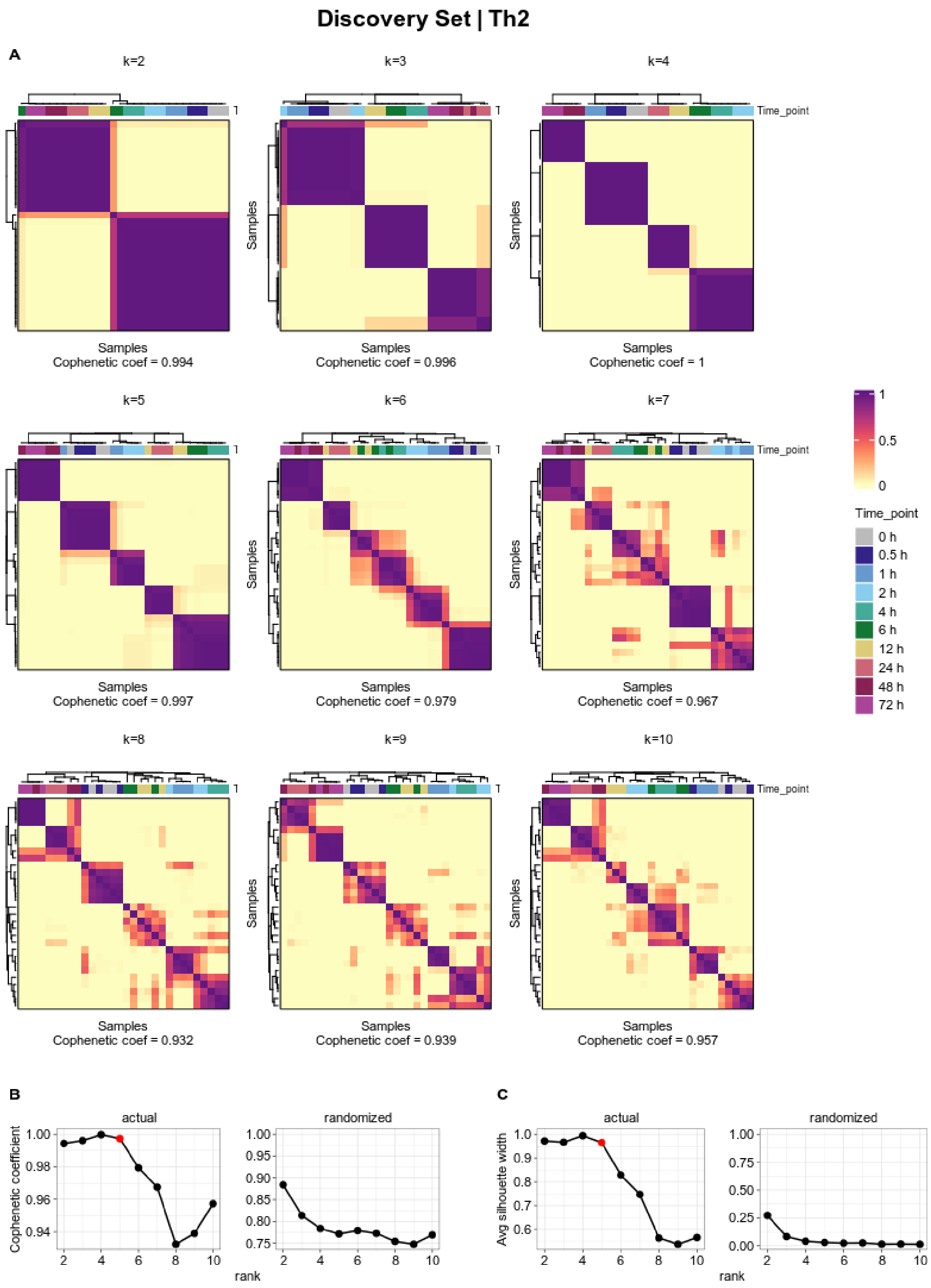
Discovery Set | Th2
A
k=2
k=3
k=4
Time_point
Samples
Samples
Cophenetic coef = 0.994
Cophenetic coef = 0.996
Cophenetic coef = 1
k=5
k=6
k=7
Cophenetic coef = 0.997
Cophenetic coef = 0.979
Cophenetic coef = 0.967
k=8
k=9
k=10
Cophenetic coef = 0.932
Cophenetic coef = 0.939
Cophenetic coef = 0.957
1
0.5
0
Time_point
0 h
0.5 h
1 h
2 h
4 h
6 h
12 h
24 h
48 h
72 h
B
C
actual
randomized
Cophenetic coefficient
Avg silhouette width
rank

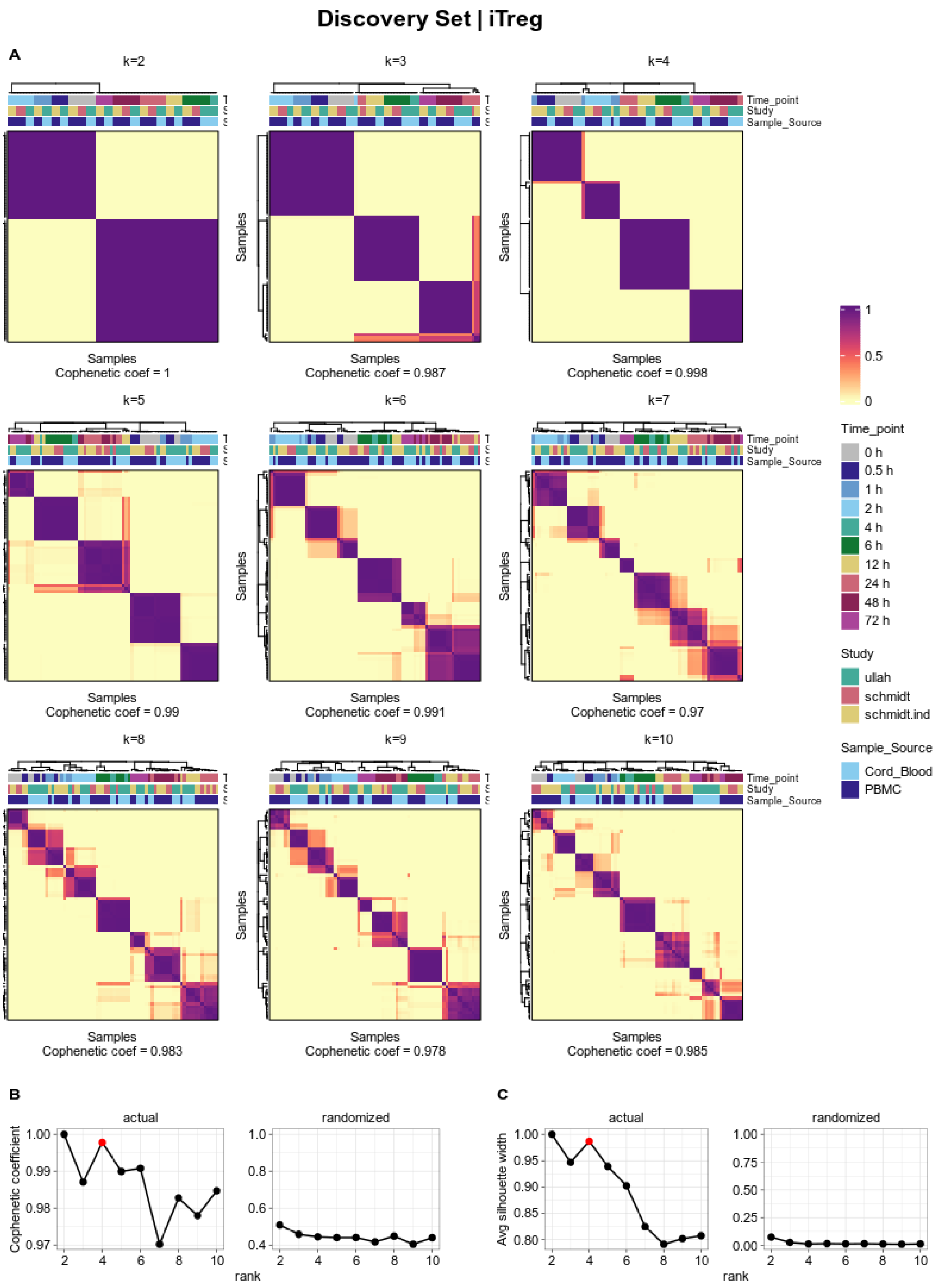
Discovery Set | iTreg
A
k=2
k=3
k=4
k=5
k=6
k=7
k=8
k=9
k=10
Time_point
Study
Sample_Source
Samples
Cophenetic coef = 1
Cophenetic coef = 0.987
Cophenetic coef = 0.998
Cophenetic coef = 0.99
Cophenetic coef = 0.991
Cophenetic coef = 0.97
Cophenetic coef = 0.983
Cophenetic coef = 0.978
Cophenetic coef = 0.985
1
0.5
0
Time_point
0 h
0.5 h
1 h
2 h
4 h
6 h
12 h
24 h
48 h
72 h
Study
ullah
schmidt
schmidt.ind
Sample_Source
Cord_Blood
PBMC
B
C
actual
randomized
Cophenetic coefficient
Avg silhouette width
rank

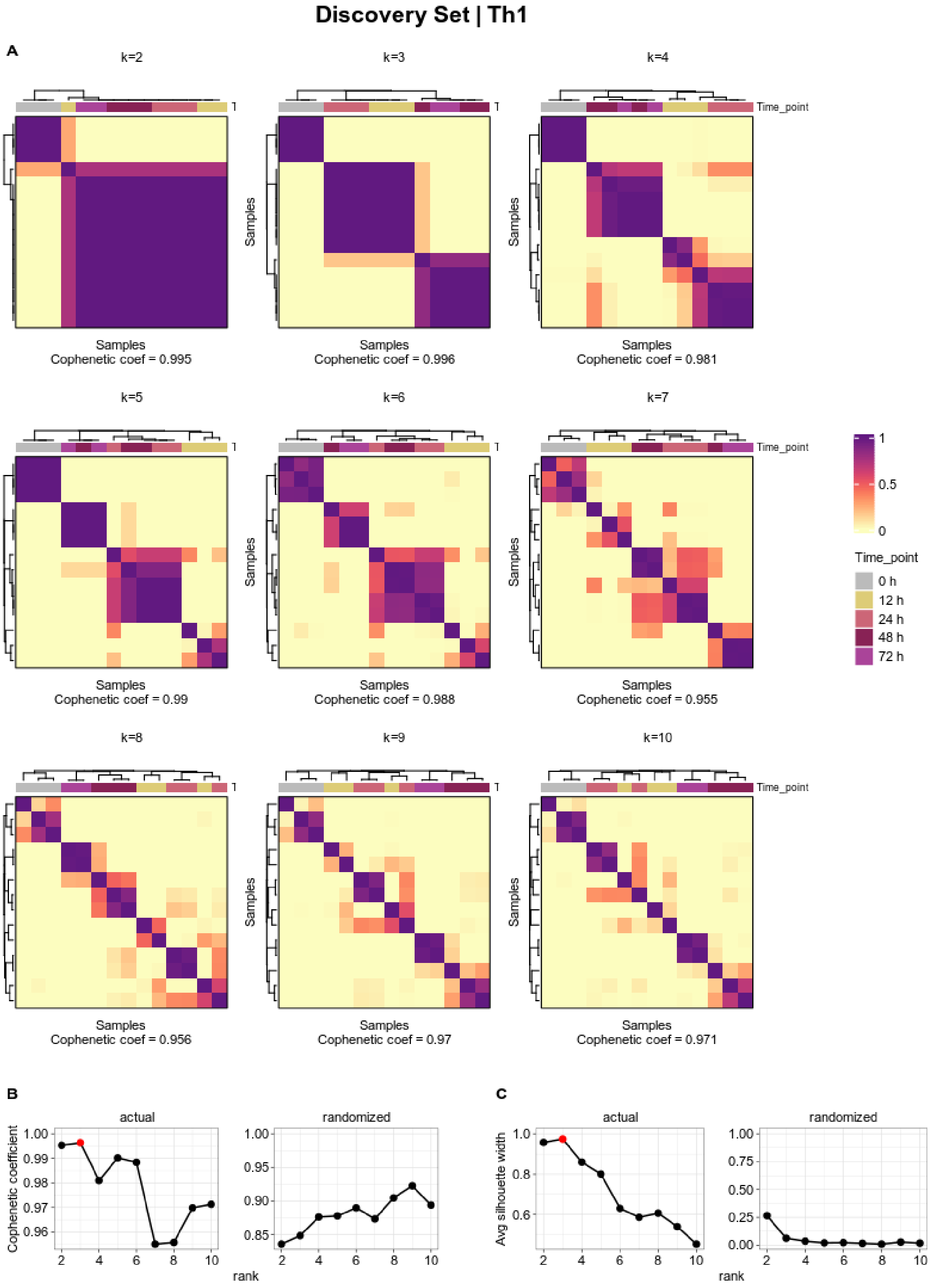
Discovery Set | Th1
A
k=2
k=3
k=4
k=5
k=6
k=7
k=8
k=9
k=10
Time_point
Samples
Cophenetic coef = 0.995
Cophenetic coef = 0.996
Cophenetic coef = 0.981
Cophenetic coef = 0.99
Cophenetic coef = 0.988
Cophenetic coef = 0.955
Cophenetic coef = 0.956
Cophenetic coef = 0.97
Cophenetic coef = 0.971
1
0.5
0
Time_point
0 h
12 h
24 h
48 h
72 h
B
actual
randomized
Cophenetic coefficient
rank
C
Avg silhouette width

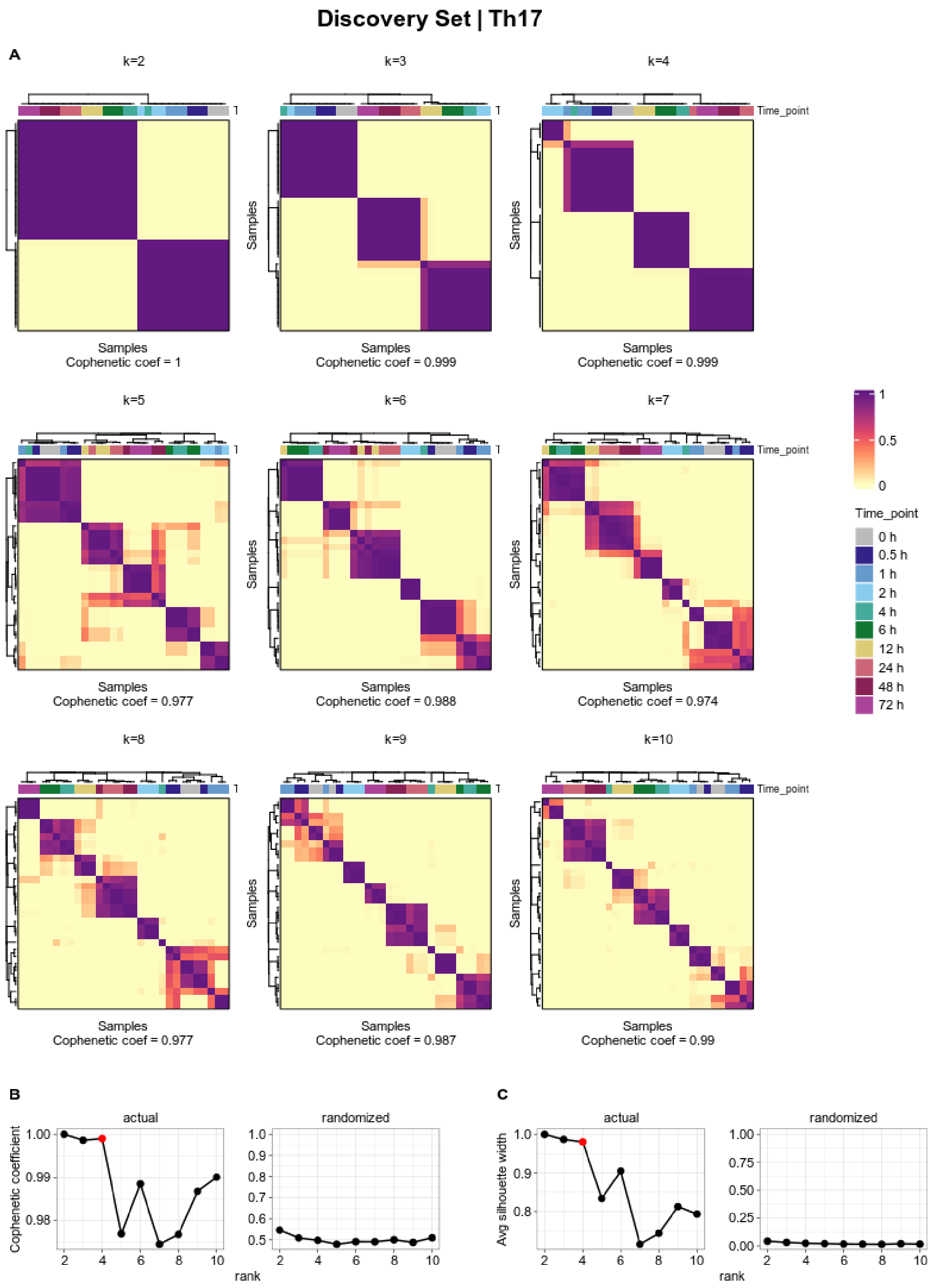
Discovery Set | Th17
A
k=2
k=3
k=4
Time_point
Samples
Cophenetic coef = 1
Cophenetic coef = 0.999
Cophenetic coef = 0.999
k=5
k=6
k=7
Cophenetic coef = 0.977
Cophenetic coef = 0.988
Cophenetic coef = 0.974
k=8
k=9
k=10
Cophenetic coef = 0.977
Cophenetic coef = 0.987
Cophenetic coef = 0.99
1
0.5
0
Time_point
0 h
0.5 h
1 h
2 h
4 h
6 h
12 h
24 h
48 h
72 h
B
C
actual
randomized
Cophenetic coefficient
Avg silhouette width
rank

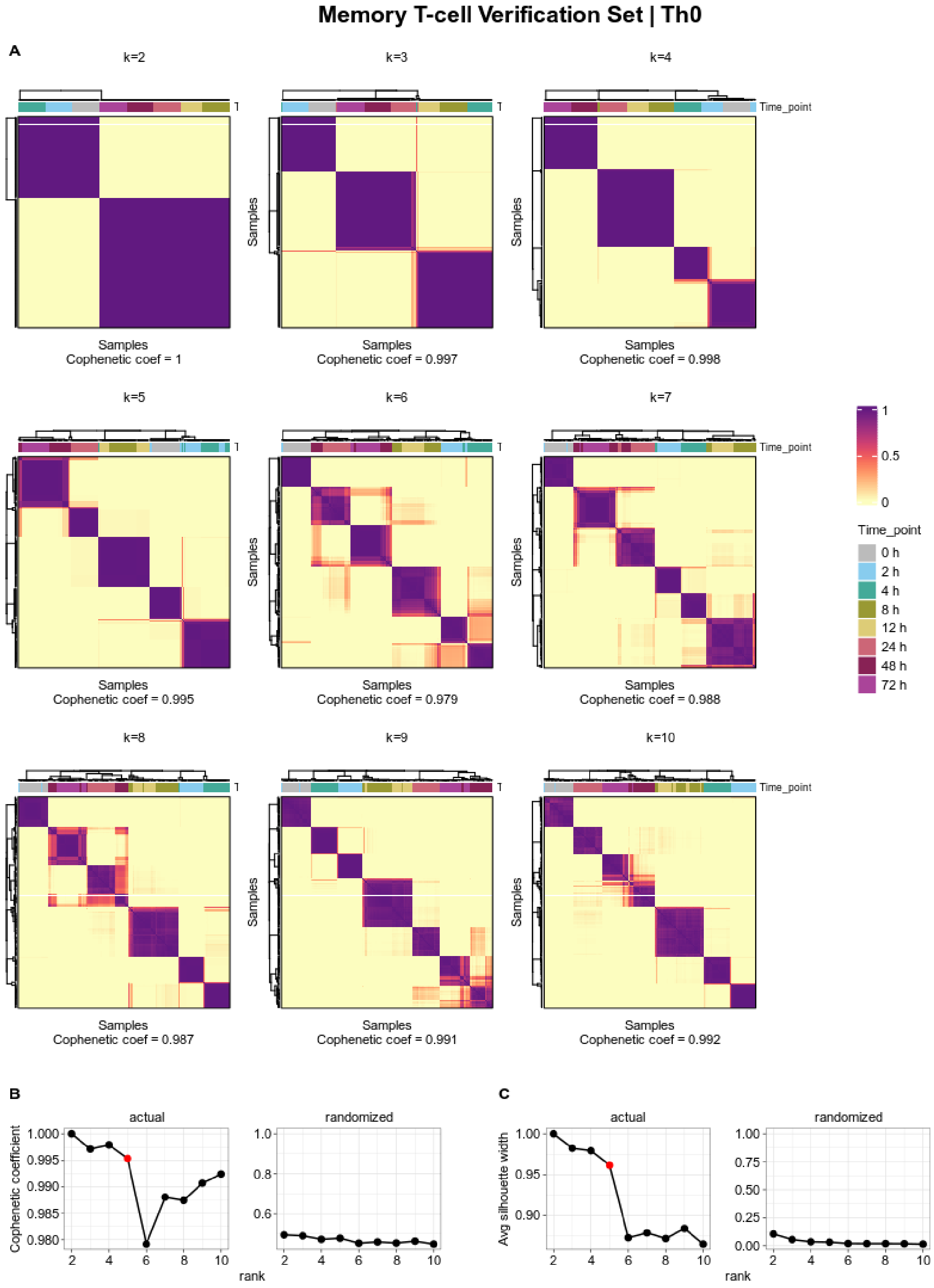
Memory T-cell Verification Set | Th0
A
k=2
k=3
k=4
k=5
k=6
k=7
k=8
k=9
k=10
Time_point
Samples
Cophenetic coef = 1
Cophenetic coef = 0.997
Cophenetic coef = 0.998
Cophenetic coef = 0.995
Cophenetic coef = 0.979
Cophenetic coef = 0.988
Cophenetic coef = 0.987
Cophenetic coef = 0.991
Cophenetic coef = 0.992
1
0.5
0
0 h
2 h
4 h
8 h
12 h
24 h
48 h
72 h
B
C
actual
randomized
Cophenetic coefficient
Avg silhouette width
rank

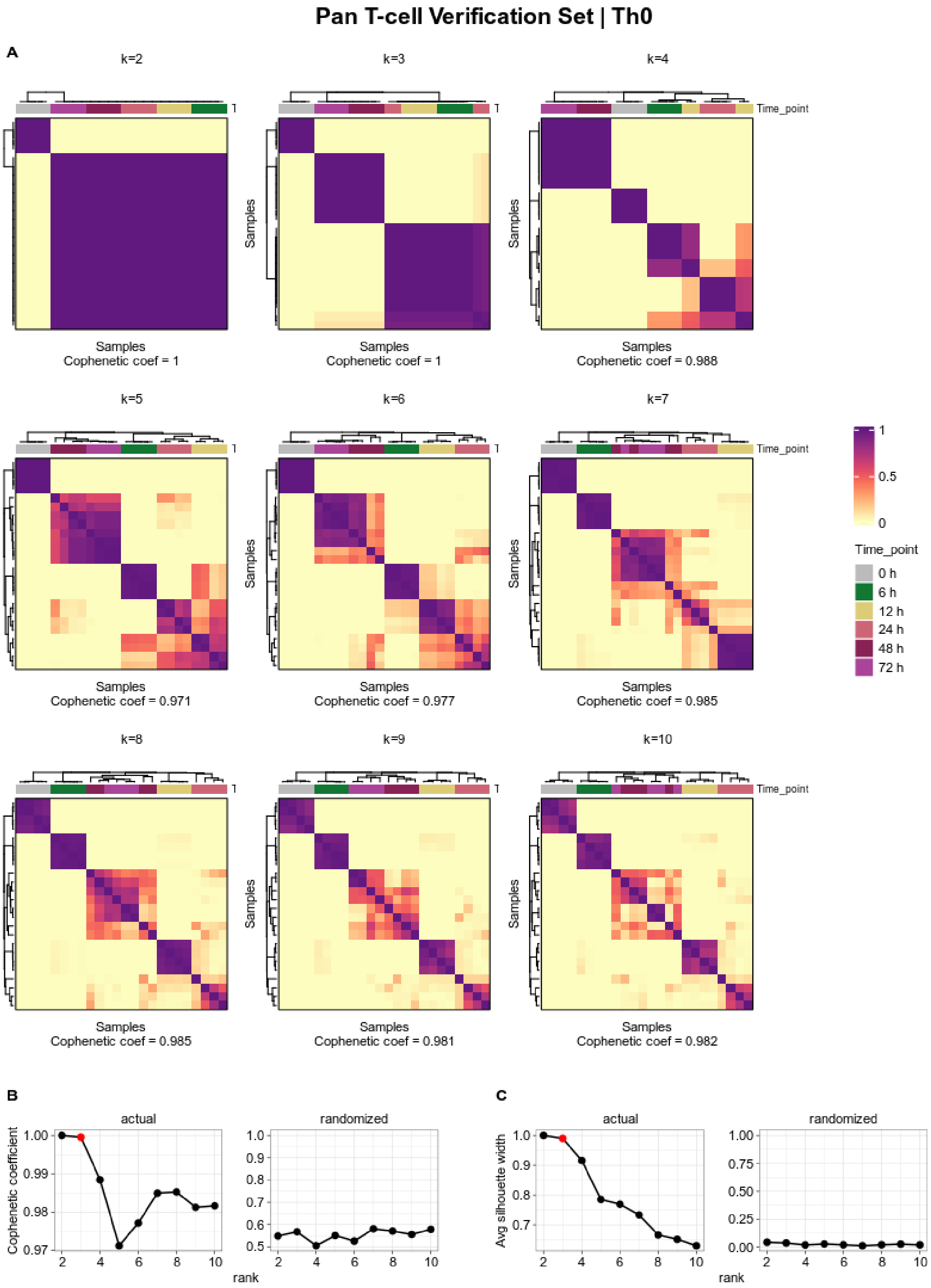

Fig. S15: **NMF consensus-clustering**. **(A)** Consensus matrices averaging 200 connectivity matrices and computed for factorization ranks k = 2–10 (number of metagenes). NMF was performed for each T-cell population separately. NMF computation and model selection were performed according to Brunet et al. [26] as described in the methods. Samples are hierarchically clustered by using distances derived from consensus clustering matrix entries, colored from 0 (samples are never in the same cluster) to 1 (samples are always in the same cluster). **(B-C)** Cophenetic correlation coefficient for the above hierarchically clustered matrices and average silhouette-width plots. NMF was also run with randomized data with same parameters as for the actual data. Red colored dots represent the chosen factorization rank for downstream analysis.

#### 5.1 *Discovery Set*

##### 5.1.1 25 highest ranked genes associated with metagenes

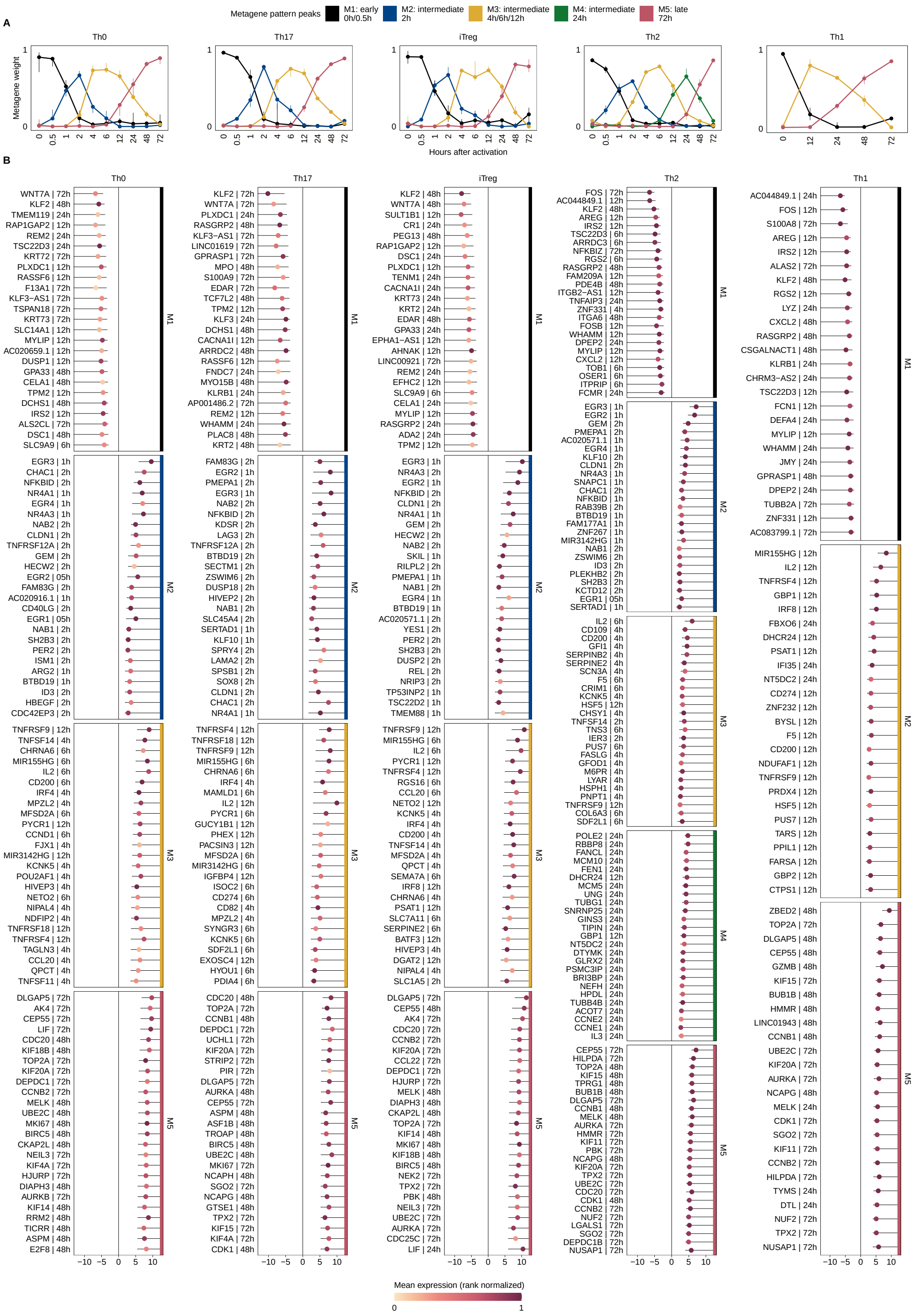

Fig. S16: ***Discovery Set*: 25 highest ranked genes associated with metagenes for each T-cell population**. **(A)** The pattern matrix for each T-cell population from the *Discovery Set* is shown as continuous profiles, with samples assigned to time points of activation. We scaled each column in the pattern matrix to sum up to one. Dots depict the median weights for all samples from identical analysis time points. Vertical lines represent interquartile ranges. We annotated and colored the metagenes based on their maximum median values across all analysis time points. The time point with maximum median value is depicted in the legend. **(B)** For each CD4$^+$ T-cell population and gene used for NMF, we used the highest absolute "confect" value estimated in the DGEA across all contrasts (e.g. 12h vs. 0h). Genes are ranked by "confect" values. Dots represent log2 fold changes for contrasts with the highest absolute "confect" value. The time point to the right of the gene represents the contrast with the highest absolute "confect" value. Color of dots correspond to rank normalized average expression values of the activation group in the contrast with the highest absolute "confect". The inner end of the horizontal line shows the "confect" value (inner confidence bound).

##### 5.1.2 Enrichment analysis for gene sets associated with metagenes

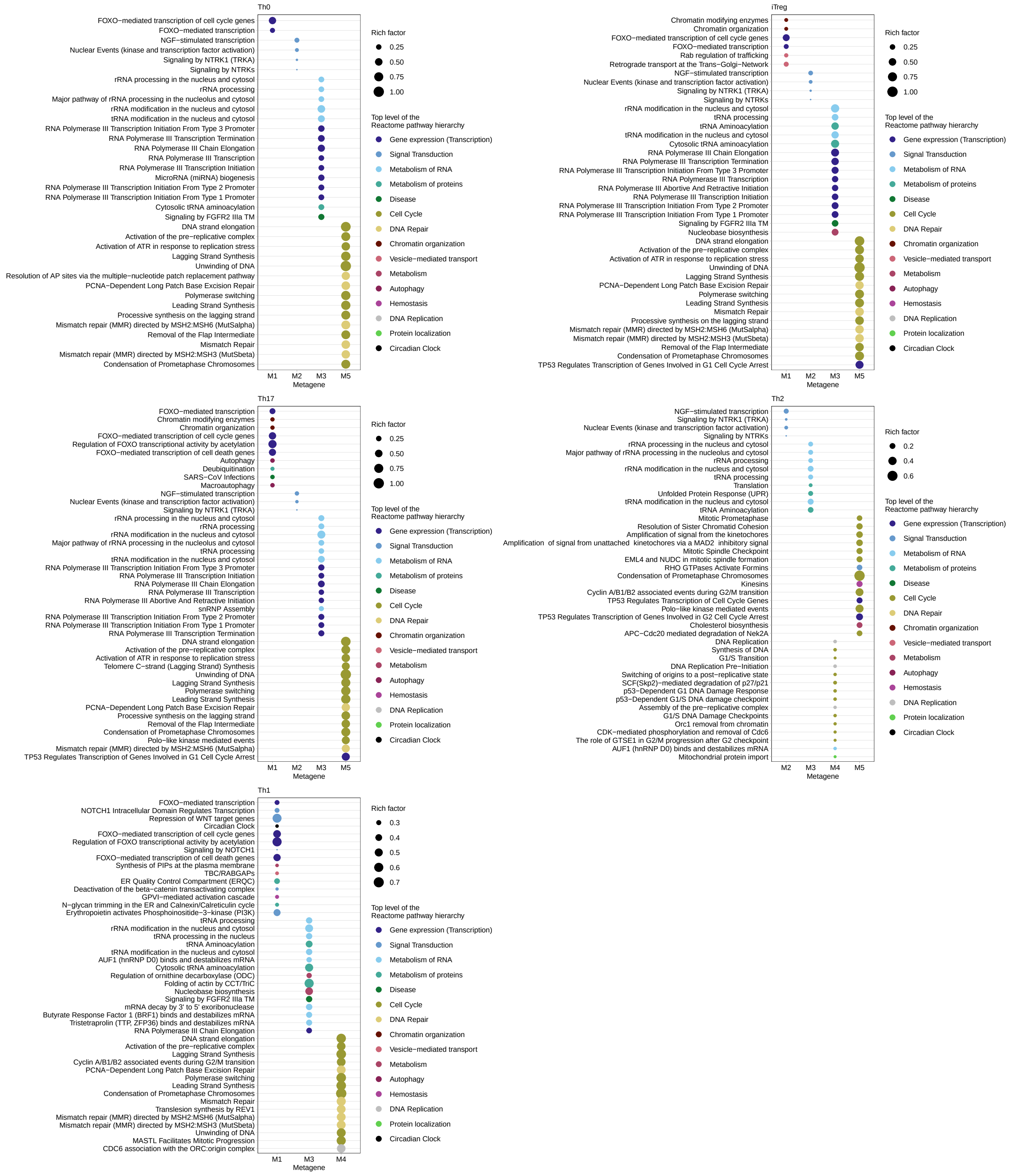

Fig. S17: ***Discovery Set*: Enrichment analysis of metagene associated genes for each T-cell population.** The top 15 (sorted by rich factor) significantly (FDR <0.05) enriched in Reactome pathways of metagene-associated genes for each CD4$^+$ T-cell population. The dot size indicates the rich factor, which is the number of metagene associated genes in the pathway divided by the number of background genes in that pathway. Colors depicted the uppermost hierarchical level of the Reactome database.

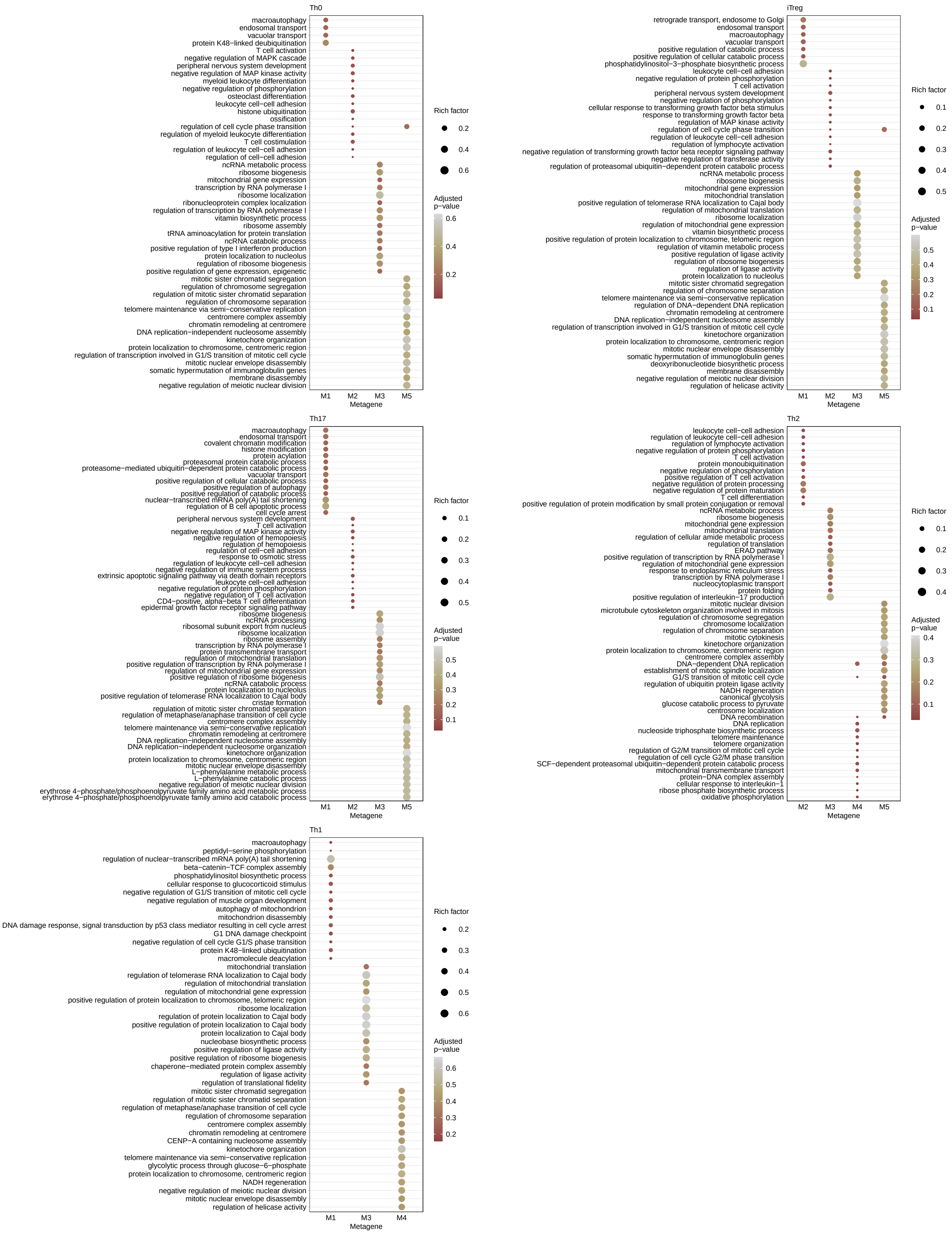

Fig. S18: ***Discovery Set*: Enrichment analysis of metagene associated genes for each T-cell population.** The top 15 (sorted by rich factor) significantly (FDR <0.05) enriched in Gene-Ontology terms (bottom) of metagene associated genes for each CD4$^+$ T-cell population. The dot size indicates the rich factor, which is the number of metagene associated genes in the term divided by the number of background genes in that term.

##### 5.1.3 Metagene landscape (combined pattern matrix)

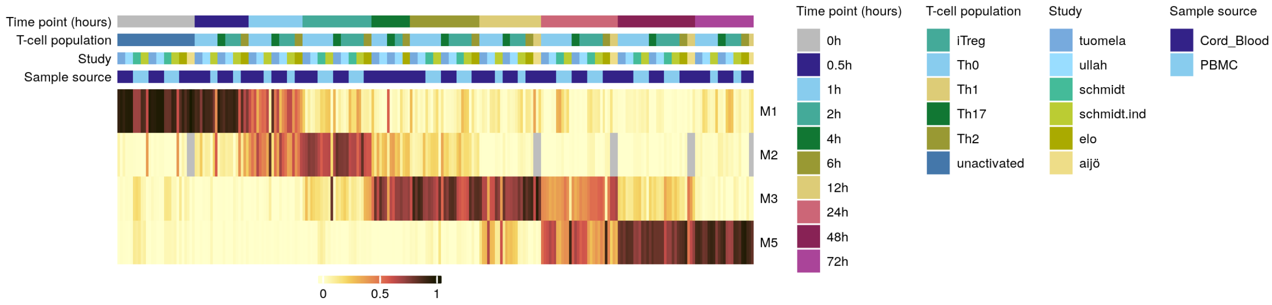

Fig. S19: ***Discovery Set*: Heatmap of the concatenated pattern matrix**. The pattern matrix $H$ was obtained from the factorization that achieved the lowest approximation error across 200 runs of the algorithm from Brunet et al. [26]. Factorization and subsequent rank determination was performed for each T-cell population (see methods). The matrices $H$ from each population were concatenated. For this, we used only temporally consistent metagenes across the T-cell populations. The columns in the combined matrix $H$ were scaled to sum to one and ordered by analysis time points. Since metagene 2 (expression peak at 2 hours) is not available for the Th1 population (only the analysis time points 12 to 72 hours are available), the values at Th1 were set to NA for metagene 2. The gray values in the combined matrix H represented the NA values. The combined pattern matrix was used to develop the Metagene landscape (see main methods).

##### 5.1.4 Enrichment analysis (Reactome) for gene sets from the consensus gene expression profiles

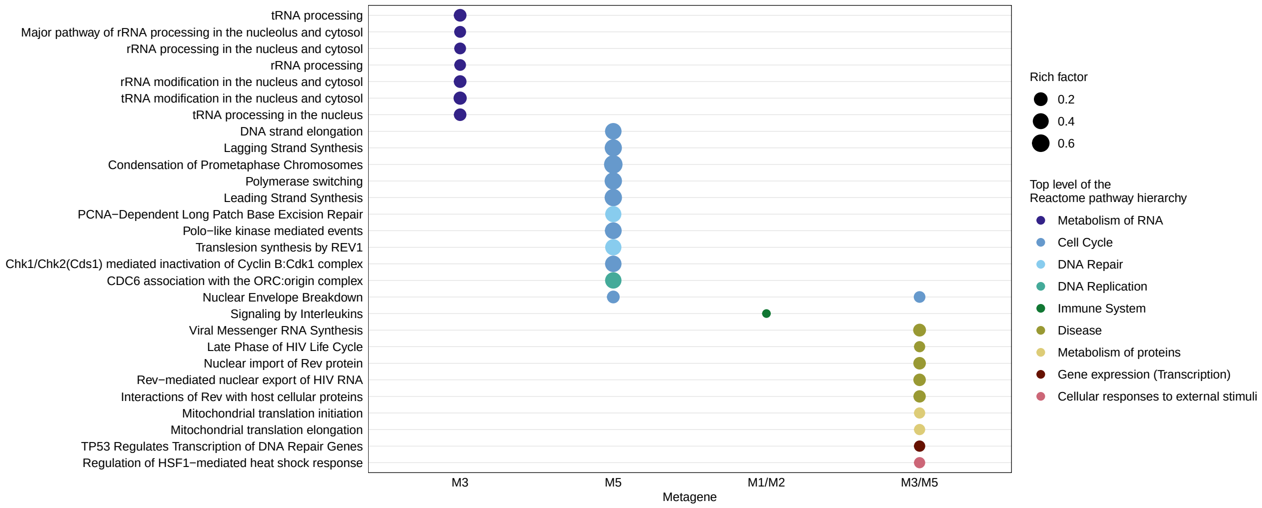

Fig. S20: ***Discovery Set*: Reactome enrichment analysis for genes sets from the consensus gene expression profiles.** Top 10 (sorted by rich factor) significantly enriched Reactome pathways (FDR <0.05) are shown. The dot size indicates the rich factor, which is the number of metagene associated genes in the pathway divided by the number of background genes in that pathway. Colors depicted the uppermost hierarchical level of Reactome database.

#### 5.2 *Verification Sets*

##### 5.2.1 Temporal profiles and highest ranked genes for metagenes

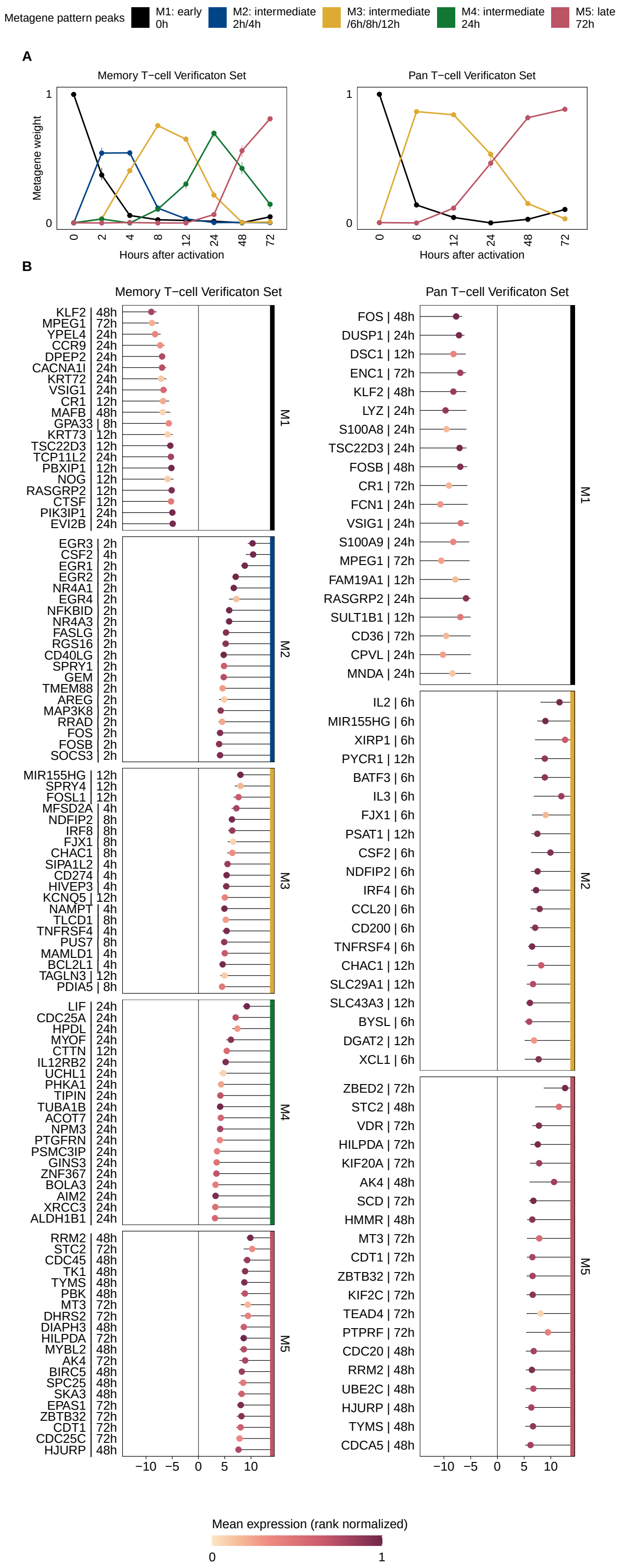

Fig. S21: ***Verification Sets*: Temporal profiles and 25 highest ranked genes for metagenes obtained from NMF.** **(A)** The pattern matrix for each T-cell population from the *Verification Sets* is shown as continuous profiles, with samples assigned to time points of activation. We scaled each column in the matrix to sum up to one. Dots depict median weights for all samples from identical analysis time points. Vertical lines represent interquartile ranges. We annotated and colored the metagenes based on their maximum median values across all analysis time points. The time point with maximum median value is depicted in the legend. **(B)** Top 25 genes associated with metagenes for each T-cell population. For each T-cell population and gene used for NMF, we used the highest absolute "confect" value estimated in the DGEA across all contrasts (e.g. 12h vs. 0h). Genes are ranked by "confect" values. Dots represent log2 fold changes for contrasts with the highest absolute "confect" value. The time point to the right of the gene represents the contrast with the highest absolute "confect" value. The Color of the dots correspond to rank normalized average expression values of the activation group in the contrast with the highest absolute "confect". The inner end of the horizontal line shows the "confect" value (inner confidence bound).

##### 5.2.2 Summary of consensus gene expression profiles verified for temporal consistency by the *Memory T-cell Verification Set*

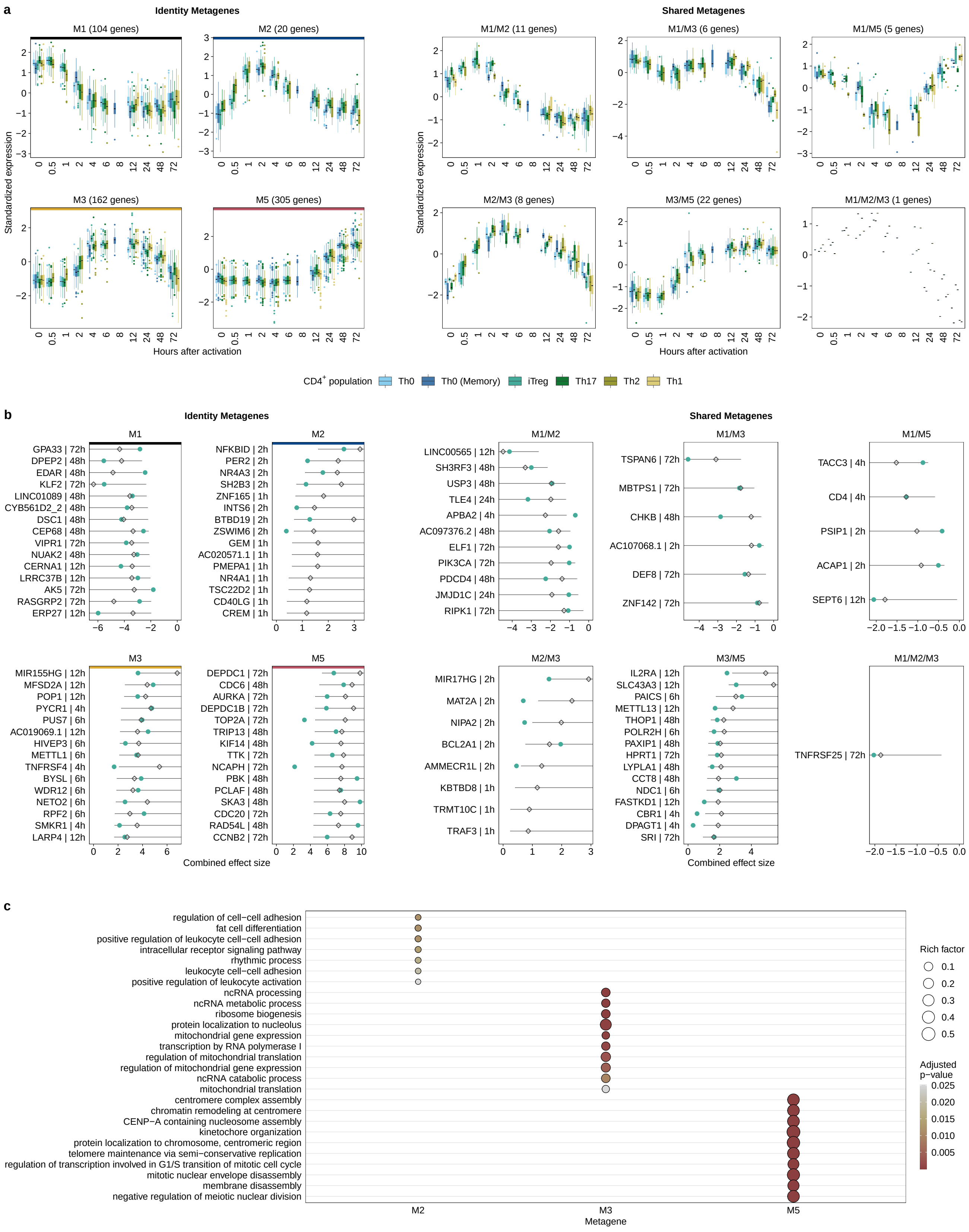

Fig. S22: **Consensus gene expression profiles verified for temporal consistency by the *Memory T-cell Verification Set*.** **(A)** We grouped the consensus expression profiles over the course by genes associated with identity and shared metagenes. Each boxplot represents one $CD4^{+}$ T-cell population from the *Discovery Set* and *Memory T-cell Verification Set*. The y-axis depicts standardized median expression of genes from samples with identical analysis time points. The number in parentheses represents the number of genes for the corresponding metagene. **(B)** Top 15 genes associated with identity and shared metagenes. For each gene belonging to the consistent metagenes, we used the highest absolute "confect" value estimated in the meta-analysis. Genes were ranked according to their absolute highest "confect" value. Diamonds represent combined effect size from the meta-analysis for the activation time point of the highest absolute "confect" value. The time point to the right of the gene represents the time point with the highest absolute "confect" value. The inner end the of horizontal line shows the "confect" value (inner confidence bound). The black dots represent the Hedges' g values of the genes from the *Memory T-cell Verification Set*. Since the analysis time point 6 hours is missing in the *Memory T-cell Verification Set*, the mean of the Hedge's g values from 4 hours and 8 hours was formed here. The time points of the largest "confect" value in the *Discovery Set* are not always present in the *Verification Set* and therefore cannot be mapped in the plot. **(C)** The top 10 (sorted by rich factor) significantly enriched GO terms of biological processes (FDR <0.05) of metagene associated genes identified by enrichment analysis. The dot size indicates the rich factor, which is the number of metagene associated genes in the GO term divided by the number of background genes of the term. Colors indicate adjusted p-values of significantly enriched GO terms For enrichment analysis of Reactome pathway see Figure S23.

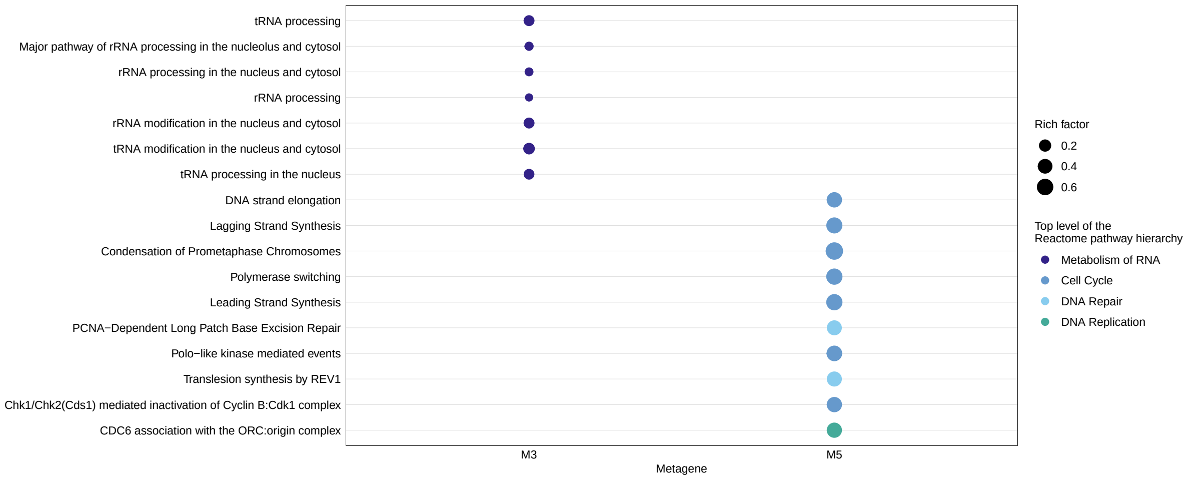

Fig. S23: **Reactome enrichment analysis for genes from the consensus gene expression profiles that passed the *Memory T-cell Verification Set*.** Top 10 (sorted by rich factor) significantly enriched Reactome pathways (FDR <0.05) are shown. The dot size indicates the rich factor, which is the number of metagene associated genes in the pathway divided by the number of background genes in that pathway. Colors depicted the uppermost hierarchical level of Reactome database.

##### 5.2.3 *Pan T-cell Verification Set*: Enrichment Analysis

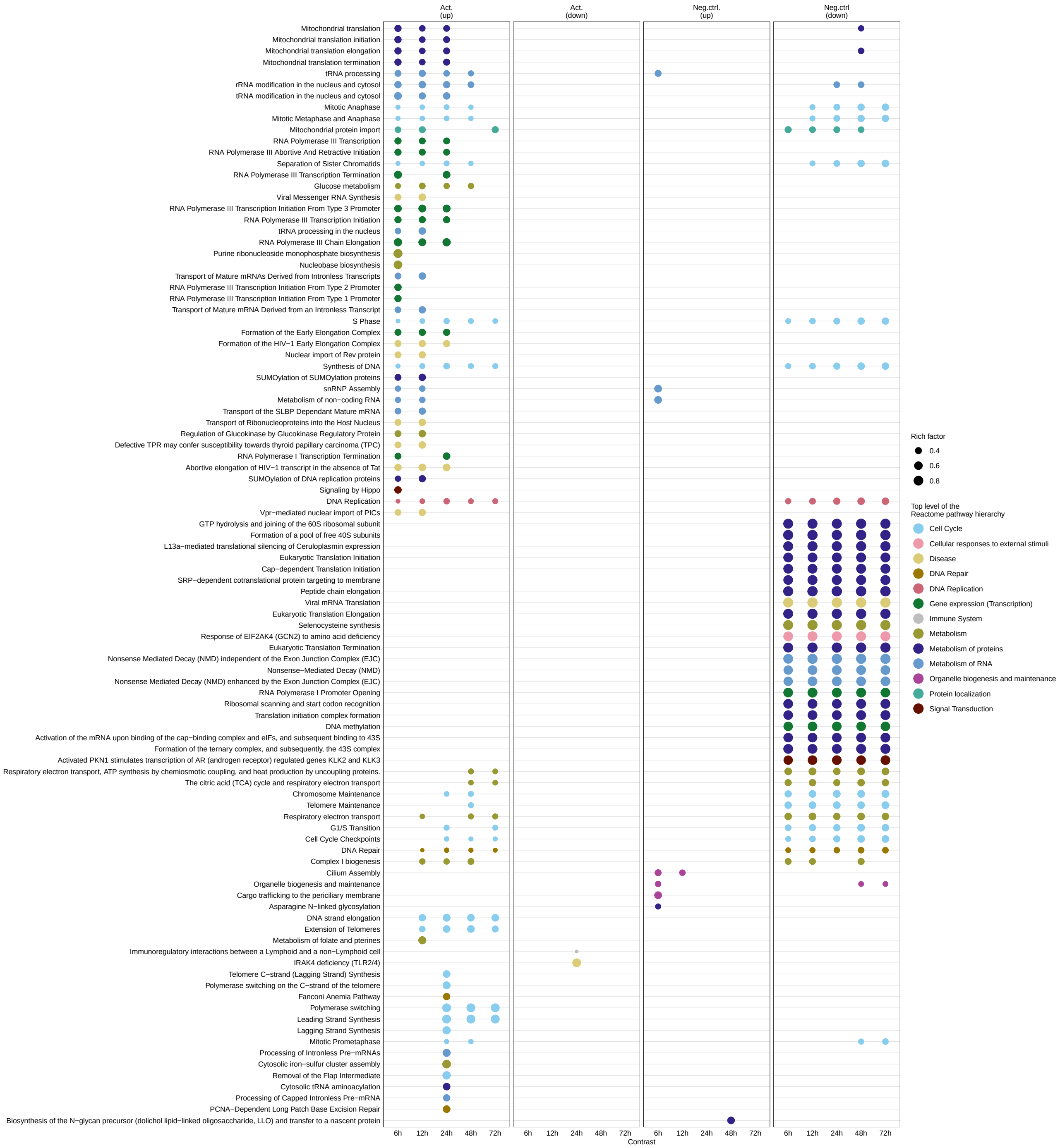

Fig. S24: **Reactome pathway enrichment analysis for DE genes from the *Pan T-cell Verification Set*.** For each contrast (6 to 72 hours of activation vs 0h and 6 to 72 hours without activation vs. 0h), we performed a gene enrichment analysis of DE genes (FDR <0.05). Act.(up/down) = DE genes that were upregulated/downregulated under activated conditions but not DE under unactivated conditions. Neg.ctrl.(up/down) = DE genes that were upregulated/downregulated under unactivated conditions (Genes can also be DE under activated conditions). Shown are the 10 most significant (FDR <0.05) enriched Reactome pathways per group (sorted by rich factor). The dot size indicates the rich factor which is the number of DE genes in the pathway divided by the number of background genes in the pathway. Colors depicted the uppermost hierarchical level of Reactome database.

##### 5.2.4 *Pan T-cell Verification Set*: DGEA

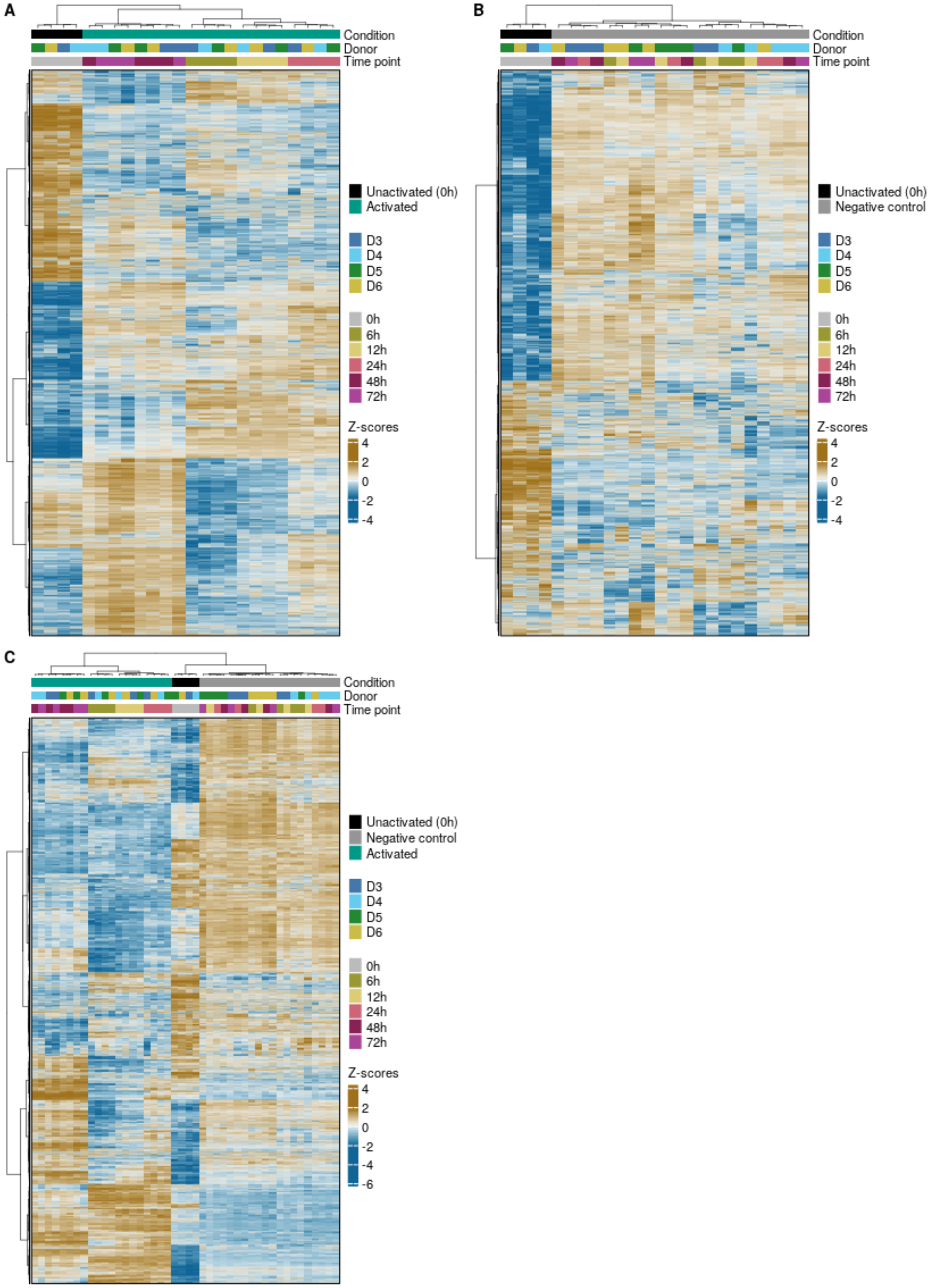

Fig. S25: **Hierarchical clustering of DE genes from the *Pan T-cell Verification Set*** in at least one contrast (6 to 72 hours vs. 0 hours) under **(A)** activation conditions, **(B)** unactivated conditions and **(C)** under both activated and unactivated conditions. Euclidean distance and Ward as clustering algorithm was applied to visualize similarity between samples. Each column represents a sample, each row represents a gene. CPM values for each gene were z-score standardized.

##### 5.2.5 *Pan T-cell Verification Set*: Verifying of the consensus gene expression profiles with unactivation kinetics (negative controls)

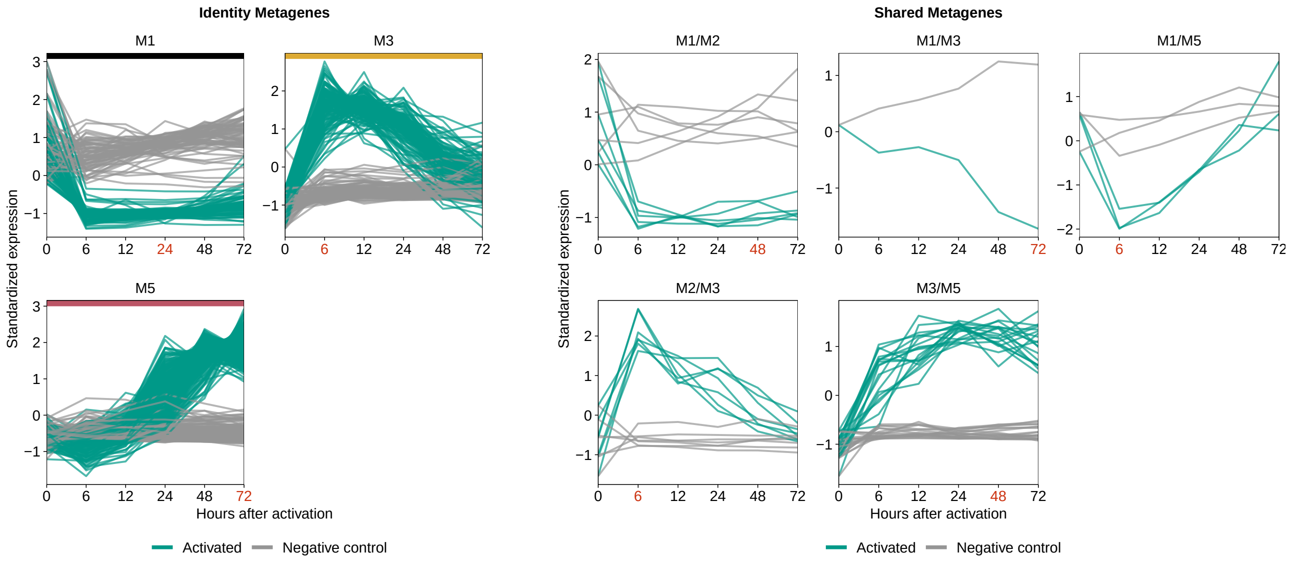

Fig. S26: ***Pan T-cell Verification Set*: Comparison of the expression profiles of the activation kinetics with the kinetics of the negative controls.** For the *Pan T-cell Verification Set*, the temporal expression pattern of genes from the consensus signatures that passed the 2 Verification Sets (activation and negative control kinetics) are shown. CPM values were z-score standardized. The red colored time points indicate the maximum centroid threshold (see main methods). Since only the analysis time points 6 to 72 hours are available, we could not make any conclusion about the time course of expression with respect to intermediate metagene 2 (expression peak after 2 hours). Therefore, we excluded metagene 2 from this analysis.

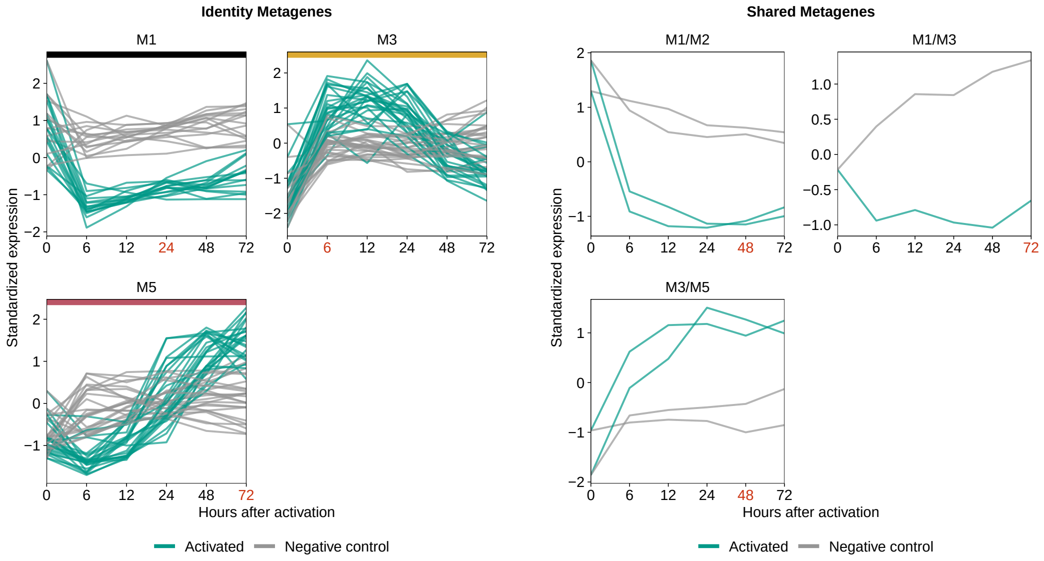

Fig. S27: For the *Pan T-cell Verification Set*, the temporal expression pattern of genes from the consensus signatures that passed the activation kinetics of the 2 *Verification sets* but **not** the kinetics of the negative controls is shown. CPM values were z-score standardized. Identity metagenes were depicted by a colored horizontal line

##### 5.2.6 Summary of consensus gene expression profiles verified for temporal consistency by the *Memory T-cell and Pan T-cell Verification Set*

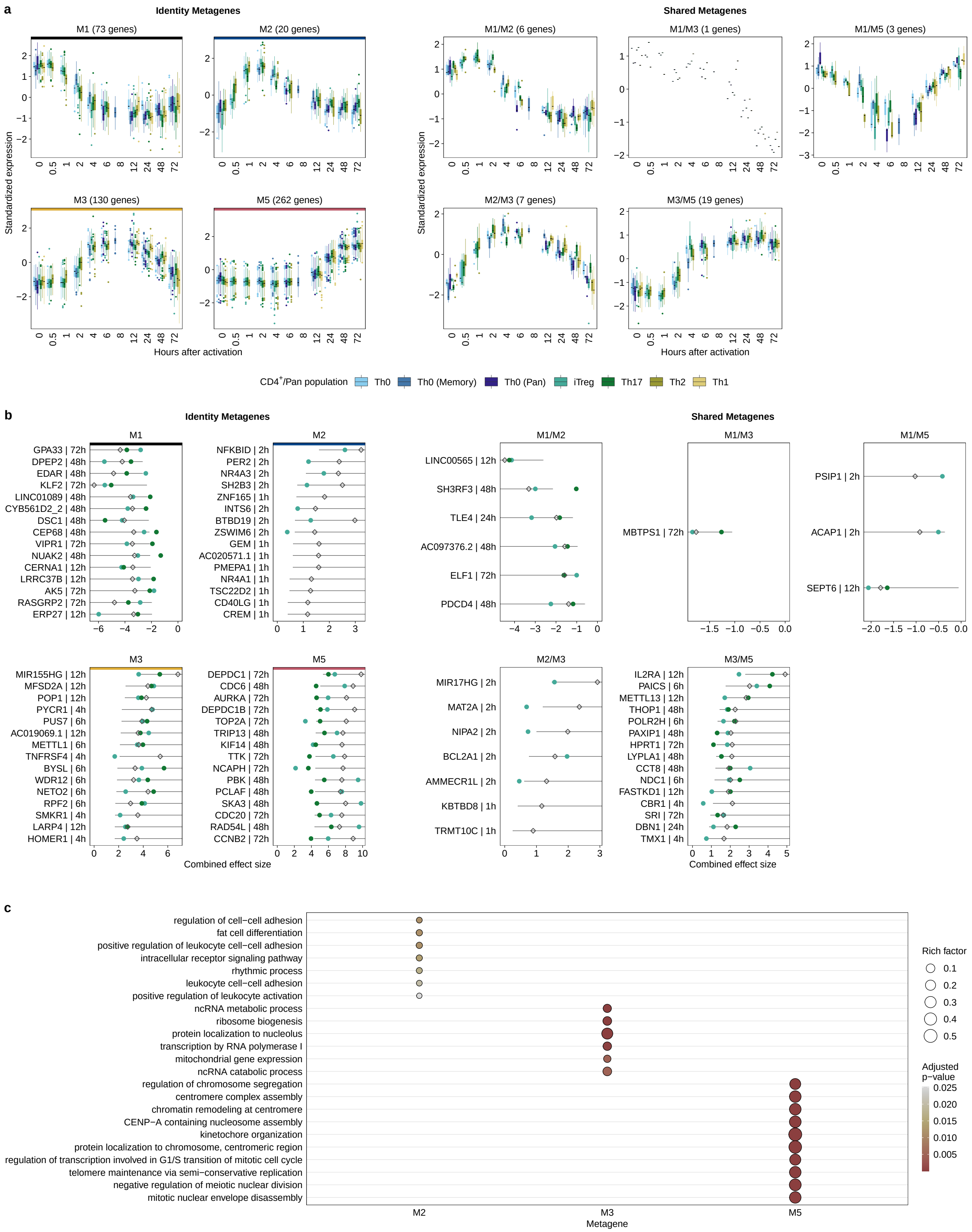

Fig. S28: **Consensus gene expression profiles verified for temporal consistency by both *Verification Sets* (activation and negative control kinetics)**. **(A)** We grouped the consensus expression profiles over the course by genes associated with identity and shared metagenes. Each boxplot represents one T-cell population from the *Discovery Set*, *Memory T-cell Verification Set* and *Pan T-cell Verification Set*. The y-axis depicted the standardized median expression of genes from samples with identical analysis time points. The number in parentheses represents the number of genes for the corresponding metagene. **(B)** Top 15 genes associated with identity and shared metagenes. For each gene belonging to the consistent metagenes, we used the highest absolute "confect" value estimated in the meta-analysis. Genes were ranked according to their absolute highest "confect" value. Diamonds represent combined effect size from the meta-analysis for the activation time point with the highest absolute "confect" value. The time point to the right of the gene represents the time point with the highest absolute "confect" value. The inner end the of horizontal line shows the "confect" value (inner confidence bound). The green and dark green dots represent the Hedges' g values of the genes from the *Memory T-cell Verification Set* and *Pan T-cell Verification Set*, respectively. Since the analysis time point 6 hours is missing in the *Memory T-cell Verification Set*, the mean of the Hedge's g values from 4 hours and 8 hours was formed here. The time points of the largest "confect" value in the *Discovery Set* are not always present in the *Verification Set* and therefore cannot be mapped in the plot. **(C)** The top 7 (sorted by rich factor) significantly enriched GO terms of biological processes (FDR <0.05) of metagene associated genes identified by enrichment analysis. The dot size indicates the rich factor, which is the number of metagene associated genes in the GO term divided by the number of background genes of the term. Colors indicate adjusted p-values of significantly enriched GO terms For enrichment analysis of Reactome pathway see Figure S29.

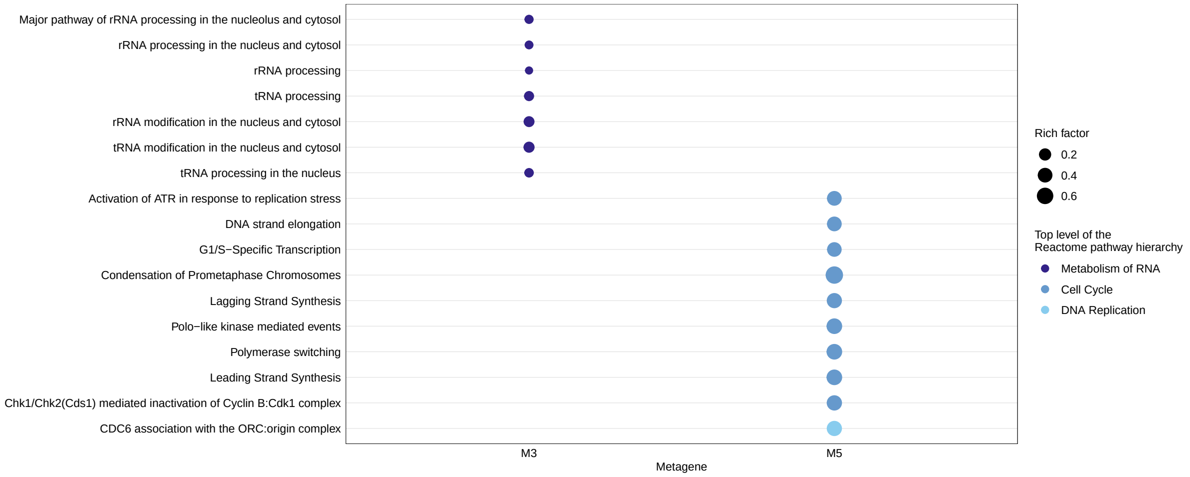

Fig. S29: **Reactome enrichment analysis for genes from the consensus gene expression profiles that passed both *Verification Sets*.** Top 10 (sorted by rich factor) significantly enriched Reactome pathways (FDR <0.05) are shown. The dot size indicates the rich factor, which is the number of metagene associated genes in the pathway divided by the number of background genes in that pathway. Colors depicted the uppermost hierarchical level of Reactome database.

### 6 Re-analysis of single-cell RNA sequencing data

#### 6.1 Known T-cell state/molecular: Cluster expression

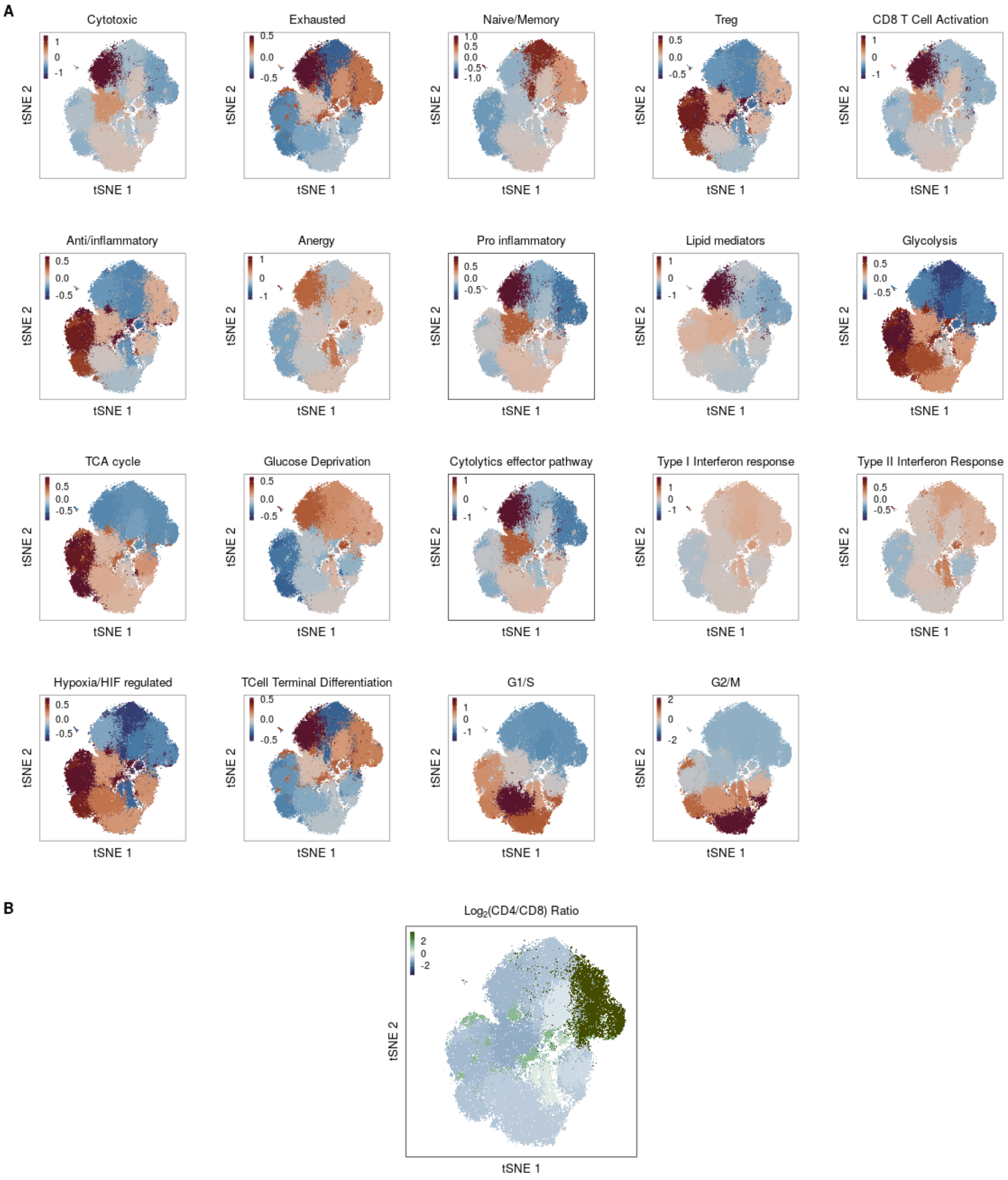

Fig. S30: **T-cell state/molecular mechanisms markers (A)** Standardized average expression of known T-cell state markers as well as markers for T-cell molecular mechanisms (see methods) for each cluster. **(B)** To obtain CD4/CD8 ratios for each cluster, we used the normalized expression values of $CD8^{-}CD4^{+}$ and $CD8^{+}CD4^{-}$ (average expression of CD8A and CD8B) in each cell. For both CD4 and CD8 we computed the average expression for each cluster. These values were then divided by the average for all cells. The log ratio of these values for CD4 and CD8 was then calculated. All cells in each cluster are labeled with the cluster's log-ratio.

#### 6.2 Known T-cell state/molecular mechanisms' marker: Differences in aggregated expression between patients with low- and high-grade ICANS

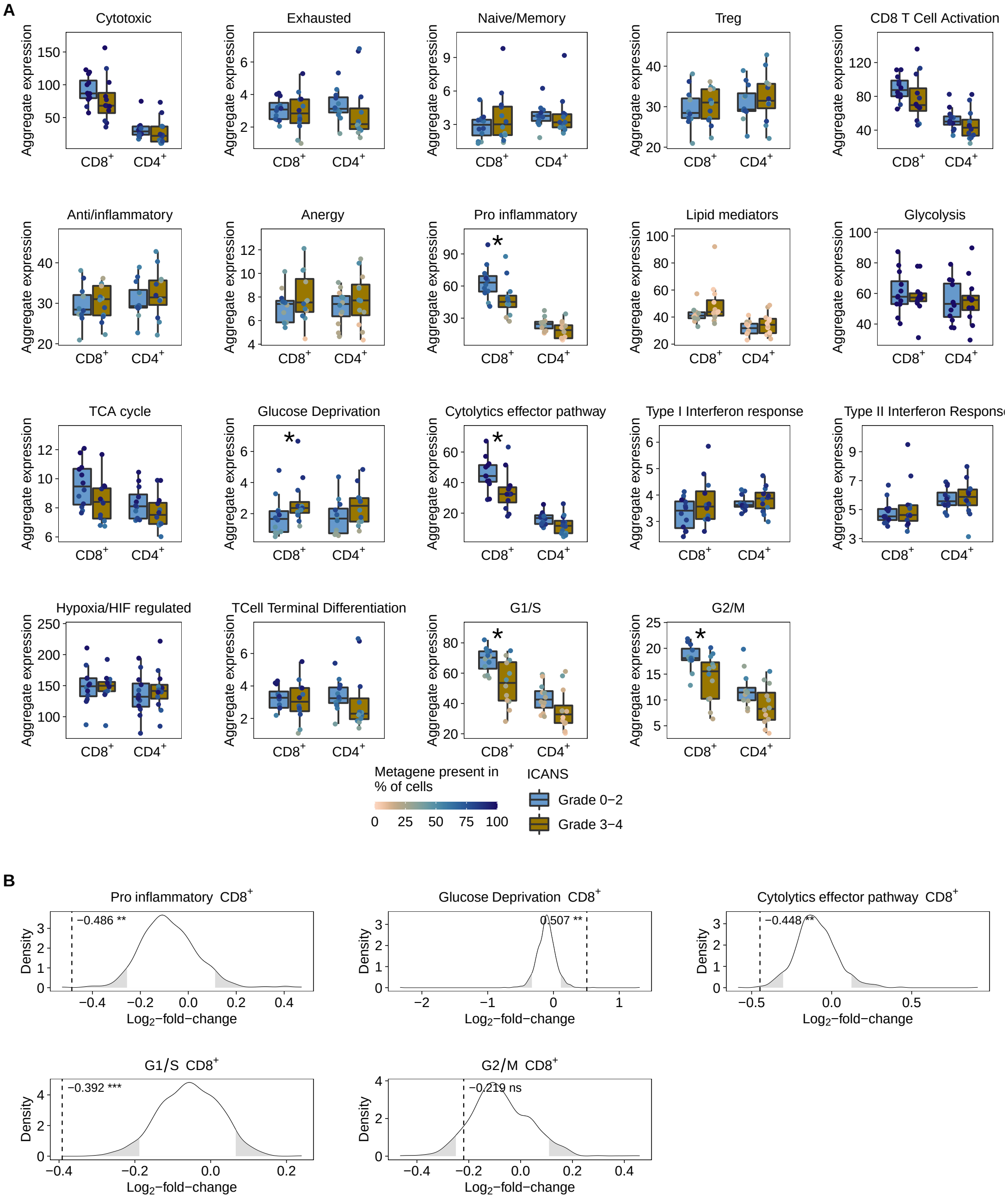

Fig. S31: **Differences in aggregate expression between patients with low- and high-grade ICANS using the Wilcoxon-rank-sum and permutation test for known T-cell state/molecular mechanisms markers. (A)** For each T-cell state/molecular mechanisms markers (denoted as gene sets) and patient, we calculated aggregated expression (summed average expression) for $CD8^+CD4^-$ and $CD8^-CD4^+$ cells. Differences in aggregated expression between patients with low- and high-grade ICANS were evaluated using the Wilcoxon rank-sum test. The colors of the dots in the boxplots indicate the percentage of cells for each patient in which the corresponding gene set is present. We considered a gene set as present in a cell if at least 25% of all associated genes had at least one UMI count. **(B)** For each gene set and T-cell population that was significant ($p < 0.05$) in the Wilcoxon rank-sum test, we generated a null distribution in order to confirm the results (see methods). Dashed vertical lines indicate median log2 fold change of aggregated expression between low- and high-grade ICANS patients of the gene set. Rejection regions (empirical p-value $< 0.05$) are highlighted in grey (* $p < 0.05$, ** $p < 0.01$, *** $p < 0.001$).

#### 6.3 Potential batch effects between aggregated expression and patient characteristics

| Metagene | Tcell_Lineage | Sex (p-value) | Lymphoma (p-value) | Age (p-value) |
|---|---|---|---|---|
| M5 | CD4-CD8+ | 0.264 | 0.850 | 0.211 |
| M5 | CD4+CD8- | 0.417 | 0.689 | 0.182 |
| M3 | CD4-CD8+ | 0.192 | 0.643 | 0.045 |
| M3 | CD4+CD8- | 0.172 | 0.367 | 0.242 |

Fig. S32: **Potential batch effects between aggregated expression and patient characteristics**. Only metagens with significant differences (p-value <0.05) in aggregated expression between patients with low-grade and high-grade ICANS were analysed. To determine the correlation between patient characteristics and metagenes/T cell lines, the following tests were performed: For sex, a Wilcoxon rank sum test was applied. The Kruskal-Wallis test was performed for the B-cell lymphoma subtypes (DLBCL=diffuse large B-cell lymphoma, PMBCL=primary mediastinal B-cell lymphoma, tFL= transformed follicular lymphoma). The significance for age was computed with Spearman's rank correlation using the `cor.test` `R` package.

#### 6.4 Aggregate expression between patients with low- and high-grade CRS or clinical outcome

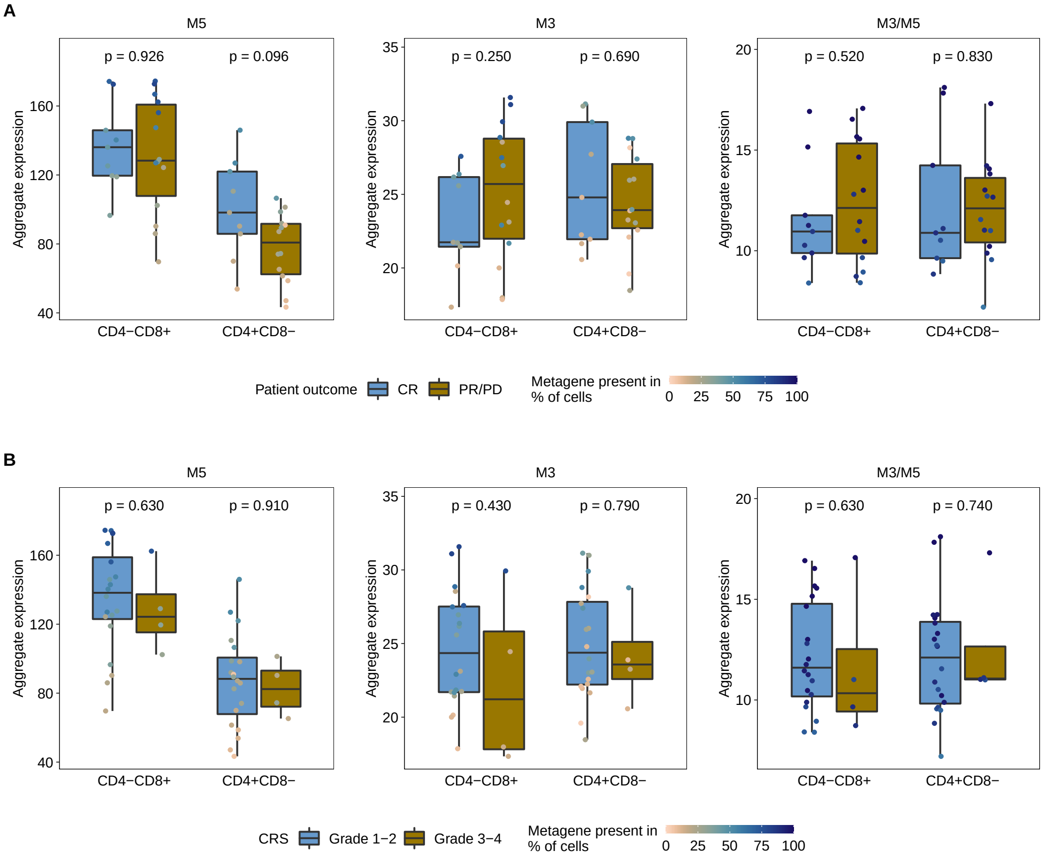

Fig. S33: **Test for differences in aggregate expression between patients with low- and high-grade CRS or clinical outcome (CR compared to PR/PD).** For each metagene and patient, we calculated aggregated expression (summed average expression) for $CD8^{+}CD4^{-}$ and $CD8^{-}CD4^{+}$ cells. Differences in aggregated expression were compared between patients with **(A)** complete response (CR) and partial response (PR)/progressive disease (PD), **(B)** low- and high-grade Cytokine Release Syndrome (CRS). Significance were calculated using the Wilcoxon rank sum test.
