## Additional File 2 for "A time-resolved meta-analysis of consensus gene expression profiles during human T-cell activation": additional_file_2.html


### Additional file 2

Author: Michael Rade


### 1 Consensus gene expression profiles

Fig. 4C of the publication: Consensus gene expression profiles for CD4+ T-cells from the Discovery Set. We grouped the consensus expression profiles over the course by genes associated with identity and shared metagenes. Each boxplot represents one CD4+ T-cell population from the Discovery Set. The y-axis depicts standardized median expression of genes from samples with identical analysis time points. The number in parentheses represents the number of genes for the corresponding metagene

#### 1.1 Summary

Shown are all genes from the consensus signatures that we identified based on the *Discovery Set*.

- **Mg:** Assignment of genes to metagenes based on the *Discovery Set*
- **Time\_point:** Time point of activation (hours) with the highest absolute “confect” value from the meta-analysis.
- **Effect:** Combined effect size from the meta-analysis for the time point with the highest absolute “confect” value.
- **P\_value:** Adjusted p-values for the combined effect size using the method of Benjamini-Hochberg.
- **Confect:** Confident effect sizes or “confect” for the combined effect size from the meta-analysis.
- **Passed\_V1:** Boolean value indicating whether the genes have passed the *Memory T-cell Verification set*.
- **Passed\_V2:** Boolean value indicating whether the genes have passed the *Pan T-cell Verification set*.
- **Mg\_V1/V2:** Assignment of genes to metagenes in the 2 *Verification Sets*. The following special cases are possible:
  - Due to the lack of analysis time points (0.5 to 4 hours) in the *Pan T-cell Verification set* (V2), we annotated metagene M2 in column “Mg\_V2” as “unknown”.
  - n.s = A gene was not significant (FDR <0.05) in any contrast of the differential gene expression analysis.
  - n.p = A gene was not available in den datasset (Only occurs in V1).
- **Passed\_V2\_Neg\_Ctrl:** Filtering step for time series negative controls (unactivated Pan T-cells after 6 to 72 hours) when comparing their gene expression profiles with the kinetics of the activated Pan T-cells. A boolean of “True” means that the gene passed the filtering step.
- **Pass\_Housekeeping\_Filter:** Genes that are present in the “Housekeeping and Reference Transcript Atlas database” (PMID: 32663312). A boolean of “True” means that the gene is not present in the database.

#### 1.2 Gene expression profiles

Shown are all genes from the consensus signatures that we identified based on the *Discovery Set*. If you click on the gene and press the “Backspace” key, you can then search for a gene by typing the symbol.

- **D:** Discovery set
- **V1: Th0 (Memory):** Memory T-cell Verification Set\*\*
- **V2: Th0 (Pan):** Pan T-cell Verification set\*\*.
- **V2: Neg. ctrl. (Pan):** Pan T-cell Verification set (negative controls)\*\*.

Search for a gene

The y-axis represents the median expression of genes from samples with identical analysis time points. The x-axis depicted quantile normalized count-per-million (CPM) for RNA-Seq and the normalized intensities (RMA normalization) for microarrays (Th1, Th2). Vertical lines represent the interquartile ranges.

### 2 Enrichment analysis

- Enrichment analysis test settings for the R/Bioconductor package clusterProfiler
  - P-value cutoff: 0.05 (cutoff for adjusted p-values)
  - Correction for multiple testing: Benjamini-Hochberg
  - Minimal size of genes annotated by term for testing: 10
  - Maximal size of genes annotated by term for testing: 500
  - Background gene set: All GENCODEv29 genes

- Meaning of BgRatio (M/N) and GeneRatio (k/n)
  - M = size of the gene-set (e.g. number of genes in the cell cycle term). More precisely, the total number of genes (ENTREZ Gene IDs) from the universe found in this gene-set.
  - N = size of all unique genes in the collection of gene-sets (e.g. the GO term collection for BPs). More precisely, the total number of genes (ENTREZ Gene IDs) from the universe found in this collection
  - k = size of overlap of genes with a specific gene-set (e.g. cell cycle term). Only unique ENTREZ Gene IDs were considered.
  - n = size of overlap of genes with all genes in the collection of gene-sets (e.g. the GO term collection for BPs). Only unique ENTREZ Gene IDs were considered.
  - The rich factor is k / M

#### 2.1 Discovery Set

All genes from the consensus signatures that we identified based on the *Discovery Set* were used for enrichment analysis.

##### 2.1.1 Gene Ontology term enrichment

Gene ontology category: Biological Process

##### 2.1.2 Reactome pathway enrichment

#### 2.2 Verified by the Memory T-cell Verification Set

All genes from the consensus signatures that we identified based on the *Discovery Set* and verified for temporal consistency by the *Memory T-cell Verification Set* were used for enrichment analysis.

##### 2.2.1 Gene Ontology term enrichment

Gene ontology category: Biological Process

##### 2.2.2 Reactome pathway enrichment

#### 2.3 Verified by both Verificaton Sets

All genes from the consensus signatures that we identified based on the *Discovery Set* and verified for temporal consistency by the *Memory* and *Pan T-cell Verification Set* were used for enrichment analysis.

##### 2.3.1 Gene Ontology term enrichment

Gene ontology category: Biological Process

##### 2.3.2 Reactome pathway enrichment
