## Additional File 3 for "A time-resolved meta-analysis of consensus gene expression profiles during human T-cell activation": Discovery_Set_GSE52260_qc.yaml

```
## Quality Data Assement for human RNA Seq Data
## Project ID: Tuomela-26967054-GSE52260-human-thp-th0-th17

cluster:
    default_job_quota: 100
    default_submit_options: "--cpus-per-task=#{CORES} --ntasks=1"
    default_pre_job_command: "echo 'Run Started!'"
    default_post_job_command: "echo 'Run Finished!'"

destination_path: work/uap_out/2018-MAVO-MCF/ag-indep/qc-Tuomela-26967054-GSE52260-thp-th0-th17/

constants:
    - &hisat2_genome_index_hg38 'references/hg38/hg38'
    - &genome_faidx_hg38 'references/hg38/hg38.fa.fai'
    - &hg38_ucscs_refflat 'annotations/hg38/ucsc/refFlat.txt'
    - &rseqc_gencode_v28 'work/2018-MAVO-MCF/annotations/hg38/gencode_v28/gencode.v28.annotation.basic.bed'
    - &bowtie2_genome_index_human_rRNA 'references/human_rRNA/human_rRNA'
    - &bowtie2_genome_index_rnammer 'references/rRNA/rnammer-1.2'
    - &fastq_screen 'src/fastq_screen/fastq_screen'
    - &fastq_screen_config 'src/fastq_screen/fastq_screen.conf'
    - &scs 'helper/collect_scs.py'

steps:
## >>>>>>>>>>>>>>>>>>>>>>>>>>>>>>>>>>>>>>>>>>>>>>>>>>>>>>>>>>>>>>>>>>>>>>>>>>>>>>>>>
## LOAD TRUNCATED READS (AFTER ADAPTERREMOVAL)
## >>>>>>>>>>>>>>>>>>>>>>>>>>>>>>>>>>>>>>>>>>>>>>>>>>>>>>>>>>>>>>>>>>>>>>>>>>>>>>>>>
    fq_trimmed (raw_file_source):
        pattern: work/uap_out/2018-MAVO-MCF/ag-indep/main-Tuomela-26967054-GSE52260-thp-th0-th17/adapterremoval/GSM*/*.truncated.fastq.gz
        group: (.*)\.truncated.fastq.gz

## >>>>>>>>>>>>>>>>>>>>>>>>>>>>>>>>>>>>>>>>>>>>>>>>>>>>>>>>>>>>>>>>>>>>>>>>>>>>>>>>>
## LOAD SORTED BAM
## >>>>>>>>>>>>>>>>>>>>>>>>>>>>>>>>>>>>>>>>>>>>>>>>>>>>>>>>>>>>>>>>>>>>>>>>>>>>>>>>>
    sorted_by_chr (raw_file_source):
        pattern: work/uap_out/2018-MAVO-MCF/ag-indep/main-Tuomela-26967054-GSE52260-thp-th0-th17/sort_by_chr_hg38/*/*sorted.bam
        group: (.*)\.sorted.bam

    bam_index (raw_file_source):
        pattern: work/uap_out/2018-MAVO-MCF/ag-indep/main-Tuomela-26967054-GSE52260-thp-th0-th17/bam_index/*/*sorted.bam
        group: (.*)\.sorted.bam

## >>>>>>>>>>>>>>>>>>>>>>>>>>>>>>>>>>>>>>>>>>>>>>>>>>>>>>>>>>>>>>>>>>>>>>>>>>>>>>>>>
## FASTQC (AFTER ADAPTERREMOVAL)
## >>>>>>>>>>>>>>>>>>>>>>>>>>>>>>>>>>>>>>>>>>>>>>>>>>>>>>>>>>>>>>>>>>>>>>>>>>>>>>>>>
    fastqc_after_trimming (fastqc):
        _depends: [fq_trimmed]
        _connect:
            in/first_read: fq_trimmed/raw
            in/second_read: empty

## >>>>>>>>>>>>>>>>>>>>>>>>>>>>>>>>>>>>>>>>>>>>>>>>>>>>>>>>>>>>>>>>>>>>>>>>>>>>>>>>>
## FASTQ SCREEN WITH SUBSAMPLES (READ 1)
## >>>>>>>>>>>>>>>>>>>>>>>>>>>>>>>>>>>>>>>>>>>>>>>>>>>>>>>>>>>>>>>>>>>>>>>>>>>>>>>>>
    fastqscreen (fastq_screen):
        _cluster_submit_options: "--cpus-per-task=8"
        _depends: [fq_trimmed]
        _connect:
            in/first_read: [fq_trimmed/raw]
        subset: 1000000
        config: *fastq_screen_config
        cores: 8

## >>>>>>>>>>>>>>>>>>>>>>>>>>>>>>>>>>>>>>>>>>>>>>>>>>>>>>>>>>>>>>>>>>>>>>>>>>>>>>>>>
## RSeQC AND COLLECTRNASEQMETRICS(PICARD TOOLS)
## >>>>>>>>>>>>>>>>>>>>>>>>>>>>>>>>>>>>>>>>>>>>>>>>>>>>>>>>>>>>>>>>>>>>>>>>>>>>>>>>>
    rseqc (rseqc): 
        _depends: bam_index
        _connect:
            in/alignments: bam_index/raw
        reference: *rseqc_gencode_v28
        treatAs: 'single'

    picard_collectRnaMetrics (picard_collectrnaseqmetrics):
        _depends: sorted_by_chr
        _connect:
            in/alignments: sorted_by_chr/raw
        REF_FLAT: *hg38_ucscs_refflat
        STRAND_SPECIFICITY: NONE

    picard_markDuplicates (picard_markduplicates):
        _cluster_submit_options: "--cpus-per-task=2"
        _depends: sorted_by_chr
        _connect:
            in/alignments: sorted_by_chr/raw

## >>>>>>>>>>>>>>>>>>>>>>>>>>>>>>>>>>>>>>>>>>>>>>>>>>>>>>>>>>>>>>>>>>>>>>>>>>>>>>>>>
## SUCESSIVE SCREENING FOR RRNA (READ 1)
## >>>>>>>>>>>>>>>>>>>>>>>>>>>>>>>>>>>>>>>>>>>>>>>>>>>>>>>>>>>>>>>>>>>>>>>>>>>>>>>>>
    ## 1) Align against RNAmmer 1.2
    successive_screen_bowtie2_rnammer (bowtie2):
        _cluster_submit_options: "--cpus-per-task=4 --mem 43G"
        _depends: [fq_trimmed]
        _connect:
            in/first_read: [fq_trimmed/raw]
            in/second_read: 'empty'
        index: *bowtie2_genome_index_rnammer
        unaligned: true
        cores: 4

    ## 2) Align unmapped reads against human rRNA
    successive_screen_bowtie2_human_rRNA (bowtie2):
        _cluster_submit_options: "--cpus-per-task=4 --mem 43G"
        _depends: successive_screen_bowtie2_rnammer
        _connect:
            in/first_read: successive_screen_bowtie2_rnammer/unaligned
            in/second_read: 'empty'
        index: *bowtie2_genome_index_human_rRNA
        unaligned: true
        cores: 4

    ## 3) Count rRNA subsets orientation information
    successive_screen_bowtie2_human_rRNA_count (count_rRNA):
        _depends: successive_screen_bowtie2_human_rRNA

    successive_screen_hisat2_hg38 (hisat2):
        _cluster_submit_options: "--cpus-per-task=4 --mem 43G"
        _depends: successive_screen_bowtie2_human_rRNA
        _connect:
            in/first_read: successive_screen_bowtie2_human_rRNA/unaligned
            in/second_read: 'empty'
        index: *hisat2_genome_index_hg38
        un-gz: false
        fr: false
        rf: false
        ff: false
        rna-strandness: U
        cores: 4
        q: true

    scs (collect_scs):
        _depends:
            - successive_screen_bowtie2_rnammer
            - successive_screen_bowtie2_human_rRNA_count
            - successive_screen_hisat2_hg38
        _connect:
            in/scs_metrics: 
                - successive_screen_bowtie2_rnammer/log_stderr
                - successive_screen_bowtie2_human_rRNA_count/report_rRNA
                - successive_screen_hisat2_hg38/log_stderr
        types:
            - rnammer
            - rRNA
            - hg38
        rrna-aln-pos: '2'
        library-type: 'unstranded'

tools:
## >>>>>>>>>>>>>>>>>>>>>>>>>>>>>>>>>>>>>>>>>>>>>>>>>>>>>>>>>>>>>>>>>>>>>>>>>>>>>>>>>
## External Tools #
## >>>>>>>>>>>>>>>>>>>>>>>>>>>>>>>>>>>>>>>>>>>>>>>>>>>>>>>>>>>>>>>>>>>>>>>>>>>>>>>>>
    preseq:
        path: 'preseq'
        module_load: 'lmod/libexec/lmod python load preseq/2.0.3-foss-2018b'
        module_unload: 'lmod/libexec/lmod python unload preseq/2.0.3-foss-2018b'
        exit_code: 0
    bowtie2:
        path: 'bowtie2'
        module_load: 'lmod/libexec/lmod python load Bowtie2/2.3.4.2-foss-2018b'
        module_unload: 'lmod/libexec/lmod python unload Bowtie2/2.3.4.2-foss-2018b'
        get_version: '--version'
        exit_code: 0

    fastq-sample:
        path: 'fastq-sample'
        get_version: '--version'
        module_load: 'lmod/libexec/lmod python load fastq-tools/0.8-foss-2018b'
        module_unload: 'lmod/libexec/lmod python unload fastq-tools/0.8-foss-2018b'

    fastqc:
        path: 'fastqc'
        get_version: '--version'
        module_load: 'lmod/libexec/lmod python load FastQC/0.11.8-Java-1.8'
        module_unload: 'lmod/libexec/lmod python unload FastQC/0.11.8-Java-1.8'

    fastx_reverse_complement:
        path: 'fastx_reverse_complement'
        get_version: '-h'
        module_load: 'lmod/libexec/lmod python load FASTX-Toolkit/0.0.14-foss-2018b'
        module_unload: 'lmod/libexec/lmod python unload FASTX-Toolkit/0.0.14-foss-2018b'
        exit_code: 1

    feature_counts:
        path: 'featureCounts'
        module_load: 'lmod/libexec/lmod python load Subread/1.5.2-foss-2018b'
        module_unload: 'lmod/libexec/lmod python unload Subread/1.5.2-foss-2018b'
        get_version: '-v'
        exit_code: 0

    hisat2:
        path: 'hisat2'
        get_version: '--version'
        module_load: 'lmod/libexec/lmod python load HISAT2/2.1.0-foss-2018b'
        module_unload: 'lmod/libexec/lmod python unload HISAT2/2.1.0-foss-2018b'

    htseq-count:
        path: 'htseq-count'
        module_load: 'lmod/libexec/lmod python load HTSeq/0.11.2-foss-2018b-Python-3.6.6'
        module_unload: 'lmod/libexec/lmod python unload HTSeq/0.11.2-foss-2018b-Python-3.6.6'
        exit_code: 2

# >> RSEQC START
    infer_experiment.py:
        path: 'infer_experiment.py'
        get_version: '--version'
        module_load: 'lmod/libexec/lmod python load RSeQC/3.0.0-foss-2018b-Python-3.6.6'
        module_unload: 'lmod/libexec/lmod python unload RSeQC/3.0.0-foss-2018b-Python-3.6.6'
        exit_code: 0

    bam_stat.py:
        path: 'bam_stat.py'
        get_version: '--version'
        module_load: 'lmod/libexec/lmod python load RSeQC/3.0.0-foss-2018b-Python-3.6.6'
        module_unload: 'lmod/libexec/lmod python unload RSeQC/3.0.0-foss-2018b-Python-3.6.6'
        exit_code: 0

    read_distribution.py:
        path: 'read_distribution.py'
        module_load: 'lmod/libexec/lmod python load RSeQC/3.0.0-foss-2018b-Python-3.6.6'
        module_unload: 'lmod/libexec/lmod python unload RSeQC/3.0.0-foss-2018b-Python-3.6.6'
        get_version: '--version'
        exit_code: 0

    geneBody_coverage.py:
        path: 'geneBody_coverage.py'
        module_load: 'lmod/libexec/lmod python load RSeQC/3.0.0-foss-2018b-Python-3.6.6'
        module_unload: 'lmod/libexec/lmod python unload RSeQC/3.0.0-foss-2018b-Python-3.6.6'
        get_version: '--version'
        exit_code: 0

    inner_distance.py:
        path: 'inner_distance.py'
        module_load: 'lmod/libexec/lmod python load RSeQC/3.0.0-foss-2018b-Python-3.6.6'
        module_unload: 'lmod/libexec/lmod python unload RSeQC/3.0.0-foss-2018b-Python-3.6.6'
        get_version: '--version'
        exit_code: 0

    junction_annotation.py:
        path: 'junction_annotation.py'
        module_load: 'lmod/libexec/lmod python load RSeQC/3.0.0-foss-2018b-Python-3.6.6'
        module_unload: 'lmod/libexec/lmod python unload RSeQC/3.0.0-foss-2018b-Python-3.6.6'
        get_version: '--version'
        exit_code: 0

    junction_saturation.py:
        path: 'junction_saturation.py'
        module_load: 'lmod/libexec/lmod python load RSeQC/3.0.0-foss-2018b-Python-3.6.6'
        module_unload: 'lmod/libexec/lmod python unload RSeQC/3.0.0-foss-2018b-Python-3.6.6'
        get_version: '--version'
        exit_code: 0

    read_duplication.py:
        path: 'read_duplication.py'
        module_load: 'lmod/libexec/lmod python load RSeQC/3.0.0-foss-2018b-Python-3.6.6'
        module_unload: 'lmod/libexec/lmod python unload RSeQC/3.0.0-foss-2018b-Python-3.6.6'
        get_version: '--version'
        exit_code: 0

    read_GC.py:
        path: 'read_GC.py'
        module_load: 'lmod/libexec/lmod python load RSeQC/3.0.0-foss-2018b-Python-3.6.6'
        module_unload: 'lmod/libexec/lmod python unload RSeQC/3.0.0-foss-2018b-Python-3.6.6'
        get_version: '--version'
        exit_code: 0
# >> RSEQC END

    picard-tools:
        module_load: 'lmod/libexec/lmod python load picard/2.18.29-Java-1.8'
        module_unload: 'lmod/libexec/lmod python unload picard/2.18.29-Java-1.8'
        path: ['java',  '-jar',  'software/picard/2.18.29-Java-1.8/picard.jar']
        get_version: '-h'
        exit_code: 1

    samtools:
        path: 'samtools'
        module_load: 'lmod/libexec/lmod python load SAMtools/1.9-foss-2018b'
        module_unload: 'lmod/libexec/lmod python unload SAMtools/1.9-foss-2018b'
        get_version: '--version'
        exit_code: 0

## >>>>>>>>>>>>>>>>>>>>>>>>>>>>>>>>>>>>>>>>>>>>>>>>>>>>>>>>>>>>>>>>>>>>>>>>>>>>>>>>>
## CUSTOM AND MODIFIED TOOLS
## >>>>>>>>>>>>>>>>>>>>>>>>>>>>>>>>>>>>>>>>>>>>>>>>>>>>>>>>>>>>>>>>>>>>>>>>>>>>>>>>>
    collect_scs:
        path: *scs
        get_version: '--version'
        exit_code: 0

    fastq_screen:
        path: *fastq_screen
        get_version: '--version'
        exit_code: 0

## >>>>>>>>>>>>>>>>>>>>>>>>>>>>>>>>>>>>>>>>>>>>>>>>>>>>>>>>>>>>>>>>>>>>>>>>>>>>>>>>>
## Unix Tools #
## >>>>>>>>>>>>>>>>>>>>>>>>>>>>>>>>>>>>>>>>>>>>>>>>>>>>>>>>>>>>>>>>>>>>>>>>>>>>>>>>>
    awk:
        path: 'awk'
        get_version: '--version'

    cat:
        path: 'cat'
        get_version: '--version'
        exit_code: 0

    cp:
        path: 'cp'
        get_version: '--version'
        exit_code: 0

    cut:
        path: 'cut'
        get_version: '--version'
        exit_code: 0

    dd:
        path: 'dd'
        get_version: '--version'
        exit_code: 0

    echo:
        path: '/bin/echo'
        get_version: '--version'
        exit_code: 0

    grep:
        path: 'grep'
        get_version: '--version'
        exit_code: 0

    ln:
        path: 'ln'
        get_version: '--version'
        exit_code: 0

    mkdir:
        path: 'mkdir'
        get_version: '--version'
        exit_code: 0

    mv:
        path: 'mv'
        get_version: '--version'
        exit_code: 0

    mkfifo:
        path: 'mkfifo'
        get_version: '--version'
        exit_code: 0

    pigz: 
        path: 'pigz'
        get_version: '--version'
        exit_code: 0

    pwd:
        path: 'pwd'
        get_version: '--version'
        exit_code: 0

    rm:
        path: 'rm'
        get_version: '--version'
        exit_code: 0

    sort:
        path: 'sort'
        get_version: '--version'
        exit_code: 0

    tar:
        path: 'tar'
        get_version: '--version'
        exit_code: 0

    uniq:
        path: 'uniq'
        get_version: '--version'
        exit_code: 0
```
