## Additional File 3 for "A time-resolved meta-analysis of consensus gene expression profiles during human T-cell activation": Discovery_Set_GSE90569_main.yaml

```
## Secondary Analysis for human RNA Seq Data
## Project ID: Ullah-29466736-GSE90569-human-thp-th0-iTreg

cluster:
    default_job_quota: 100
    default_submit_options: "--cpus-per-task=#{CORES} --ntasks=1"
    default_pre_job_command: "echo 'Run Started!'"
    default_post_job_command: "echo 'Run Finished!'"

destination_path: work/uap_out/2018-MAVO-MCF/ag-indep/main-Ullah-29466736-GSE90569-thp-th0-iTreg/

constants:
    - &hisat2_genome_index_hg38 "references/hg38/hg38"
    - &ref_gencode_v29_hg38 "work/2018-MAVO-MCF/annotations/hg38/gencode_v29/gencode.v29.annotation.gtf"
    - &ref_gencode_v32 "databases/gencode/homo_sapiens/v32/gtf/gencode.v32.annotation.gtf"

steps:
## >>>>>>>>>>>>>>>>>>>>>>>>>>>>>>>>>>>>>>>>>>>>>>>>>>>>>>>>>>>>>>>>>>>>>>>>>>>>>>>>>
## LOAD RAWDATA (FASTQ)
## >>>>>>>>>>>>>>>>>>>>>>>>>>>>>>>>>>>>>>>>>>>>>>>>>>>>>>>>>>>>>>>>>>>>>>>>>>>>>>>>>
    fastq_files_pe (run_folder_source):
        path: data/RNA-Seq/2018-MAVO-MCF/Ullah-29466736-GSE90569-thp-th0-iTreg
        first_read: _1
        second_read: _2
        paired_end: True

## >>>>>>>>>>>>>>>>>>>>>>>>>>>>>>>>>>>>>>>>>>>>>>>>>>>>>>>>>>>>>>>>>>>>>>>>>>>>>>>>>
## IDENTIFY ADAPTER
## >>>>>>>>>>>>>>>>>>>>>>>>>>>>>>>>>>>>>>>>>>>>>>>>>>>>>>>>>>>>>>>>>>>>>>>>>>>>>>>>>
    idendify_adapters (identify_adapters):
        _depends: [fastq_files_pe]

## >>>>>>>>>>>>>>>>>>>>>>>>>>>>>>>>>>>>>>>>>>>>>>>>>>>>>>>>>>>>>>>>>>>>>>>>>>>>>>>>>
## TRIMMING
## >>>>>>>>>>>>>>>>>>>>>>>>>>>>>>>>>>>>>>>>>>>>>>>>>>>>>>>>>>>>>>>>>>>>>>>>>>>>>>>>>
    adapterremoval (adapterremoval):
        _cluster_submit_options: "--cpus-per-task=8 --mem 43G"
        _depends: [fastq_files_pe]
        treatAs: paired
        adapter1: AGATCGGAAGAGCACACGTCT
        adapter2: AGATCGGAAGAGCGTCGTGTA
        collapse: False
        trimns: True
        trimqualities: True
        minquality: 20
        threads: 8
        minlength: 30

## >>>>>>>>>>>>>>>>>>>>>>>>>>>>>>>>>>>>>>>>>>>>>>>>>>>>>>>>>>>>>>>>>>>>>>>>>>>>>>>>>
## HISAT2 and HTSeq
## >>>>>>>>>>>>>>>>>>>>>>>>>>>>>>>>>>>>>>>>>>>>>>>>>>>>>>>>>>>>>>>>>>>>>>>>>>>>>>>>>
    hisat2_hg38 (hisat2):
        _cluster_submit_options: "--cpus-per-task=8 --ntasks=1 --time 0-09:59 --mem 43G"
        _depends: adapterremoval
        _connect:
            in/first_read: adapterremoval/pair1.truncated
            in/second_read: adapterremoval/pair2.truncated
        index: *hisat2_genome_index_hg38
        fr: true
        rf: false
        ff: false
        rna-strandness: U
        cores: 8
        q: true

    sort_by_chr_hg38 (samtools_sort):
        _cluster_submit_options: "--cpus-per-task=8 --ntasks=1 --time 0-09:59 --mem 43G"
        _depends: hisat2_hg38
        sort-by-name: False
        O: 'BAM'

    bam_index (samtools_index):
        _cluster_submit_options: "--cpus-per-task=8 --ntasks=1 --time 0-09:59 --mem 43G"
        _depends: sort_by_chr_hg38
        index_type: 'bai'

    htseq_hisat2 (htseq_count):
        _cluster_submit_options: "--cpus-per-task=2 --ntasks=1 --time 5-24:59"
        _depends: [sort_by_chr_hg38]
        _connect:
            in/alignments: [sort_by_chr_hg38/alignments]
        stranded: 'no'
        mode: intersection-strict
        type: exon
        idattr: gene_id
        order: pos
        feature-file: *ref_gencode_v29_hg38

tools:
## >>>>>>>>>>>>>>>>>>>>>>>>>>>>>>>>>>>>>>>>>>>>>>>>>>>>>>>>>>>>>>>>>>>>>>>>>>>>>>>>>
## External Tools #
## >>>>>>>>>>>>>>>>>>>>>>>>>>>>>>>>>>>>>>>>>>>>>>>>>>>>>>>>>>>>>>>>>>>>>>>>>>>>>>>>>
    adapterremoval:
        path: 'AdapterRemoval'
        get_version: '--version'
        module_load: 'lmod/lmod/libexec/lmod python load AdapterRemoval/2.2.2-foss-2018b'
        module_unload: 'lmod/lmod/libexec/lmod python unload AdapterRemoval/2.2.2-foss-2018b'

    hisat2:
        path: 'hisat2'
        get_version: '--version'
        module_load: 'lmod/lmod/libexec/lmod python load HISAT2/2.1.0-foss-2018b'
        module_unload: 'lmod/lmod/libexec/lmod python unload HISAT2/2.1.0-foss-2018b'

    htseq-count:
        path: 'htseq-count'
        module_load: 'lmod/lmod/libexec/lmod python load HTSeq/0.11.2-foss-2018b-Python-3.6.6'
        module_unload: 'lmod/lmod/libexec/lmod python unload HTSeq/0.11.2-foss-2018b-Python-3.6.6'
        get_version: '--help'
        exit_code: 0

    samtools:
        path: 'samtools'
        module_load: 'lmod/lmod/libexec/lmod python load SAMtools/1.9-foss-2018b'
        module_unload: 'lmod/lmod/libexec/lmod python unload SAMtools/1.9-foss-2018b'
        get_version: '--version'
        exit_code: 0

    stringtie:
        path: 'stringtie'
        get_version: '--version'
        module_load: 'lmod/lmod/libexec/lmod python load StringTie/2.0.3-GCCcore-7.3.0'
        module_unload: 'lmod/lmod/libexec/lmod python unload StringTie/2.0.3-GCCcore-7.3.0'
        exit_code: 0 

## >>>>>>>>>>>>>>>>>>>>>>>>>>>>>>>>>>>>>>>>>>>>>>>>>>>>>>>>>>>>>>>>>>>>>>>>>>>>>>>>>
## Unix Tools
## >>>>>>>>>>>>>>>>>>>>>>>>>>>>>>>>>>>>>>>>>>>>>>>>>>>>>>>>>>>>>>>>>>>>>>>>>>>>>>>>>
    cat:
        path: 'cat'
        get_version: '--version'
        exit_code: 0

    dd:
        path: 'dd'
        get_version: '--version'
        exit_code: 0

    mv:
        path: 'mv'
        get_version: '--version'
        exit_code: 0

    ln:
        path: 'ln'
        get_version: '--version'
        exit_code: 0

    pigz:
        path: 'pigz'
        get_version: '--version'
        exit_code: 0

    pwd:
        path: 'pwd'
        get_version: '--version'
        exit_code: 0

    mkdir:
        path: 'mkdir'
        get_version: '--version'
        exit_code: 0
```
