## Additional File 3 for "A time-resolved meta-analysis of consensus gene expression profiles during human T-cell activation": Verification_Set_Pan_Tcells_main.yaml

```
## Secondary Analysis for human RNA Seq Data
## Project ID: Verification Set 2

cluster:
    default_job_quota: 100
    default_submit_options: "--cpus-per-task=#{CORES}"
    default_pre_job_command: "echo 'Run Started!'"
    default_post_job_command: "echo 'Run Finished!'"

destination_path: 'work/uap_out/2018-MAVO-MCF/ag-indep/main-izi-200312_D00594_0106_BCD1RVANXX-thp-th0/'

constants:
    - &hisat2_genome_index_hg38 'genomes/mammalia/rodentia/homo_sapiens/hg38/indices/hisat2/hg38'
    - &genome_faidx_hg38 'genomes/mammalia/rodentia/homo_sapiens/hg38/fasta/hg38.fa.fai'
    - &gencode_v29_hg38 'metadata/gencode/homo_sapiens/v29/gtf/gencode.v29.annotation.gtf'

steps:
## >>>>>>>>>>>>>>>>>>>>>>>>>>>>>>>>>>>>>>>>>>>>>>>>>>>>>>>>>>>>>>>>>>>>>>>>>>>>>>>>>
## LOAD RAWDATA (FASTQ)
## >>>>>>>>>>>>>>>>>>>>>>>>>>>>>>>>>>>>>>>>>>>>>>>>>>>>>>>>>>>>>>>>>>>>>>>>>>>>>>>>>
    fq (fastq_source):
        pattern: HiSeq/200312_D00594_0106_BCD1RVANXX_KEEP/fastq/MyCellFight/*.gz
        group: (RIB\d+)_S.*fastq.gz
        first_read: '_R1'
        second_read: '_R2'
        paired_end: True

## >>>>>>>>>>>>>>>>>>>>>>>>>>>>>>>>>>>>>>>>>>>>>>>>>>>>>>>>>>>>>>>>>>>>>>>>>>>>>>>>>
## ADAPTER TRIMMING
## >>>>>>>>>>>>>>>>>>>>>>>>>>>>>>>>>>>>>>>>>>>>>>>>>>>>>>>>>>>>>>>>>>>>>>>>>>>>>>>>>
    adapterremoval (adapterremoval):
        _volatile: no
        _cluster_submit_options: "--cpus-per-task=8"
        _depends: fq
        treatAs: paired
        adapter1: AGATCGGAAGAGCACACGTCTGAACTCCAGTCA
        adapter2: AGATCGGAAGAGCGTCGTGTAGGGAAAGAGTGT
        collapse: False
        trimns: True
        trimqualities: True
        minquality: 20
        cores: 8
        threads: 8
        minlength: 30
        seed: 1234

## >>>>>>>>>>>>>>>>>>>>>>>>>>>>>>>>>>>>>>>>>>>>>>>>>>>>>>>>>>>>>>>>>>>>>>>>>>>>>>>>>
## ALIGNMENT
## >>>>>>>>>>>>>>>>>>>>>>>>>>>>>>>>>>>>>>>>>>>>>>>>>>>>>>>>>>>>>>>>>>>>>>>>>>>>>>>>>
    hisat2 (hisat2):
        _volatile: no
        _cluster_submit_options: "--cpus-per-task=8 --mem=60G"
        _depends: adapterremoval
        _connect:
            in/first_read: adapterremoval/pair1.truncated
            in/second_read: adapterremoval/pair2.truncated
        index: *hisat2_genome_index_hg38
        fr: true
        rf: false
        ff: false
        rna-strandness: R
        cores: 8
        q: true

## >>>>>>>>>>>>>>>>>>>>>>>>>>>>>>>>>>>>>>>>>>>>>>>>>>>>>>>>>>>>>>>>>>>>>>>>>>>>>>>>>
## SORTING BY NAME
## >>>>>>>>>>>>>>>>>>>>>>>>>>>>>>>>>>>>>>>>>>>>>>>>>>>>>>>>>>>>>>>>>>>>>>>>>>>>>>>>>
    sort_by_name (sam_to_sorted_bam):
        _cluster_submit_options: "--cpus-per-task=8"
        _depends: hisat2
        genome-faidx: *genome_faidx_hg38
        sort-by-name: True

## >>>>>>>>>>>>>>>>>>>>>>>>>>>>>>>>>>>>>>>>>>>>>>>>>>>>>>>>>>>>>>>>>>>>>>>>>>>>>>>>>
## SORTING BY CHR
## >>>>>>>>>>>>>>>>>>>>>>>>>>>>>>>>>>>>>>>>>>>>>>>>>>>>>>>>>>>>>>>>>>>>>>>>>>>>>>>>>
    sort_by_chr (sam_to_sorted_bam):
        _cluster_submit_options: "--cpus-per-task=8"
        _depends: hisat2
        genome-faidx: *genome_faidx_hg38
        sort-by-name: False

## >>>>>>>>>>>>>>>>>>>>>>>>>>>>>>>>>>>>>>>>>>>>>>>>>>>>>>>>>>>>>>>>>>>>>>>>>>>>>>>>>
## COUNTING
## >>>>>>>>>>>>>>>>>>>>>>>>>>>>>>>>>>>>>>>>>>>>>>>>>>>>>>>>>>>>>>>>>>>>>>>>>>>>>>>>>
    htseq_count (htseq_count):
        _cluster_submit_options: "--cpus-per-task=2"
        _depends: sort_by_name
        stranded: 'reverse'
        mode: intersection-strict
        type: exon
        idattr: gene_id
        order: name
        feature-file: *gencode_v29_hg38

tools:
## >>>>>>>>>>>>>>>>>>>>>>>>>>>>>>>>>>>>>>>>>>>>>>>>>>>>>>>>>>>>>>>>>>>>>>>>>>>>>>>>>
## External Tools
## >>>>>>>>>>>>>>>>>>>>>>>>>>>>>>>>>>>>>>>>>>>>>>>>>>>>>>>>>>>>>>>>>>>>>>>>>>>>>>>>>
    adapterremoval:
        path: 'AdapterRemoval'
        module_load: 'lmod/lmod/libexec/lmod python load AdapterRemoval/2.3.1-foss-2018b'
        module_unload: 'lmod/lmod/libexec/lmod python unload AdapterRemoval/2.3.1-foss-2018b'
        get_version: '--version'
        exit_code: 0

    hisat2:
        path: 'hisat2'
        module_load: 'lmod/lmod/libexec/lmod python load HISAT2/2.1.0-foss-2018b'
        module_unload: 'lmod/lmod/libexec/lmod python unload HISAT2/2.1.0-foss-2018b'
        get_version: '--version'
        exit_code: 0

    htseq-count:
        path: 'htseq-count'
        module_load: 'lmod/lmod/libexec/lmod python load HTSeq/0.11.2-foss-2018b-Python-3.6.6'
        module_unload: 'lmod/lmod/libexec/lmod python unload HTSeq/0.11.2-foss-2018b-Python-3.6.6'
        get_version: '--help'
        exit_code: 0

    samtools:
        path: 'samtools'
        module_load: 'lmod/lmod/libexec/lmod python load SAMtools/1.9-foss-2018b'
        module_unload: 'lmod/lmod/libexec/lmod python unload SAMtools/1.9-foss-2018b'
        get_version: '--version'
        exit_code: 0

## >>>>>>>>>>>>>>>>>>>>>>>>>>>>>>>>>>>>>>>>>>>>>>>>>>>>>>>>>>>>>>>>>>>>>>>>>>>>>>>>>
## Unix Tools
## >>>>>>>>>>>>>>>>>>>>>>>>>>>>>>>>>>>>>>>>>>>>>>>>>>>>>>>>>>>>>>>>>>>>>>>>>>>>>>>>>
    cat:
        path: 'cat'
        get_version: '--version'
        exit_code: 0

    dd:
        path: 'dd'
        get_version: '--version'
        exit_code: 0

    mv:
        path: 'mv'
        get_version: '--version'
        exit_code: 0

    ln:
        path: 'ln'
        get_version: '--version'
        exit_code: 0

    pigz:
        path: 'pigz'
        get_version: '--version'
        exit_code: 0

    pwd:
        path: 'pwd'
        get_version: '--version'
        exit_code: 0

    mkdir:
        path: 'mkdir'
        get_version: '--version'
        exit_code: 0
```
