## Additional File 3 for "A time-resolved meta-analysis of consensus gene expression profiles during human T-cell activation": Verification_Set_Pan_Tcells_qc.yaml

```
## Quality Data Assement for human RNA Seq Data
## Project ID: Verification Set 2

cluster:
    default_job_quota: 100
    default_submit_options: "--cpus-per-task=#{CORES}"
    default_pre_job_command: "echo 'Run Started!'"
    default_post_job_command: "echo 'Run Finished!'"

destination_path: 'work/uap_out/2018-MAVO-MCF/ag-indep/qc-izi-200312_D00594_0106_BCD1RVANXX-thp-th0/'

constants:
    - &hisat2_genome_index_hg38 'genomes/mammalia/rodentia/homo_sapiens/hg38/indices/hisat2/hg38'
    - &hg38_ucscs_refflat 'metadata/ucsc/hg38/databases/refFlat.txt'
    - &rseqc_gencode_v31 'metadata/ucsc/hg38/tables/wgEncodeGencodeBasicV31/wgEncodeGencodeBasicV31.bed'
    - &bowtie2_genome_index_human_rRNA 'metadata/ncbi/refseq/human-rRNA/indices/human_rRNA'
    - &bowtie2_genome_index_rnammer 'metadata/rnammer/indices/rnammer-1.2'
    - &fastq_screen_config 'src/fastq_screen.conf'
    - &scs 'src/collect_scs.py'

steps:
## >>>>>>>>>>>>>>>>>>>>>>>>>>>>>>>>>>>>>>>>>>>>>>>>>>>>>>>>>>>>>>>>>>>>>>>>>>>>>>>>>
## LOAD TRUNCATED READS (AFTER ADAPTERREMOVAL)
## >>>>>>>>>>>>>>>>>>>>>>>>>>>>>>>>>>>>>>>>>>>>>>>>>>>>>>>>>>>>>>>>>>>>>>>>>>>>>>>>>
    fq_trimmed_R1 (raw_file_source):
        pattern: 'work/uap_out/2018-MAVO-MCF/ag-indep/main-izi-200312_D00594_0106_BCD1RVANXX-thp-th0/adapterremoval/*/*.pair1.truncated.fastq.gz'
        group: '(.*)\.pair1.truncated.fastq.gz'

    fq_trimmed_R2 (raw_file_source):
        pattern: 'work/uap_out/2018-MAVO-MCF/ag-indep/main-izi-200312_D00594_0106_BCD1RVANXX-thp-th0/adapterremoval/*/*.pair2.truncated.fastq.gz'
        group: '(.*)\.pair2.truncated.fastq.gz'

## >>>>>>>>>>>>>>>>>>>>>>>>>>>>>>>>>>>>>>>>>>>>>>>>>>>>>>>>>>>>>>>>>>>>>>>>>>>>>>>>>
## LOAD SORTED BAM
## >>>>>>>>>>>>>>>>>>>>>>>>>>>>>>>>>>>>>>>>>>>>>>>>>>>>>>>>>>>>>>>>>>>>>>>>>>>>>>>>>
    sorted_by_chr (raw_file_source):
        pattern: 'work/uap_out/2018-MAVO-MCF/ag-indep/main-izi-200312_D00594_0106_BCD1RVANXX-thp-th0/sort_by_chr/*/*.sorted.bam'
        group: '(.*)\.sorted.bam'

    bam_index (samtools_index):
        _cluster_submit_options: "--cpus-per-task=4"
        _depends: sorted_by_chr
        _connect:
            in/alignments: sorted_by_chr/raw
        index_type: 'bai'

## >>>>>>>>>>>>>>>>>>>>>>>>>>>>>>>>>>>>>>>>>>>>>>>>>>>>>>>>>>>>>>>>>>>>>>>>>>>>>>>>>
##  REVERSE COMPLEMENT
## >>>>>>>>>>>>>>>>>>>>>>>>>>>>>>>>>>>>>>>>>>>>>>>>>>>>>>>>>>>>>>>>>>>>>>>>>>>>>>>>>
    revcom (fastx_reverse_complement):
        _depends: [fq_trimmed_R2]
        _cluster_submit_options: "--cpus-per-task=1"
        _connect:
            in/fastx: fq_trimmed_R2/raw
        prefix: 'revcom_R2'

## >>>>>>>>>>>>>>>>>>>>>>>>>>>>>>>>>>>>>>>>>>>>>>>>>>>>>>>>>>>>>>>>>>>>>>>>>>>>>>>>>
## Preseq
## >>>>>>>>>>>>>>>>>>>>>>>>>>>>>>>>>>>>>>>>>>>>>>>>>>>>>>>>>>>>>>>>>>>>>>>>>>>>>>>>>
    preseq (preseq_complexity_curve):
            _cluster_submit_options: "--cpus-per-task=4 --mem 40G"
            _depends: sorted_by_chr
            _connect:
                in/alignments: sorted_by_chr/raw
            pe: false

## >>>>>>>>>>>>>>>>>>>>>>>>>>>>>>>>>>>>>>>>>>>>>>>>>>>>>>>>>>>>>>>>>>>>>>>>>>>>>>>>>
## FASTQC (AFTER ADAPTERREMOVAL)
## >>>>>>>>>>>>>>>>>>>>>>>>>>>>>>>>>>>>>>>>>>>>>>>>>>>>>>>>>>>>>>>>>>>>>>>>>>>>>>>>>
    fastqc_after_trimming (fastqc):
      _cluster_submit_options: "--cpus-per-task=1"
      _depends: [fq_trimmed_R1, fq_trimmed_R2]
      _connect:
        in/first_read: fq_trimmed_R1/raw
        in/second_read: fq_trimmed_R2/raw

## >>>>>>>>>>>>>>>>>>>>>>>>>>>>>>>>>>>>>>>>>>>>>>>>>>>>>>>>>>>>>>>>>>>>>>>>>>>>>>>>>
## FASTQ SCREEN
## >>>>>>>>>>>>>>>>>>>>>>>>>>>>>>>>>>>>>>>>>>>>>>>>>>>>>>>>>>>>>>>>>>>>>>>>>>>>>>>>>
    fastqscreen (fastq_screen):
        _cluster_submit_options: "--cpus-per-task=8 --mem 40G --exclude=ribnode018"
        _depends: [fq_trimmed_R1, revcom]
        _connect:
            in/first_read: [fq_trimmed_R1/raw, revcom/fastx]
        subset: 0
        config: *fastq_screen_config
        cores: 8

## >>>>>>>>>>>>>>>>>>>>>>>>>>>>>>>>>>>>>>>>>>>>>>>>>>>>>>>>>>>>>>>>>>>>>>>>>>>>>>>>>
## RSeQC AND COLLECTRNASEQMETRICS(PICARD TOOLS)
## >>>>>>>>>>>>>>>>>>>>>>>>>>>>>>>>>>>>>>>>>>>>>>>>>>>>>>>>>>>>>>>>>>>>>>>>>>>>>>>>>
    rseqc (rseqc):
        _cluster_submit_options: "--cpus-per-task=4 --mem 60G"
        _depends: bam_index
        reference: *rseqc_gencode_v31
        treatAs: 'paired'

    picard_collectRnaMetrics (picard_collectrnaseqmetrics):
        _cluster_submit_options: "--cpus-per-task=1"
        _depends: sorted_by_chr
        _connect:
            in/alignments: sorted_by_chr/raw
        REF_FLAT: *hg38_ucscs_refflat
        STRAND_SPECIFICITY: SECOND_READ_TRANSCRIPTION_STRAND

    picard_markDuplicates (picard_markduplicates):
        _cluster_submit_options: "--cpus-per-task=8"
        _depends: sorted_by_chr
        _connect:
            in/alignments: sorted_by_chr/raw

## >>>>>>>>>>>>>>>>>>>>>>>>>>>>>>>>>>>>>>>>>>>>>>>>>>>>>>>>>>>>>>>>>>>>>>>>>>>>>>>>>
## DOWNSTREAM QC WITH SUBSAMPLES (1 MIO)
## >>>>>>>>>>>>>>>>>>>>>>>>>>>>>>>>>>>>>>>>>>>>>>>>>>>>>>>>>>>>>>>>>>>>>>>>>>>>>>>>>
    subsample (fastqsample):
        _cluster_submit_options: "--cpus-per-task=1"
        _depends: [fq_trimmed_R1, fq_trimmed_R2]
        _connect:
           in/first_read: fq_trimmed_R1/raw
           in/second_read: fq_trimmed_R2/raw
        n: 1000000

    subsample_revcom (fastx_reverse_complement):
      _depends: [subsample]
      _connect:
        in/fastx: subsample/second_read
      prefix: 'revcom'

## >>>>>>>>>>>>>>>>>>>>>>>>>>>>>>>>>>>>>>>>>>>>>>>>>>>>>>>>>>>>>>>>>>>>>>>>>>>>>>>>>
## SUCESSIVE SCREENING FOR RRNA (READ 1)
## >>>>>>>>>>>>>>>>>>>>>>>>>>>>>>>>>>>>>>>>>>>>>>>>>>>>>>>>>>>>>>>>>>>>>>>>>>>>>>>>>
    ## 1) Align against RNAmmer 1.2
    successive_screen_bowtie2_rnammer (bowtie2):
        _cluster_submit_options: "--cpus-per-task=4 --mem 10G"
        _depends: [subsample, subsample_revcom]
        _connect:
            in/first_read: [subsample/first_read, subsample_revcom/fastx]
            in/second_read: 'empty'
        index: *bowtie2_genome_index_rnammer
        unaligned: true
        cores: 8

    ## 2) Align unmapped reads against human rRNA
    successive_screen_bowtie2_human_rRNA (bowtie2):
        _cluster_submit_options: "--cpus-per-task=4 --mem 10G"
        _depends: successive_screen_bowtie2_rnammer
        _connect:
            in/first_read: successive_screen_bowtie2_rnammer/unaligned
            in/second_read: 'empty'
        index: *bowtie2_genome_index_human_rRNA
        unaligned: true
        cores: 8

    ## 3) Count rRNA subsets orientation information
    successive_screen_bowtie2_human_rRNA_count (count_rRNA):
        _depends: successive_screen_bowtie2_human_rRNA

    successive_screen_hisat2 (hisat2):
        _cluster_submit_options: "--cpus-per-task=4 --mem 30G"
        _depends: successive_screen_bowtie2_human_rRNA
        _connect:
            in/first_read: successive_screen_bowtie2_human_rRNA/unaligned
            in/second_read: 'empty'
        index: *hisat2_genome_index_hg38
        un-gz: false
        fr: false
        rf: false
        ff: false
        rna-strandness: R # rna-strandness:  fr-firststrand ->  R, fr-secondstrand -> F 
        cores: 8
        q: true

    scs (collect_scs):
        _cluster_submit_options: "--cpus-per-task=1"
        _depends:
            - successive_screen_bowtie2_rnammer
            - successive_screen_bowtie2_human_rRNA_count
            - successive_screen_hisat2
        _connect:
            in/scs_metrics: 
                - successive_screen_bowtie2_rnammer/log_stderr
                - successive_screen_bowtie2_human_rRNA_count/report_rRNA
                - successive_screen_hisat2/log_stderr
        types:
            - rnammer
            - rRNA
            - hg38
        rrna-aln-pos: '2'
        library-type: 'firststranded'

tools:
## >>>>>>>>>>>>>>>>>>>>>>>>>>>>>>>>>>>>>>>>>>>>>>>>>>>>>>>>>>>>>>>>>>>>>>>>>>>>>>>>>
## External Tools #
## >>>>>>>>>>>>>>>>>>>>>>>>>>>>>>>>>>>>>>>>>>>>>>>>>>>>>>>>>>>>>>>>>>>>>>>>>>>>>>>>>
    bowtie2:
        path: 'bowtie2'
        module_load: 'lmod/lmod/libexec/lmod python load Bowtie2/2.3.4.2-foss-2018b'
        module_unload: 'lmod/lmod/libexec/lmod python unload Bowtie2/2.3.4.2-foss-2018b'
        get_version: '--version'
        exit_code: 0

    fastq-sample:
        path: 'fastq-sample'
        module_load: 'lmod/lmod/libexec/lmod python load fastq-tools/0.8-foss-2018b'
        module_unload: 'lmod/lmod/libexec/lmod python unload fastq-tools/0.8-foss-2018b'
        get_version: '--version'
        exit_code: 0

    fastqc:
        path: 'fastqc'
        module_load: 'lmod/lmod/libexec/lmod python load FastQC/0.11.8-Java-1.8'
        module_unload: 'lmod/lmod/libexec/lmod python unload FastQC/0.11.8-Java-1.8'
        get_version: '--version'
        exit_code: 0

    fastq_screen:
        path: 'fastq_screen'
        module_load: 'lmod/lmod/libexec/lmod python load FastQ_Screen/0.14.0-foss-2018b-Perl-5.28.0'
        module_unload: 'lmod/lmod/libexec/lmod python unload FastQ_Screen/0.14.0-foss-2018b-Perl-5.28.0'
        get_version: '--version'
        exit_code: 0

    fastx_reverse_complement:
        path: 'fastx_reverse_complement'
        module_load: 'lmod/lmod/libexec/lmod python load FASTX-Toolkit/0.0.14-foss-2018b'
        module_unload: 'lmod/lmod/libexec/lmod python unload FASTX-Toolkit/0.0.14-foss-2018b'
        get_version: '-h'
        exit_code: 1

    hisat2:
        path: 'hisat2'
        module_load: 'lmod/lmod/libexec/lmod python load HISAT2/2.1.0-foss-2018b'
        module_unload: 'lmod/lmod/libexec/lmod python unload HISAT2/2.1.0-foss-2018b'
        get_version: '--version'
        exit_code: 0


# >> RSEQC START
    infer_experiment.py:
        path: 'infer_experiment.py'
        module_load: 'lmod/lmod/libexec/lmod python load RSeQC/3.0.0-foss-2018b-Python-3.6.6'
        module_unload: 'lmod/lmod/libexec/lmod python unload RSeQC/3.0.0-foss-2018b-Python-3.6.6'
        get_version: '--version'
        exit_code: 0

    bam_stat.py:
        path: 'bam_stat.py'
        module_load: 'lmod/lmod/libexec/lmod python load RSeQC/3.0.0-foss-2018b-Python-3.6.6'
        module_unload: 'lmod/lmod/libexec/lmod python unload RSeQC/3.0.0-foss-2018b-Python-3.6.6'
        get_version: '--version'
        exit_code: 0

    read_distribution.py:
        path: 'read_distribution.py'
        module_load: 'lmod/lmod/libexec/lmod python load RSeQC/3.0.0-foss-2018b-Python-3.6.6'
        module_unload: 'lmod/lmod/libexec/lmod python unload RSeQC/3.0.0-foss-2018b-Python-3.6.6'
        get_version: '--version'
        exit_code: 0

    geneBody_coverage.py:
        path: 'geneBody_coverage.py'
        module_load: 'lmod/lmod/libexec/lmod python load RSeQC/3.0.0-foss-2018b-Python-3.6.6'
        module_unload: 'lmod/lmod/libexec/lmod python unload RSeQC/3.0.0-foss-2018b-Python-3.6.6'
        get_version: '--version'
        exit_code: 0

    inner_distance.py:
        path: 'inner_distance.py'
        module_load: 'lmod/lmod/libexec/lmod python load RSeQC/3.0.0-foss-2018b-Python-3.6.6'
        module_unload: 'lmod/lmod/libexec/lmod python unload RSeQC/3.0.0-foss-2018b-Python-3.6.6'
        get_version: '--version'
        exit_code: 0

    junction_annotation.py:
        path: 'junction_annotation.py'
        module_load: 'lmod/lmod/libexec/lmod python load RSeQC/3.0.0-foss-2018b-Python-3.6.6'
        module_unload: 'lmod/lmod/libexec/lmod python unload RSeQC/3.0.0-foss-2018b-Python-3.6.6'
        get_version: '--version'
        exit_code: 0

    junction_saturation.py:
        path: 'junction_saturation.py'
        module_load: 'lmod/lmod/libexec/lmod python load RSeQC/3.0.0-foss-2018b-Python-3.6.6'
        module_unload: 'lmod/lmod/libexec/lmod python unload RSeQC/3.0.0-foss-2018b-Python-3.6.6'
        get_version: '--version'
        exit_code: 0

    read_duplication.py:
        path: 'read_duplication.py'
        module_load: 'lmod/lmod/libexec/lmod python load RSeQC/3.0.0-foss-2018b-Python-3.6.6'
        module_unload: 'lmod/lmod/libexec/lmod python unload RSeQC/3.0.0-foss-2018b-Python-3.6.6'
        get_version: '--version'
        exit_code: 0

    read_GC.py:
        path: 'read_GC.py'
        module_load: 'lmod/lmod/libexec/lmod python load RSeQC/3.0.0-foss-2018b-Python-3.6.6'
        module_unload: 'lmod/lmod/libexec/lmod python unload RSeQC/3.0.0-foss-2018b-Python-3.6.6'
        get_version: '--version'
        exit_code: 0

# >> RSEQC END

    picard-tools:
        path: ['java', '-jar', 'software/picard/2.18.29-Java-1.8/picard.jar']
        module_load: 'lmod/lmod/libexec/lmod python load picard/2.18.29-Java-1.8'
        module_unload: 'lmod/lmod/libexec/lmod python unload picard/2.18.29-Java-1.8'
        get_version: '-h'
        exit_code: 1

    preseq:                                                                                                                                                                                                                               
        path: 'preseq'
        module_load: 'lmod/lmod/libexec/lmod python load preseq/2.0.3-foss-2018b'
        module_unload: 'lmod/lmod/libexec/lmod python unload preseq/2.0.3-foss-2018b'
        get_version: '--version'
        exit_code: 0

    samtools:
        path: 'samtools'
        module_load: 'lmod/lmod/libexec/lmod python load SAMtools/1.9-foss-2018b'
        module_unload: 'lmod/lmod/libexec/lmod python unload SAMtools/1.9-foss-2018b'
        get_version: '--version'
        exit_code: 0

## >>>>>>>>>>>>>>>>>>>>>>>>>>>>>>>>>>>>>>>>>>>>>>>>>>>>>>>>>>>>>>>>>>>>>>>>>>>>>>>>>
## CUSTOM AND MODIFIED TOOLS
## >>>>>>>>>>>>>>>>>>>>>>>>>>>>>>>>>>>>>>>>>>>>>>>>>>>>>>>>>>>>>>>>>>>>>>>>>>>>>>>>>
    collect_scs:
        path: *scs
        get_version: '--version'
        exit_code: 0

## >>>>>>>>>>>>>>>>>>>>>>>>>>>>>>>>>>>>>>>>>>>>>>>>>>>>>>>>>>>>>>>>>>>>>>>>>>>>>>>>>
## Unix Tools #
## >>>>>>>>>>>>>>>>>>>>>>>>>>>>>>>>>>>>>>>>>>>>>>>>>>>>>>>>>>>>>>>>>>>>>>>>>>>>>>>>>

    # TODO: check tools, remove not required tools

    awk:
        path: 'awk'
        get_version: '--version'

    cat:
        path: 'cat'
        get_version: '--version'
        exit_code: 0

    cp:
        path: 'cp'
        get_version: '--version'
        exit_code: 0

    cut:
        path: 'cut'
        get_version: '--version'
        exit_code: 0

    dd:
        path: 'dd'
        get_version: '--version'
        exit_code: 0

    echo:
        path: '/bin/echo'
        get_version: '--version'
        exit_code: 0

    grep:
        path: 'grep'
        get_version: '--version'
        exit_code: 0

    ln:
        path: ln
        get_version: '--version'
        exit_code: 0

    mkdir:
        path: 'mkdir'
        get_version: '--version'
        exit_code: 0

    mv:
        path: 'mv'
        get_version: '--version'
        exit_code: 0

    mkfifo:
        path: 'mkfifo'
        get_version: '--version'
        exit_code: 0

    # URL: http://zlib.net/pigz/
    pigz: 
        path: 'pigz'
        get_version: '--version'
        exit_code: 0

    pwd:
        path: 'pwd'
        get_version: '--version'
        exit_code: 0

    rm:
        path: 'rm'
        get_version: '--version'
        exit_code: 0      

    sort:
        path: 'sort'
        get_version: '--version'
        exit_code: 0

    tar:
        path: 'tar'
        get_version: '--version'
        exit_code: 0

    uniq:
        path: 'uniq'
        get_version: '--version'
        exit_code: 0
```
